# Stem-loop induced ribosome queuing in the uORF2/*ATF4* overlap fine-tunes stress-induced human *ATF4* translational control

**DOI:** 10.1101/2023.07.12.548609

**Authors:** Anna M. Smirnova, Vladislava Hronová, Mahabub Pasha Mohammad, Anna Herrmannová, Stanislava Gunišová, Denisa Petráčková, Petr Halada, Štěpán Coufal, Michał Świrski, Justin Rendleman, Kristína Jendruchová, Maria Hatzoglou, Petra Beznosková, Christine Vogel, Leoš Shivaya Valášek

## Abstract

ATF4 is a master transcriptional regulator of the integrated stress response leading cells towards adaptation or death. ATF4’s induction under stress was thought to be mostly due to delayed translation reinitiation, where the reinitiation-permissive uORF1 plays a key role. Accumulating evidence challenging this mechanism as the sole source of *ATF4* translation control prompted us to investigate additional regulatory routes. We identified a highly conserved stem-loop in the uORF2/*ATF4* overlap, immediately preceded by a near-cognate CUG, which introduces another layer of regulation in the form of ribosome queuing. These elements explain how the inhibitory uORF2 can be translated under stress, confirming prior observations, but contradicting the original regulatory model. We also identified two highly conserved, potentially modified adenines performing antagonistic roles. Finally, we demonstrate that the canonical ATF4 translation start site is substantially leaky-scanned. Thus, *ATF4’s* translational control is more complex than originally described underpinning its key role in diverse biological processes.

## INTRODUCTION

Eukaryotic cells have evolved several mechanisms to cope with various environmental stressors. These include a complex signaling pathway referred to as the Integrated Stress Response (ISR) (Pakos-Zebrucka et al., 2016; Wek et al., 2006). Whereas its external triggers are oxygen, nutrient deprivation or viral infections, the main internal stressor is the accumulation of unfolded proteins in the lumen of the endoplasmic reticulum (ER). Importantly, the ISR can also be induced by activation of oncogenes (Denoyelle et al., 2006). The response is “integrated” as all stress signals are transduced by a family of four serine/threonine kinases and converge into a single event which is the phosphorylation of the α subunit of eukaryotic translation initiation factor 2 (eIF2α) (Donnelly et al., 2013).

eIF2 assembles into a ternary complex (TC) together with GTP and the initiator Met-tRNA_i_^Met^. This complex, together with other eIFs, facilitates the Met-tRNA_i_^Met^ recruitment to the ribosomal P-site so that the pre-initiation complex (PIC) can recognize the translation start site and initiate translation (Valášek, 2012). After the AUG selection process, eIF2 with hydrolyzed GDP is ejected from the initiation complex. To participate in the next translational cycle, it must be regenerated by the eIF2B-mediated exchange of GDP for GTP (Hinnebusch, 2014). The ISR-transduced phosphorylation of the α subunit of eIF2 prevents this regeneration step, leading to a significant decrease in TC levels and a general translational shutdown (reviewed in (Dever et al., 2023; Gunisova et al., 2018)).

A handful of specific mRNAs escape this shutdown and become efficiently translated to begin adaptation to the acute stress (Harding et al., 2000). One of them encodes the transcription factor *ATF4,* a master regulator of the ISR (Lu et al., 2004; Vattem and Wek, 2004). Upon relief from the acute stress, eIF2α is dephosphorylated, the ISR ceases, and general protein synthesis resumes (Novoa et al., 2003). However, if the stress is too intense or persists for too long, it becomes chronic (Guan et al., 2017), the adaptive response capacity is exhausted and programmed cell death is triggered. Thus, the final outcome of the ISR depends largely on the level and duration of eIF2α phosphorylation and, consequently, on the extent of upregulation of factors escaping the shutdown (Pakos-Zebrucka et al., 2016).

Therefore, the timing and level of ATF4 expression are critical parameters determining cell fate. Even minor disturbances in the ATF4 function contribute to serious pathologies such as Parkinson’s, Alzheimer’s and Huntington’s neurodegenerative diseases, prion diseases and various types of retinal degeneration (Pitale et al., 2017). Furthermore, persistent over-activation of ATF4, which does not induce apoptosis, is linked to many cancers because of the continuous expression of adaptive genes that sustain the stress response (Wortel et al., 2017).

Under normal conditions, human *ATF4* is constitutively transcribed at low levels, but the protein is undetectable. Induction of its expression is mediated by the *ATF4* 5’ mRNA leader containing two upstream open reading frames (uORFs): a short 4-codon uORF1 followed by a longer 60-codon uORF2, which overlaps with the *ATF4* reading frame in a −1 frame. In analogy with its extensively studied yeast functional homologue *GCN4* (Dever et al., 2023; Gunisova et al., 2018; Gunisova and Valasek, 2014; Hinnebusch, 2005), it has been proposed that the primary route of *ATF4* translational control occurs via the so-called delayed translational reinitiation (REI) (Lu et al., 2004; Vattem and Wek, 2004). This model is underpinned by the stress-induced decrease in TC levels and the opposing properties of the two *ATF4* uORFs (Fig. 1A) (Dever et al., 2023; Gunisova et al., 2018).

**Figure 1.**
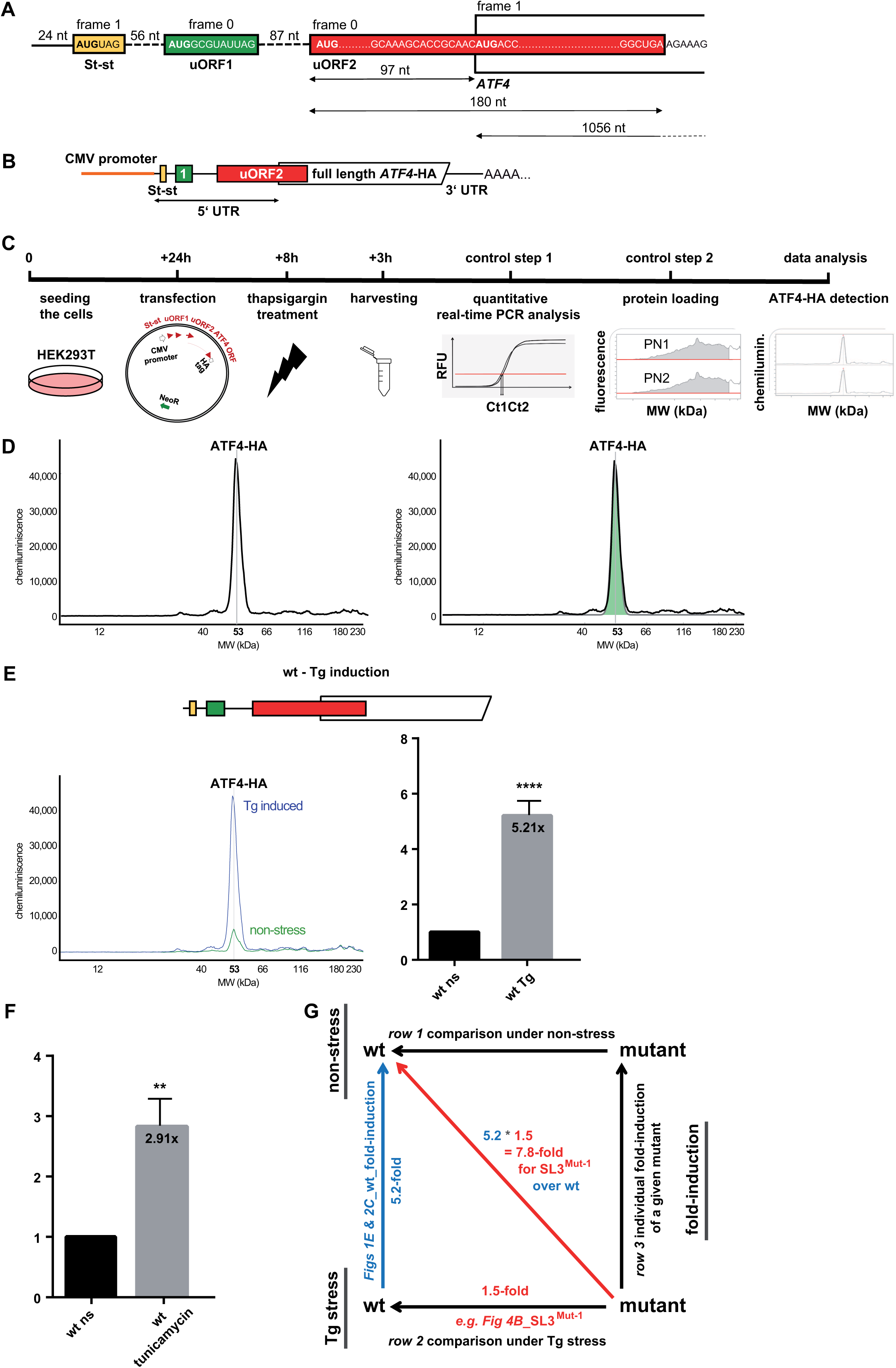
Revisiting translational control of human *ATF4* – reporters and experimental set-up. (A) Schematic showing the 5’ end of the mRNA encoding the human activating transcription factor 4 (*ATF4*; NM_182810.2, transcript variant 2), featuring the color-coded Start-stop (St-st) element (yellow), REI-permissive (green) 3-codon uORF1, and inhibitory uORF2 (red) overlapping with the beginning of main *ATF4* ORF. Frames and distances are given in nucleotides (nt). (B) Schematic of the wt CMV-driven ATF4-HA tagged construct; the HA tag was placed immediately upstream of the *ATF4* stop codon. (C) Experimental workflow described in the form of a time-line diagram; PN - JESS Protein Normalization Detection Module. (D) All HEK293T cell lysates were subjected to protein separation and immunodetection using the JESS system. The signal, detected in the capillary is represented as electropherogram (a single peak of the ATF4-HA tag full-length protein size of 53 kDa, left panel) and automatically quantified (right panel); expression of wt construct under 3h Tg stress conditions 8 h post-transfection detected by anti-mouse HA tag antibodies is shown. (E) Stress-induced upregulation of the ATF4-HA protein expression under Tg stress (in blue) compared to non-stress conditions (in green) is shown. Quantified “fold-induction” data were plotted (right panel). The differences between experimental groups were tested by the t-test. Variables are presented as mean ± SD and *p* values of < 0.05 were considered statistically significant (i.e., p<0.05 *, p<0.01 **, p<0.001 ***, p< 0.0001 ****). (F) Stress-induced upregulation of the ATF4-HA protein expression from the wt reporter under tunicamycin stress compared to non-stress conditions (set to 1). Quantified “fold-induction” data were plotted and analyzed as in panel E. (G) Experimental set-up illustrating determination of fold-change values when comparing: i) each mutant reporter construct *versus* the wt construct separately under non-stress (upper horizontal arrow) and Tg stress conditions (lower horizontal arrow), and ii) the fold-induction expression of a mutant under stress with the same mutant under non-stress (right vertical arrow), or wt under stress *vs*. no stress (blue left vertical arrow). The red diagonal arrow indicates calculated fold-induction changes of a given mutant under stress *versus* wt under non-stress conditions; i.e. how much each mutant increases or decreases the ~5.2-fold induction of the wt reporter.

Specifically, after translation of the REI-permissive uORF1, only the large ribosomal subunit is recycled, while the small subunit remains associated with the mRNA and, upon regaining scanning ability, can reinitiate downstream. This process is enabled by the short length of uORF1 and the presence of stimulatory flanking sequences interacting with the initiation factor eIF3 (Hronova et al., 2017). This interaction allows eIF3 to remain transiently associated with elongating and terminating ribosomes on short uORFs to allow REI (Mohammad et al., 2017; Mohammad et al., 2021; Wagner et al., 2020).

A key aspect of this delayed REI mechanism is that reacquisition of the scanning competence of the post-termination 40S is limited by its binding of a new TC, and that the time required for TC acquisition increases significantly with decreasing TC levels (Lu et al., 2004; Vattem and Wek, 2004). The original model posits that, in the absence of stress, the 40S acquires the TC relatively quickly, recognizes the uORF2 AUG, and due to its overlap with *ATF4*, no ATF4 protein is produced. Conversely, the stress-induced decrease in TC levels delays the TC acquisition by the majority of 40Ses traversing downstream of uORF1. This allows them to skip the AUG of uORF2 and acquire the TC while *en route* to the *ATF4* start codon to induce its synthesis.

Despite the depth of the original studies (Lu et al., 2004; Vattem and Wek, 2004), a growing body of evidence suggests the existence of additional regulatory modes beyond the delayed REI mechanism. For example, three independent studies have reported substantial translation of uORF2 even under stress (Andreev et al., 2015; Sidrauski et al., 2015; Starck et al., 2016). In addition, leaky scanning over uORF2 was shown to contribute to *ATF4* induction (Starck et al., 2016). It was proposed that a N^6^-methyladenosine (m^6^A) post-transcriptional modification in the non-overlapping region of uORF2 of mouse *ATF4* mRNA functions as a barrier to ribosomes scanning downstream of uORF1 and that stress triggers demethylation of m^6^A removing this block (Zhou et al., 2018). QTI-seq performed in the same study also implicitly questioned the true initiation start site of the *ATF4* coding sequence, which begins with three nearly consecutive AUGs. (Note that the *ATF4* transcript in mouse is largely conserved with the human transcript (also see Fig. S1A) and was examined in the original studies (Vattem and Wek, 2004).) Employing the Sel-TCP-Seq technique (Wagner et al., 2022), we showed that the 5’ most proximal and so far the least studied uORF0 (hereafter referred to as Start-stop [St-st] because it consists only of an initiation and termination codon), which precedes uORF1 (Fig. 1A), represents an additional barrier that may inhibit ATF4 expression in the absence of stress (Wagner et al., 2020). All these findings suggest more complex regulatory system exceeding the relatively straightforward mode of delayed REI, whose full understanding may help to explain all aspects of the *ATF4* contribution to ISR as its master regulator.

Here, we subjected the human *ATF4* mRNA leader to complex bioinformatics analysis and uncovered several novel *cis*-acting features. Following detailed mutational analysis using our new and well-controlled reporter assay revealed that: 1) a highly conserved stem-loop in the uORF2/*ATF4* overlap causes ribosome queuing and stalled or slowed-down ribosomes may initiate at a near-cognate CUG codon upstream of this stem-loop; 2) the inhibitory uORF2 has an unexpected stress-inducible character and is relatively highly translated even under stress, which could be explained by the ribosome queuing; 3) the first two AUGs of ATF4 are substantially leaky-scanned; 4) two prospective, highly conserved adenine modification sites contribute to overall regulation in opposing ways, one of them antagonizing the role of the stem-loop. Overall, our data suggest that translational control of ATF4 comprises a multilayered regulatory circuit of diverse sequence and structural elements that fine-tune its expression under different stimuli.

## RESULTS

### Revisiting translational control of human *ATF4*

Published analyses of ATF4’s translational control were carried out using a mouse *ATF4* mRNA reporter (mouse and possum *ATF4* lack St-st, which is otherwise highly conserved among vertebrates (Fig. S1A)) (Rendleman et al., 2023), and *Luciferase* was fused with *ATF4* only two codons downstream of its AUG (Vattem and Wek, 2004). The wild type (wt) and specific mutation(s)-carrying reporters were individually co-transfected with a control *Renilla* plasmid into MEF cells and Luciferase activity was measured. We perceived two potential bottlenecks in this arrangement. 1) In addition to lacking St-st, the original ATF4 reporter also lacked much of the uORF2/*ATF4* overlap (as well as the entire ATF4 ORF), which limited investigation of its role. 2) Its activity was not normalized to *ATF4* mRNA levels which are known to change (Dey et al., 2010)) but to Renilla activity, which by definition drops rapidly upon stress-induced translational shutdown. To resolve these discrepancies, we designed an entirely new ATF4-based reporter, this time using the human sequence, and an experimental workflow that avoided normalizing ATF4 expression levels to any reference genes whose expression would be translationally shut down under stress conditions.

Our newly designed CMV promoter-driven reporter preserves the entire V2 transcript leader sequence and the entire coding region of *ATF4*, which is extended with an HA tag at its C-terminus (Fig. 1B). Twenty-four hours after seeding, HEK293T cells were individually transfected with the same amount of a control wt reporter or its mutant derivatives and after 8 hours, one half of the cultures was treated with thapsigargin (Tg; induces unfolded protein response - UPR), the other half with DMSO (control)(Fig. 1C). Therefore, the wt control was used as a reference in every single experiment. Exactly 3 hours after Tg treatment, cells were harvested using a standard Glo lysis buffer supplemented with protease inhibitors by the vendor (Promega) and the lysates were subjected to a fully quantitative, automated, and capillary-based high throughput immunoassay (“western blotting”) using JESS Protein Simple instrument, as described in detail in the Methods (Fig. 1C).

Figure 1D shows an example of the output data in the form of an electropherogram, which displays the intensity detected along the length of the capillaries; the generated peaks can be quantified by calculating the area under the curve. In case of ATF4-HA, the highly specific signal detected by the anti-HA tag antibodies is displayed as a single peak at 53 kDa and shows ~5.2-fold induction (*p*<0.0001) upon Tg treatment 8-hours post-transfection (Fig. 1E, see Table S1 for average values of fold-induction for wt construct and Table S2 for individual values from each experiment). Tunicamycin-stressed cells (treated the same way only exposed to stress for four hours) generated nearly 3-fold induction (Fig. 1F, Table S1 and Table S2). For statistical analysis, see Methods. Please note that we routinely recorded a slightly increased activity of our reporter even under non-stress conditions (Fig. 1E, green peak), the source of which is explained in Supplementary Information (SI). This feature of our reporter system proved to be very useful throughout the study because it served as a “background” value used to compare the effects of the mutations on both basal ATF4 expression and its inducibility under Tg stress; this would have been impossible with a background value of zero. For experiments validating that our new reporter system faithfully mimics endogenous *ATF4* regulation, please see SI and Figs. S1B – D and S2A.

### St-st modestly inhibits ATF4 expression, uORF1 allows for 50% downstream REI, and a solitary, inhibitory uORF2 allows for high ATF4 stress-inducibility

We began our analysis by testing the effect of the St-st and re-testing the effects of uORFs (on their own) on ATF4 expression by mutating the AUGs of the respective other two elements in the otherwise wt construct (Fig. S3A and B). First, we measured the control construct (d-all), where AUGs of all three elements were mutated. This construct determined the maximal expression level of ATF4, which was ~22.7-fold and ~3.3-fold higher in “non-stress” and “Tg stress” conditions, respectively, compared to the values of the wt construct that were set to 1 for each of these two conditions individually (Fig. S3A and B; Tables S3 and S4). The last plot in the d-all column (“fold-induction”) with the d-all non-stress value set to 1 indicates that this construct is expectedly not stress-inducible, in contrast to wt (Figs. 1E and S3C).

Note that all “fold-induction” experiments throughout the study, e.g. in row 3 of Figure S3A, were conducted separately from the “non-stress” and “Tg stress” experiments (e.g. rows 1 and 2, Fig. S3A). To further clarify our set-up, in the “non-stress” and “Tg stress” experiments, we compared individual mutants with wt separately in the respective conditions, whereas in the “fold-induction” experiments, we compared expression of a given mutant under stress with that of the same mutant under no stress (DMSO) (Fig. 1G). This means that both sets of experiments were individually controlled for the equal amount of reporter mRNA and protein levels, as described above and SI.

Our results, presented in detail in SI and Figure S3, confirmed major aspects of the original model and supported the idea that St-st acts as a general repressive element, i.e., a roadblock, as observed before (Rendleman et al., 2023; Wagner et al., 2020). Unexpectedly, they also revealed a stress induction capability of the uORF2-only reporter of an unknown mechanism (see below).

### Sequence analysis of the 5’ UTR of the human *ATF4* mRNA reveals additional elements potentially contributing to *ATF4* translational control

Next, we carefully screened the *ATF4*’s 5’ UTR and the uORF2/*ATF4* overlap in the coding region for sequence and structure features that might suggest additional modes of regulation (Fig. 2A). Their detailed description is provided in SI. The most notable element, predicted by our analysis, is a stem-loop (SL3), with ΔG = −15.40 kcal/mol, which is highly conserved among vertebrates (Fig. S1A) and could be inhibitory. It is located roughly in the middle of the uORF2/*ATF4* overlap (Fig. 2A) and has a hairpin structure with the highest free energy predicted for this particular sequence region. Our analysis also predicted a total of four potential sites of m^6^A methylation within well-defined motifs (Fig. 2A). Interestingly, the *ATF4* gene begins with two nearly consecutive AUGs followed by a third in proximity, all in frame (Fig. 2A, in dark green). Both the canonical AUG1 and AUG2 have a medium Kozak initiation context; AUG1 overlaps with DRACH2, and AUG2 is located two codons downstream. AUG3 has a weak Kozak initiation context and is exposed in the open loop of SL3. It represents the 17^th^ codon downstream of AUG1. Both AUG2 and AUG3 are highly conserved among vertebrates (Fig. S1A).

**Figure 2.**
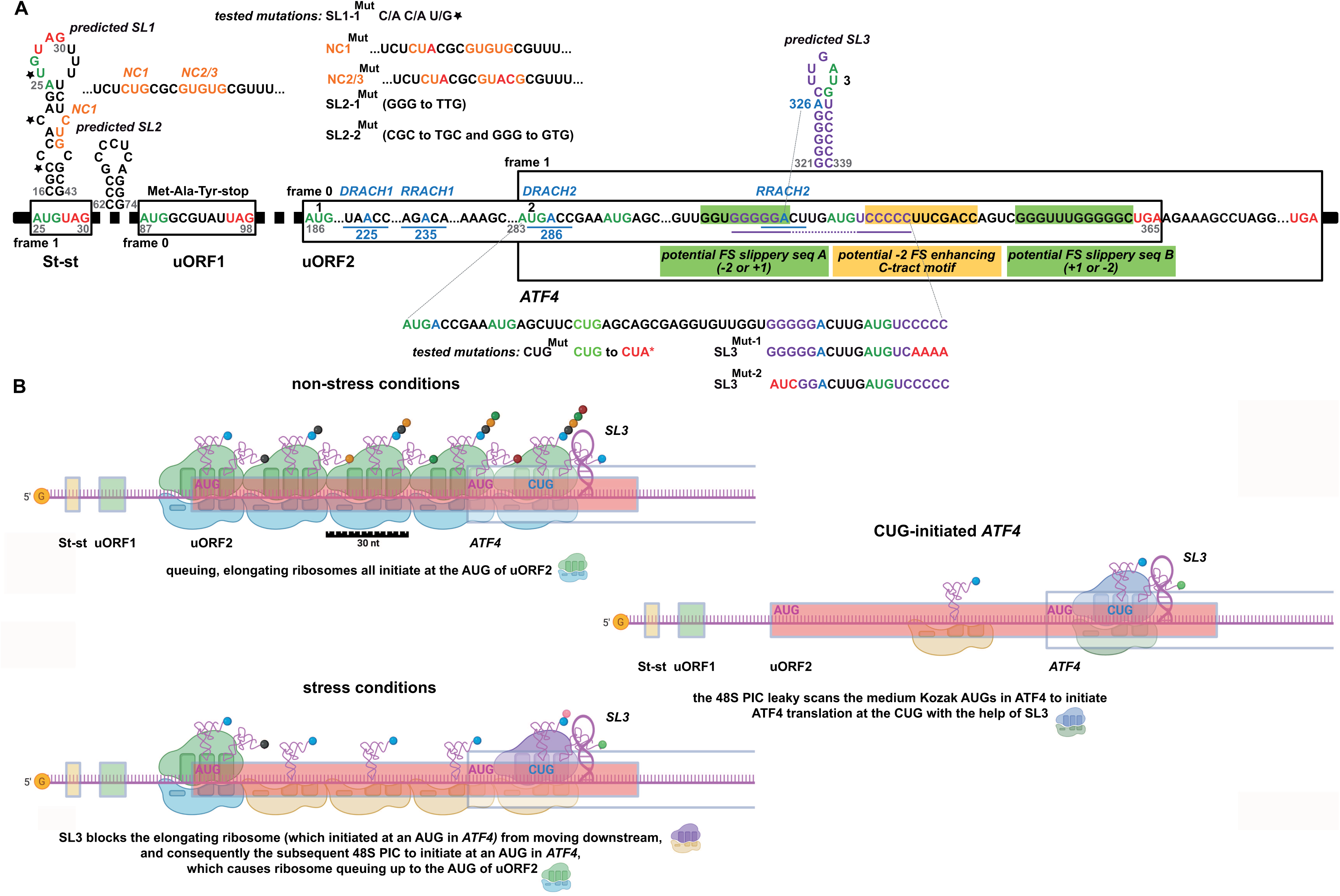
Sequence analysis of the 5’ UTR of the human *ATF4* mRNA; ribosome queuing model expanding the mode of ATF4 translational control. (A) Schematic of novel, bioinformatically predicted, potential regulatory features within the 5’ UTR of the *ATF4* mRNA and beginning of the *ATF4* main ORF; point mutations are depicted. For details, please see the main text. (B) Model of the ribosome queuing mechanism under non-stress *versus* stress condition employing SL3 and near-cognate CUG as an additional layer of *ATF4* translational control. For details, please see the main text. Created with BioRender.com.

Given this knowledge, we created an array of mutants for all these predicted elements (Fig. 2A) and first evaluated them in a large-scale screen. Subsequently, we focused our analysis on mutations in elements that had observable effects like SL3.

### SL3 delays the flow of ribosomes in the uORF2/*ATF4* overlap and genetically interacts with the upstream near-cognate CUG codon

Next, we unfolded SL3 by either quadruple C to A (in SL3^Mut-1^) or GGG to AUC (in SL3^Mut-2^) substitutions (Fig. 2A) and observed ATF4 expression to increase by ~1.3- and ~1.7-fold under non-stress, and by ~1.5- and ~1.8-fold under Tg stress, respectively, over the wt construct set to 1 (Fig. 3B – rows 1 and 2; Table S5 and Table S6). This means that SL3 mutations add a further ~50-80% increase (Fig. 3B – row 2) to the ~5.2-fold induction of the wt reporter (Fig. 3A) to reach the robust ~7.8-9.4-fold induction (5.2 multiplied by 1.5 or 1.8) under stress conditions. Figure 1G illustrates how these fold-induction changes were calculated. Accordingly, loosening SL3 also increased inducibility of both these mutant constructs measured and compared separately under stress versus non-stress conditions from ~5.2-fold to ~5.7-5.9-fold (Fig. 3A *versus* Fig. 3B). These results suggest that SL3 forms and acts as physical barrier delaying scanning/translating ribosomes under both conditions.

**Figure 3.**
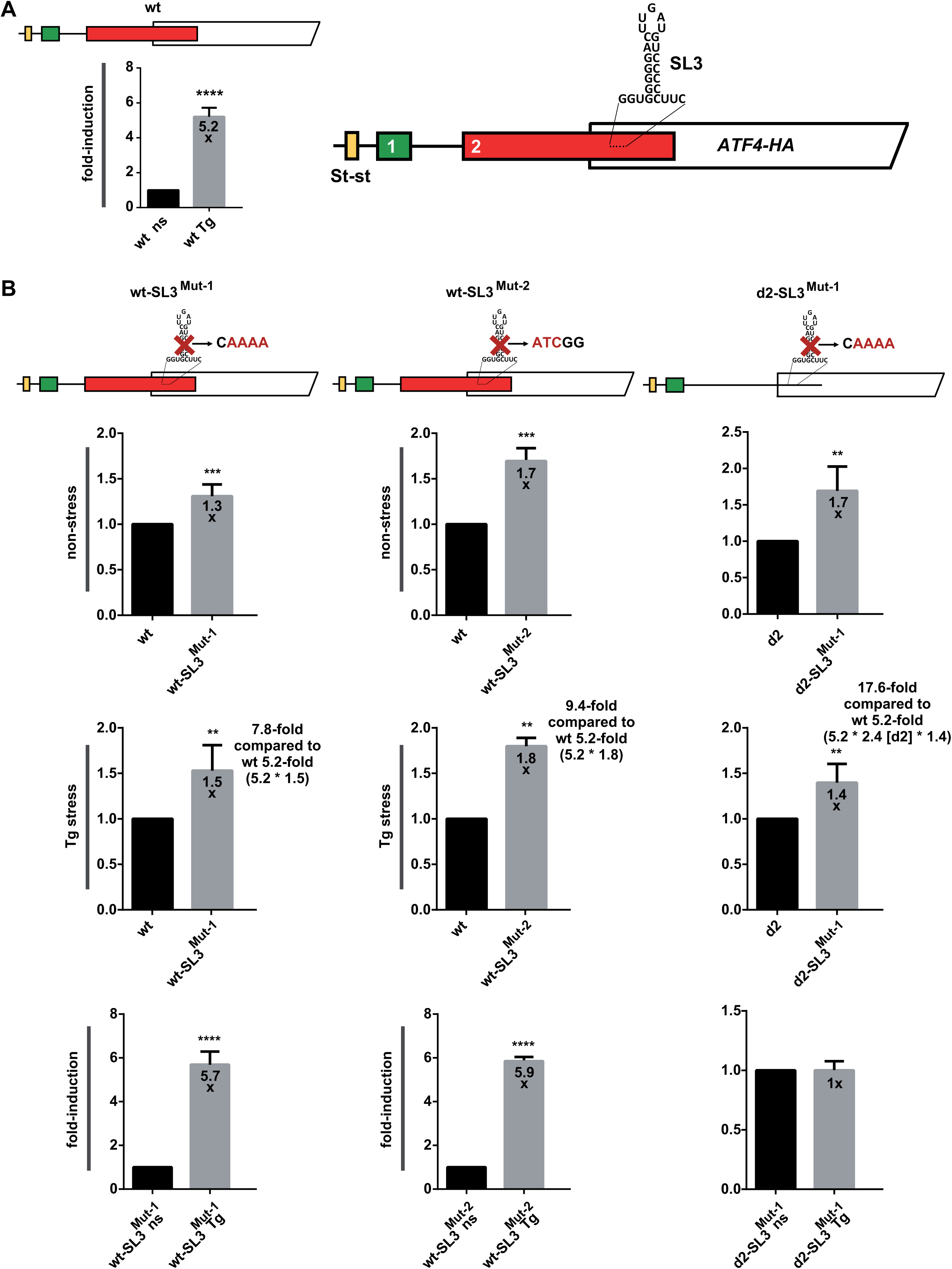
SL3 delays the flow of ribosomes in the uORF2/*ATF4* overlap. (A) Same as Figure 1E for better comparison. (B) Same as in Figure 2A except that the SL3 *ATF4* mutant constructs depicted at the top of the corresponding panels were subjected to JESS analyses. The differences between experimental groups were tested by the t-test except wt-SL3^Mut-1^ Tg stress and d2-SL3^Mut-1^ non-stress, where Mann-Whitney test was used.

Eliminating SL3 in the d2 construct (in d2-SL3^Mut-1^) showed a similar increase over the d2 construct alone as in case of the wt-SL3^Mut-1^ mutant over wt (Fig. 3B – rows 1 and 2; Table S5 and Table S6). Note that the overall induction (d2-SL3^Mut-1^ over wt) under Tg stress was robustly increased by ~17.5-fold (Fig. 3B – row 2). These results strongly suggest that both uORF2 and SL3 act together to reduce ATF4 translation.

Initiating 48S PIC with the start site placed in the P site covers ~12 nt from the mRNA entry site up to the P site (excluding it) and ~15-25 nt downstream from the P-site (including it) to the mRNA exit site (Fig. 2B) (Wagner et al., 2022; Wagner et al., 2020). We noticed a near cognate CUG codon in weak Kozak context positioned 20 nucleotides upstream of SL3 (Fig. 2A), which is highly conserved among vertebrates (Fig. S1A). This position is ideal to force a scanning 48S PIC stalled at or slowed by SL3 to initiate at CUG (Fig. 2B). To test this hypothesis, we replaced CUG with CUA (in CUG^Mut^) and observed that the mutation significantly reduced ATF4 expression by ~20% under both conditions tested, which corresponds to 4.2-fold reduction (5.2 of the wt construct multiplied by 0.8) of “fold-induction” under stress (Fig. 4B, middle panel – row 2; Table S7 and Table S8).

**Figure 4.**
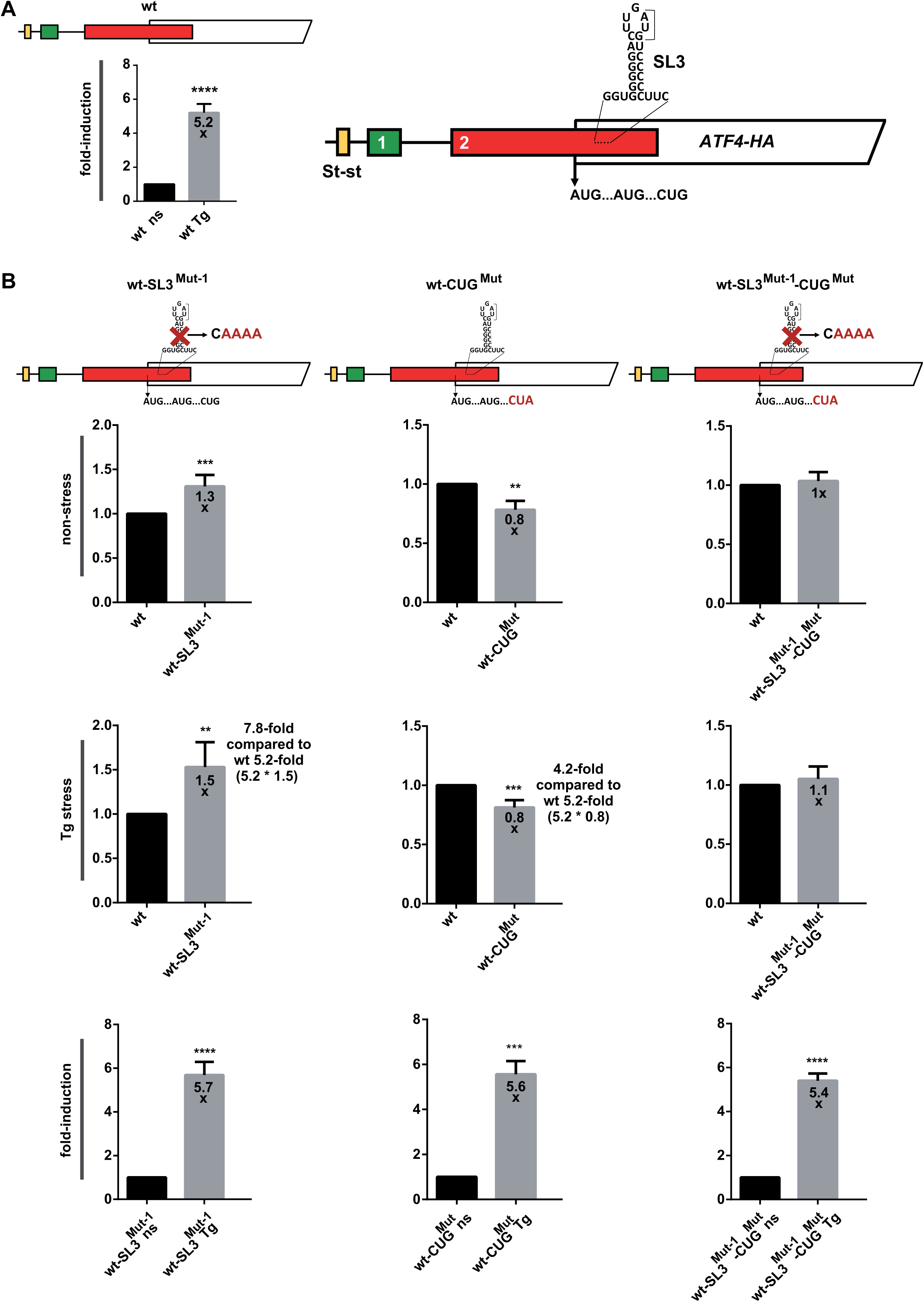
SL3 genetically interacts with the upstream CUG near-cognate codon. (A) Same as Figure 1E for better comparison. (B) Same as in Figure 2A except that the CUG to CUA and SL3^Mut-1^ *ATF4* mutant constructs depicted at the top of the corresponding panels were subjected to JESS analyses.

These findings suggest that a small proportion of the ATF4 protein might be several amino acids shorter due to this CUG-initiated synthesis downstream of the main initiation site. Interestingly, combining CUG and SL3 mutations in a single SL3^Mut-1^-CUG^Mut^ construct nullified the opposing effects of the two individual mutations, suggesting a functional interaction between the two elements (Fig. 4B, right panel; Table S7 and Table S8). Specifically, when the translation inhibitory SL3 was removed, the translation enhancing CUG was no longer needed, perhaps because a flux of ribosomes initiating at the *ATF4* was released to maximize the synthesis of the full-length ATF4 protein. It remains to be investigated whether or not the CUG-initiated ATF4 protein is stably produced in wt cells.

To further investigate the functional interplay between SL3 and CUG, we analyzed three publicly available ribo-seq libraries generated from cells exposed to various ER stressors. Namely, two sets of HEK293T cells treated with arsenite (Andreev et al., 2015; Ichihara et al., 2021) and HeLa cells subjected to the tunicamycin-induced stress (Rendleman et al., 2018). We compared the total number of footprints (FP) in the *ATF4* mRNA leader in stressed *versus* non-stressed cells (Fig. S4), as well as the distribution of FPs with ribosomes positioned with their P-site to a specific codon (Fig. S5). In addition to the pile-ups of FPs on the St-st and AUGs of uORF1 and uORF2 (the latter two are observed in only two datasets - Fig. S4A - D), all three datasets clearly revealed a pile-up of FPs ~20 nucleotides upstream of SL3, with a visible reduction under stress (Fig. S5). In contrast, there were virtually no FPs on the AUG1 of *ATF4*, even under stress, while the remainder of the coding region was correctly stress-induced in all datasets. A closer look at the FPs indicated a strong preference of the P-site locating to the CUG in all three datasets (Fig. S5). We argue that even though the FP coverage in the uORF2/*ATF4* overlap upstream of SL3 was relatively low, the repeated occurrence of this prominent “CUG” peak in 3 independent datasets did not occur by chance. Therefore, the existence and functionality of SL3 and its associated near-cognate CUG are a distinct possibility.

### Ribosome queuing contributes to the overall translational control of *ATF4*

The existence of SL3 and its interplay with the upstream CUG intrigued us because it could trigger a) uORF2-to-*ATF4* frameshifting (see SI) or b) ribosome queuing, i.e., the phenomenon implicated for example in translational control of the antizyme inhibitor mRNA (Ivanov et al., 2018). Specifically, under non-stress conditions, SL3 could form a queue of 80S ribosomes elongating from the AUG of uORF2, reducing its translation rate and thus completely eliminating any leaky scanning. This would in turn result in even tighter suppression of ATF4 translation under normal conditions. Indeed, based on an 80S ribosome’s average FP length of 30 nucleotides (Archer et al., 2016; Ingolia, 2010), altogether five 80S ribosomes could be accommodated in this queue, with the most 5’ ribosome positioned with its P-site on the AUG of uORF2. The ribosome queue would not be resolved until the barrier posed by SL3 was breached (Fig. 2B).

According to this model, under stress, the first 80S ribosome initiating at the AUG of *ATF4* would soon be halted or at least slowed down by SL3, preventing the incoming 48S PIC from moving forward to reach the *ATF4*’s AUG due to a spatial constraint (Fig. 2B). This constraint would then be extended into a queue of 48S PICs that would span up to the AUG of uORF2. Consequently, the most 5’ 48S PIC could potentially initiate at uORF2, triggering its translation under stress, although the length of such a mixed queue of 48S PICs and 80S ribosomes is hard to predict due to variable FP length of scanning 48S PICs. In any case, this model could explain that, contrary to what the “delayed REI” implies, uORF2 is translated even under stress at substantial levels, which has been observed by us (see below) and others (Andreev et al., 2015; Sidrauski et al., 2015; Starck et al., 2016). Our model would also imply that, thanks to SL3, CUG could be utilized as an alternative start site for all 40S ribosomes that leaky scan the *ATF4*’s AUG (Fig. 2B); and below we demonstrate that this AUG is indeed rather “leaky”. Such an intricate system would ensure well-balanced induction of ATF4 which is highly desirable given its key role (Pitale et al., 2017).

To test the above model, we extended the coding sequence of *ATF4* by inserting 6x c-Myc tag (30 nts each) just after AUG1 (Fig. 5, Table S9 and Table S10); i.e., two codons upstream of AUG2 (Fig. 5A). If our logic was correct, this extension would increase ATF4 expression in the otherwise wt construct under both stress and non-stress conditions, while having no effect in the absence of SL3, because extending the spacing between SL3 and the AUGs of uORF2 and *ATF4* should eliminate SL3’s negative effect. This is exactly what we observed (Fig. 5B). Further supporting this model, unfolding SL3 in the absence of uORF2 (d2_ins-SL3^Mut-1^) did not eliminate the stimulatory effect of the 6x c-Myc tag extension (Fig. 5C and D). These results further underscore the observed interplay between uORF2 and SL3 by showing that the SL3 effect relies largely on initiation at the AUG of uORF2 (see also below).

**Figure 5.**
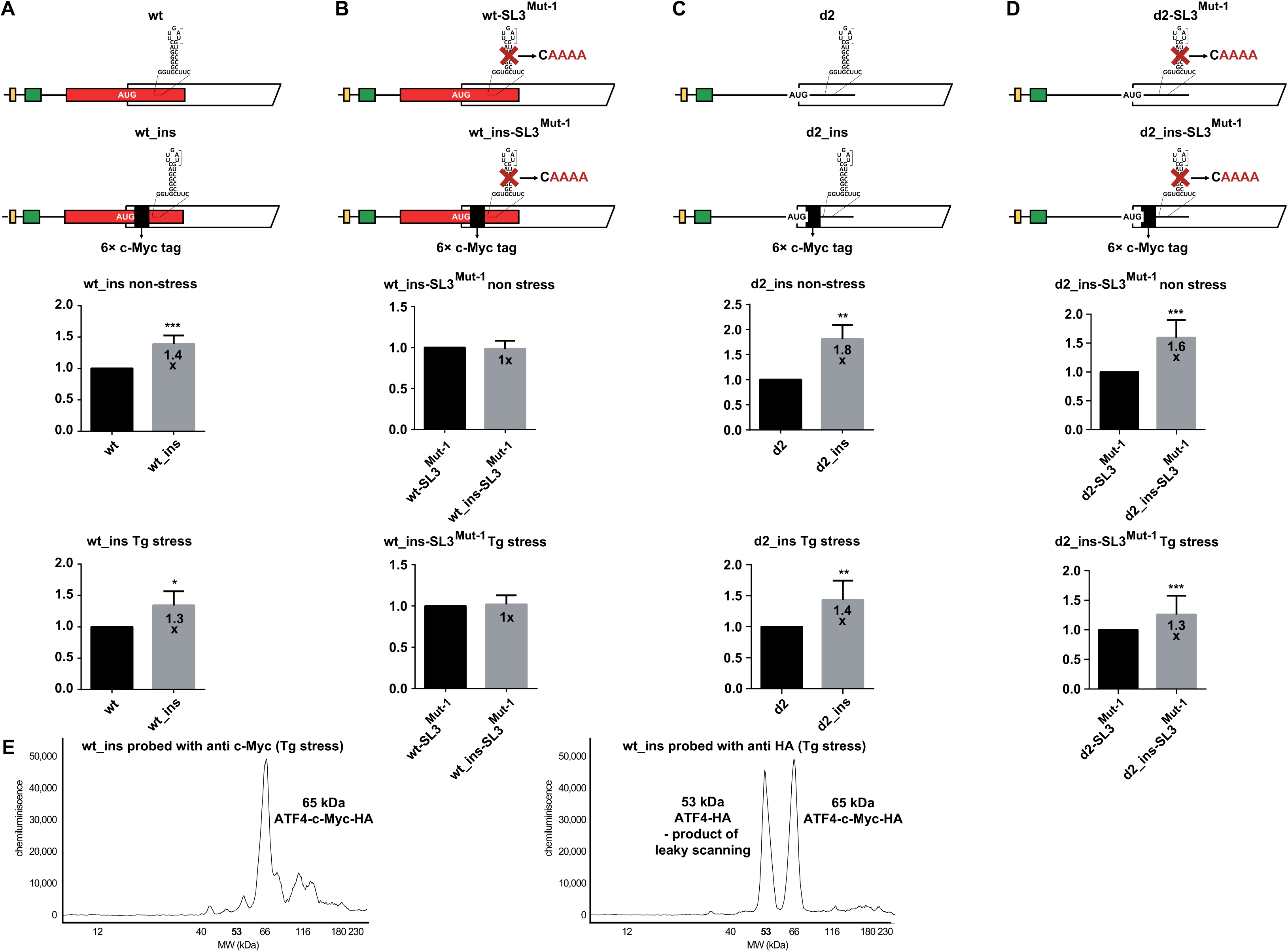
Ribosome queuing and a substantial leaky scanning at AUG1 of *ATF4* contributes to its overall translational control. (A – D) Same as in Figure 2A except that the 6x c-Myc tag insertion in-frame with *ATF4* (A), combined with the SL3^Mut-1^ mutation (B), in the otherwise wt construct (A – B) *versus* the construct lacking uORF2 (C – D), all depicted at the top of the corresponding panels, were subjected to JESS analyses. The differences between experimental groups were tested by the t-test except d2_ins_SL3^Mut-1^ Tg stress, where Mann-Whitney test was used. (E) The first AUG of the *ATF4* ORF is robustly leaky scanned. The electropherograms of the *ATF4* construct bearing the 6x c-Myc tag in-frame insertion probed with the anti-c-Myc (left panel) and anti-HA (right panel) antibodies are shown. For details, see the main text.

Importantly, while the c-myc detection revealed only a single peak of 65 kDa generated by the wt_ins construct, as expected given the 6x c-Myc tag insertion, the HA probing detected an additional peak just above, running at the size of the original ATF4 protein (53 kDa, Fig. 5E). This peak could only be explained by a shorter ATF4 variant(s) produced by initiation at AUG2 and/or the CUG and/or AUG3, in further support of substantial leaky scanning at AUG1, as inferred above.

To further support the ribosome queuing hypothesis, we treated the non-stressed and stressed HEK293T cells (without any reporter) with the cross-linking agent formaldehyde (HCHO) or control (non-crosslinking) cycloheximide, lysed the cells, applied RNase I, isolated total RNA and subjected the resulting RNA sample to RT-qPCR. For this ribosome-protection assay, three sets of primers were used: A1 (for Amplicon 1) covering almost the entire putative queuing region (132 nt of total ~150 nt, beginning with uORF2’s AUG and ending right in front of SL3); A2 of a similar length covering the region immediately downstream of SL3; and A3 of a similar length covering the region in the middle of the *ATF4* CDS (Fig. 6A). Primer sets A2 and A3 were used independently for normalization and were designed based on the publicly available riboseq data used in this study (Figs. S4, S5, and S6B) in the regions with detectable but low ribosome coverage. All 3 primer pairs had comparable efficiency ranging from 1.95 to 2.05 and generated fragments of the expected length upon RNase I treatment, as also verified by sequencing (data not shown). We reasoned that if ribosomes were queued, they should protect the putative queuing region (A1) from the RNase I digestion more so than the other two control regions (A2 and A3). Accordingly, we found a robust enrichment of Amplicon 1 over Amplicon 2 in stressed (~42-fold) and also to a lesser but still very high extent (~11-fold) in non-stressed cells in the RNase I-digested HCHO samples compared to undigested samples (Fig. 6B; for raw data see Supplementary Excel File 1). Consistently, the cycloheximide non-crosslinking control, which by definition should protect queued 80S ribosomes with lower efficiency, showed much smaller difference (~3-fold); moreover, regardless of stress (Fig. 6C). Importantly, given that only HCHO, but not cycloheximide, can protect 40S-bound mRNAs, the clear difference in fragment protection under stress *versus* non-stress conditions observed in RNase I-digested HCHO-crosslinked samples suggests the existence of a mixed queue of 48S PICs and 80S ribosomes under stress. Collectively, these data further support our model proposing the formation of the 80S queue under non-stress conditions and a mixed queue under stress conditions (Fig. 2B), the latter of which seems to be more prevalent (Fig. 6B). Note that similar results were also obtained when Amplicon 1 was compared with Amplicon 3 (Fig. S6A – B; for raw data see Supplementary Excel File 1).

**Figure 6.**
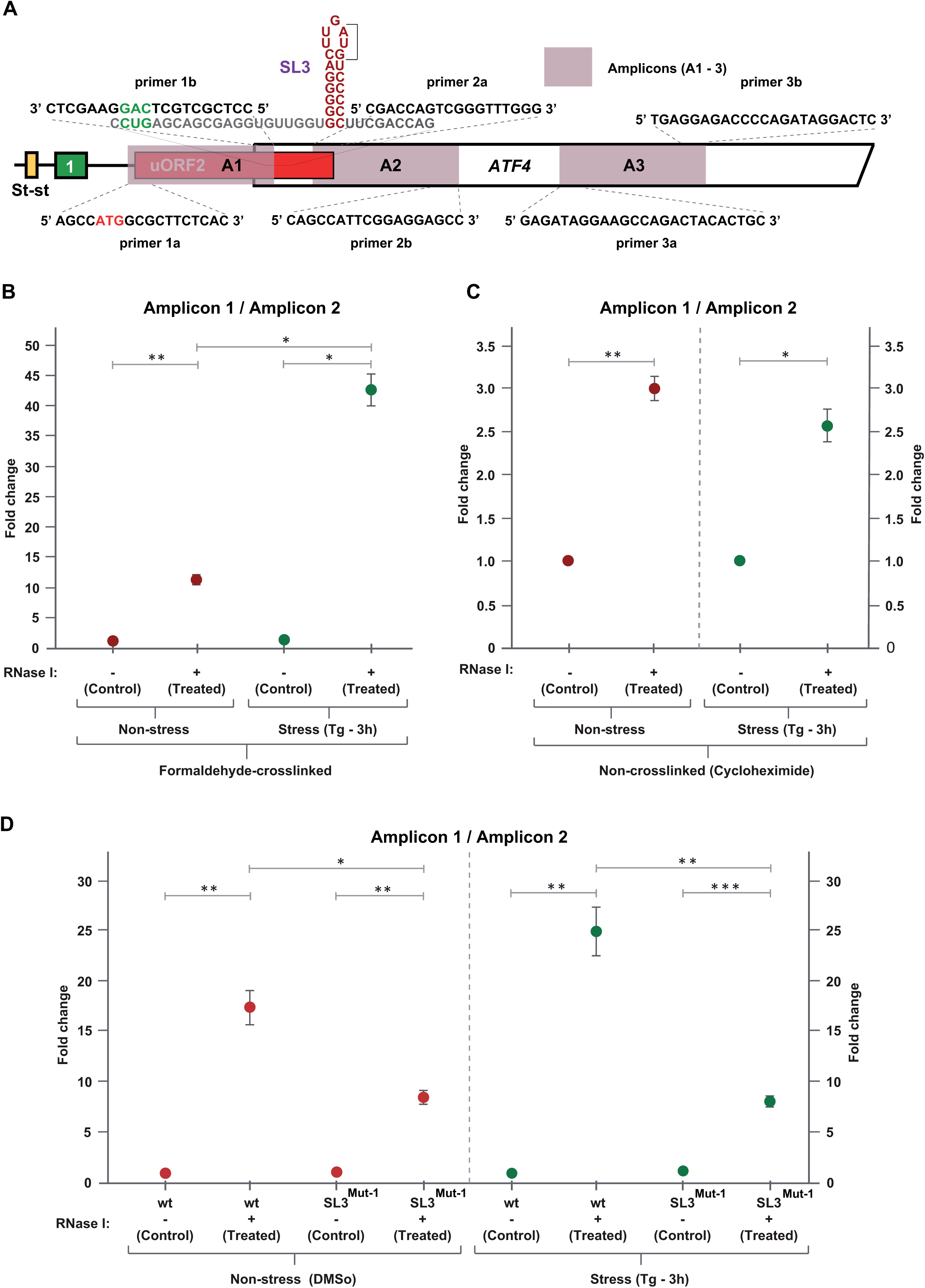
Ribosome-protection assay demonstrating that SL3 pauses ribosomes and prompts their queuing under both non-stress and stress conditions. (A) Schematic showing the *ATF4* mRNA with the sequences of three primer pairs amplifying three different amplicons (A1 – 3; indicated also in Fig. S6E): 1a-1b (the latter shown in the 3’ to 5’ direction for better illustrative purposes) for the putative queuing fragment, and 2a-2b and 3a-3b for two control fragments downstream of SL3 used in the ribosome-protection assay. (B) HEK293T cells were cross-linked with formaldehyde (HCHO) and then subjected to the ribosome-protection assay as described in Materials and Methods. qPCR product levels of the recovered putative queuing region (amplicon A1) are normalized to the region immediately downstream of SL3 (amplicon A2), as well as to the internal RNA isolation control (SPIKE) with non-stress values set to 1. Results are representative of three independent replicates and values are expressed in mean ± SD. Statistical significance was assessed using unpaired, two-sided, t-test (*p<0.01, **p<0.001) with Bonferroni correction. (C) HEK293T cells were treated with cycloheximide (non-crosslinking agent) and then subjected to the ribosome-protection assay as described in Materials and Methods. Results from three independent replicates were analyzed as described in panel B with non-stress values set to 1 (*p<0.01, **p<0.001). (D) HEK293T cells were transiently transfected with plasmids carrying either wt or SL3-mutated (in SL3^Mut-1^) *ATF4* reporters and treated as described in panel B. Results from three independent replicates were analyzed as described in panel B with the wt values set to 1 (*p<0.01, **p<0.001, ***p<0.0001).

Importantly, to demonstrate that it is the SL3 that prompts ribosome queuing, we repeated the ribosome-protection assay, but instead of non-transfected cells, we employed HEK293T cells transiently transfected with plasmids carrying either wt or SL3-mutated (SL3^Mut-1^) *ATF4* reporters. As shown in Figures 6D and S6C (for raw data see Supplementary Excel File 1), the SL3 elimination reduced significantly the difference in fragment protection efficiency of Amplicon 1 over A2 and A3 under both non-stress (by >2-fold) and stress conditions (by >3-fold). The fact that the fragment protection was not abolished completely by the SL3^Mut-1^ mutation is naturally caused by the presence of the endogenous *ATF4* mRNA with the fully preserved SL3.

Finally, we employed the recently developed RiboCrypt tool (https://ribocrypt.org) to analyze the disome-seq data (mRNA fragments protected by two stacked ribosomes) in a study where eIF5A, generally recognized as a ribosome rescue factor at poly-proline or proline–proline–glycine stretches (Schuller et al., 2017; Sfakianos et al., 2022), was knocked-down (Han et al., 2020). We observed by far the most prominent disome peak downstream of the uORF2 start codon (5‘ ends of the reads were mapped), followed by a much smaller and less sharp peak about 60 nt downstream, and another smaller peak upstream of SL3; i.e. another 60 nt downstream, in wt cells (Fig. S6E).

Thus, compared with the rest of the leader sequence, except for the extreme 5’ disome peak of unknown origin, the uORF2/ATF4 overlap region showed considerable disome coverage, with the two peaks set apart by 60 nucleotides (the expected size of a disome). This fits well with our model predicting a queue of five ribosomes, with the first one in the queue located with its P site on the AUG of uORF2, together covering the region of roughly 150 nt. Interestingly, in the eIF5A knockdown cells, the major peak downstream of uORF2 practically disappeared, as did the extreme 5’ disome peak, while the coverage of the entire stretch of the following 60 nt downstream of AUG of uORF2 increased substantially. Although we cannot explain the loss of the two major peaks, we propose that the increased coverage of this relatively broad region suggests defective collision clearance due to the lack ofeIF5A, further supporting our queuing hypothesis.

### uORF2 translation under stress is more prevalent than expected

To investigate characteristics of uORF2 translation under stress further, we next tested for evidence of uORF2 to ATF4 frameshifting. As presented in detail in SI, multiple approaches completely ruled out this possibility but clearly suggested that ATF4 might be translated from several alternative start sites, and all the resulting variants are induced by uORF2 (Figures S7–S9, and Tables S11 – S14).

Noteworthy, the final piece of evidence for the absence of frameshifting arose from an insertion of a 10x c-Myc tag into the uORF2 frame exactly 21 nt downstream of its AUG (Fig. S7B, uORF2_ins). Strikingly, using this construct, we also observed that the 10x c-Myc tag insertion in uORF2 behaved similarly to the 6x c-Myc insertion past the AUG1 of *ATF4* with respect to the role of SL3. In contrast to wt, where mutating inhibitory SL3 increased the ATF4 expression (Fig. 4A), unfolding SL3 in the 10x c-Myc tag insertion in uORF2 had no effect (Fig. S10A, left column, Table S15 and S16) under non-stress, same as in the case of the 6x c-Myc insertion past the AUG1 of *ATF4* (Fig. S10B, Table S17 and S18), and even decreased ATF4 expression under stress. Note that the uORF2 levels (probed with c-myc) seemed to be unchanged +/- SL3 under both conditions (Fig. S10A, right column, Table S15 and S16), as would be expected. Importantly, even with this SL3 mutant and uORF2-extended construct, we reproduced the unexpectedly small reduction (by ~40%) in the uORF2-cMyc expression level under Tg stress (compare Fig. S10A, right column, “fold-induction”, and Fig. S7B), further strengthening the interpretation that uORF2 is translated at much higher level even under stress than could be expected based on the original model. Collectively, our data suggests that the exact placement of SL3 within the uORF2 coding sequence with a defined length represents a precise molecular design that expands the existing delayed REI model to a more complex model that includes ribosome queuing to ensure well-tuned stimulation of ATF4 expression.

### mRNA methylation further fine-tunes ATF4 translation

It has been proposed that under non-stress conditions, A225 in the non-overlapping region of uORF2 in mouse *ATF4* mRNA is modified by N^6^-methyladenosine (m^6^A), which functions as a barrier blocking access of ribosomes scanning downstream of uORF1 towards the *ATF4* ORF (Zhou et al., 2018). (Please note that in the NCBI Reference Sequence NM_001287180.1 used here, mouse A225 corresponds to mouse A258; however, for clarity, we refer to it as it was originally described - A225.) Upon stress, this m^6^A is supposedly demethylated, thereby unblocking access. Of the four potential methylation sites we identified computationally, we chose to investigate the effect of human A235 (NCBI Reference Sequence NM_182810.2), as it seemed to match the mouse A225 best with respect to position (not sequence), and human A326, which is located within SL3 and could thus modify its function.

Both sites, highly conserved among vertebrates (Fig. S1A), were substituted with unmodifiable Gs. While A235G significantly increased (by 30% compared to wt) the basal level of ATF4 expression under non-stress, as originally reported (Zhou et al., 2018), A326G reduced expression by ~20% at both conditions (Fig. 7B, left and middle panels; Table S19 and Table S20), indicating that they may have opposing effects. To support these findings, we performed a T3-ligation assay (Liu et al., 2018b; Zhang et al., 2019) and investigated changes in the modification (possibly methylation) status of A235 and A326 in native *ATF4* transcripts isolated from a pool of poly(A) mRNAs from HEK293T and HeLa cells under non-stress and stress conditions. The T3-ligase is sensitive to modification sites like m^6^A during the ligation reaction in qPCR reactions (Liu et al., 2018b; Zhang et al., 2019). We first designed specific L (left) and R (right) probes based on the sequences flanking the two prospective m^6^A sites of interest: the *ATF4-A235* (probes L1 and R1) and *ATF4-A326* (probes L2 and R2). After ligation, the resulting fragment was amplified by qPCR using universal primers that were complementary to the same adapter sequences attached to all L and R probes (Liu et al., 2018b). Because presence of m^6^A significantly reduces the T3-ligase efficiency, less effective qPCR amplification determines under which conditions (non-stress *versus* stressed) the particular A is modified (possibly methylated). For control and normalization purposes, we also designed two additional probe sets complementary to two regions bearing A residues, but not present in any DRACH motifs: *ATF4-A267* (probes L3 and R3) and *ATF4-A311* (probes L4 and R4). Any changes in the amplification patterns of qPCR fragments resulting from their ligation therefore served as a quantitative measure reflecting changes in the abundance of *ATF4* transcripts in total poly(A) mRNAs isolates under either non-stress or stress conditions.

**Figure 7.**
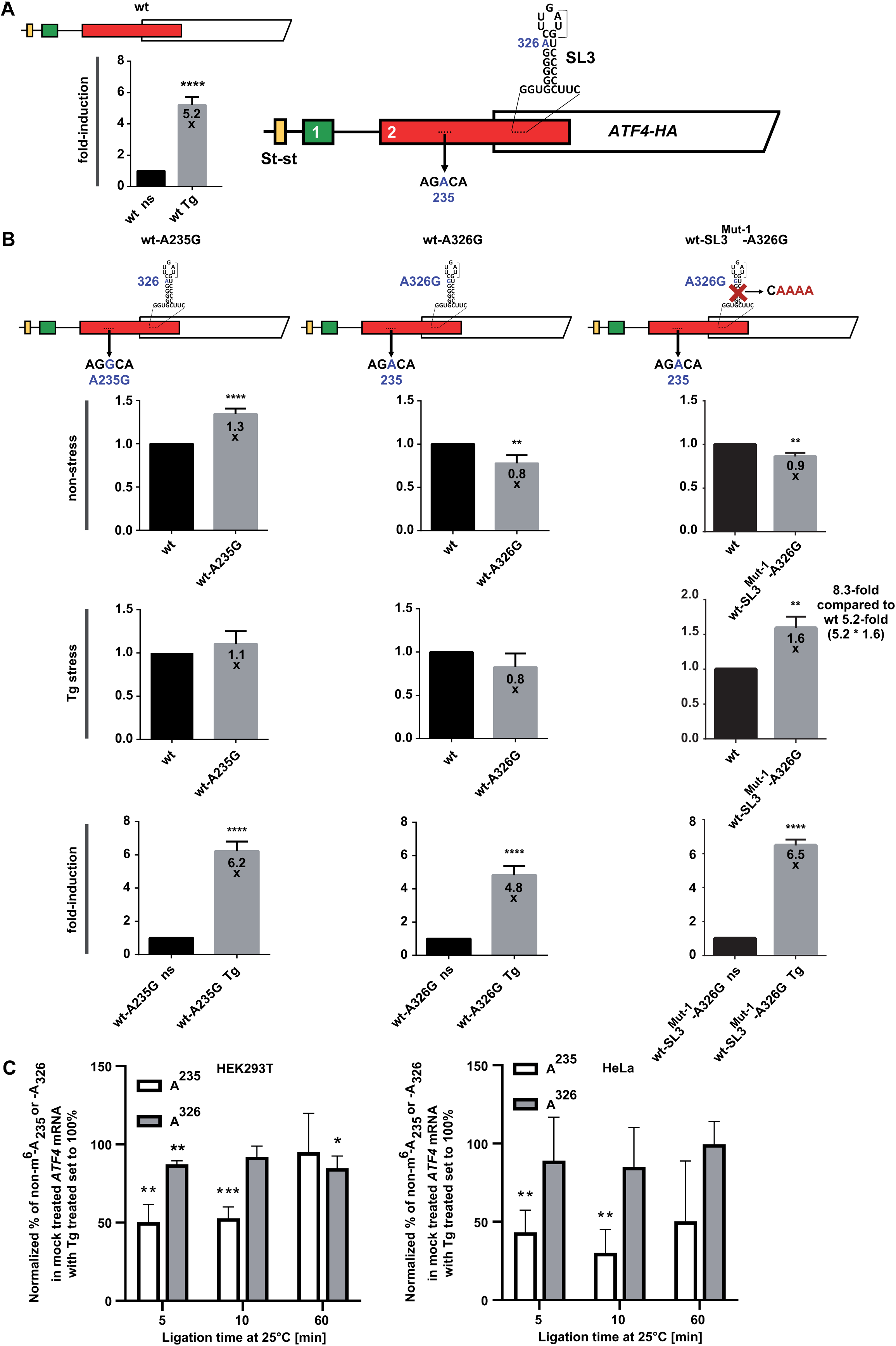
mRNA methylation further fine tunes ATF4 translation. (A) Same as Figure 1E for better comparison. (B) Same as in Figure 2A except that the A235G (left panel) and A326G either alone (middle panel) or in combination with SL3^Mut-1^ (right panel) *ATF4* mutant constructs depicted at the top of the corresponding panels were subjected to JESS analyses. (C) mRNA fragments prepared from either HEK293T (top panel) or HeLa (bottom panel) cells carrying the A235G and A326G mutations were subjected to T3-ligation assay as described in the main text. Normalized percentage of unmodified A_235_ or A_326_ bases of the *ATF4* mRNA expressed in mock treated *versus* Tg treated cells, with the latter set to 100%, was plotted shown.

Ligation reaction time intervals of 5 and 10 minutes revealed that the *ATF4* mRNA region containing the A235 but not the A326 residue had a ~30-50 % reduction in ligation efficiency under non-stressed conditions (Fig. 7C, Table S21). This result suggests that modification (possibly methylation) of A235 might indeed contribute to inhibition of *ATF4* translation under normal conditions. To the best of our knowledge, modification at A235 has not yet been described for human *ATF4* mRNA. Therefore, we do not know if a modification at A235 involves m^6^A, m^6^Am,m^1^A or something else. Nonetheless, the direct detection of m^6^A in the homologous region of the mouse *ATF4* transcript by (Zhou et al., 2018) with a similar phenotype when mutated strongly suggests that human *ATF4* A235 is also modified and serves as a barrier to ribosome progression, like SL3, but only under non-stress conditions.

Although our T3-ligation assay did not confirm that A326 is also modified, the fact that it may stimulate basal ATF4 expression, at least based on our reporter assays (Fig. 7B), and that it is situated in the open loop of inhibitory SL3, prompted us to test the effect of the A326G SL3^Mut-1^ and A235G SL3^Mut-1^ (as a control) double mutations. Whereas the latter phenocopied the effect of SL3^Mut-1^ alone under both conditions tested (data not shown), the former double mutation abolished the stimulatory effect of the SL3 elimination on ATF4 expression, same as the CUG^Mut^, but only under non stress conditions (Fig. 7B, right panel; Table S22 and S23).

These findings indicate that the residue A326 plays a stimulatory role under non-stress conditions and acts in concert with SL3, of which it is a part. Overall, our data suggest the importance of the A326 residue specifically under non-stress conditions, as its mutation to G renders SL3 even more inhibitory. At the same time, it neutralizes a stimulatory effect of the SL3 elimination. To reconcile these ostensibly contradictory observations, we propose that under normal conditions, potentially modified A326 destabilizes the inhibitory SL3 to balance its effect, but requires SL3 to be unfolded to fully unleash its stimulatory potential. The precise molecular mechanism is unknown, nor is it not known why it does not work the same way under stress conditions.

## DISCUSSION

The work presented capitalizes on an entirely new human ATF4-based reporter using an experimental workflow that avoided normalizing ATF4 expression levels to any reference genes whose expression would be translationally shut down under stress conditions. It extends the original model of delayed reinitiation to control *ATF4* translation and explains some newer evidence that have challenged some of its paradigms. In particular, we uncovered a new layer of regulation of ATF4 expression implemented by stable stem-loop (SL3) immediately downstream of a substantially leaky AUG1 of *ATF4,* co-operating with a near-cognate start codon (CUG), all inside the region overlapping with uORF2. Detailed analysis of this uORF2/ATF4 overlap region suggests ribosome queuing upstream of SL3, which may result in initiation at CUG and unexpectedly high translation of uORF2 even under stress, which the original model did not consider. Furthermore, our results confirmed, for human *ATF4*, a translation inhibitory role of m^6^A modification of A235 observed in mouse (Zhou et al., 2018) and identified a new potential modification site (A326), antagonizing the SL3 function under normal conditions. Therefore, we propose that the *ATF4* regulatory “armamentarium” is much more complex than previously thought.

Unexpectedly high uORF2 translation under stress was initially reported by Starck et al. (Starck et al., 2016), revealing a high degree of peptide expression from both uORF1 and uORF2 during both normal growth and stress conditions. The results were obtained from constructs fusing the uORF with a tracer peptide to the respective sequence. The observation was supported by results from ribosome profiling experiments by different groups (Andreev et al. 2015; Sidrauski et al. 2015). To explain this discrepancy with the original model, it was suggested that translation of the *ATF4* coding region under stress is a combination of reinitiation and leaky scanning past both uORFs. Given the complexity of the *ATF4* leader, it is conceivable that the tracer peptide employed by Starck et al. (Starck et al., 2016) could have unintentionally affected the results, e.g. by separating and/or breaking the additional *ATF4* regulatory elements we identified here. However, our novel reporter system not only confirmed the reported high levels of uORF2 expression under stress conditions, it additionally proposed a mechanistic explanation for this phenomenon based on ribosome queuing ahead of SL3.

### Ribosome queuing in the uORF2/ATF4 overlap region

The mechanism of ribosome queuing is known from other systems, such as translation of the antizyme inhibitor 1 (AZIN1) (Ivanov et al., 2018). The mRNA leader of *AZIN1* contains a relatively long, inhibitory, and non-AUG-initiated upstream conserved coding region (uCC), with a conserved, poly-amine-dependent Pro-Pro-Trp (PPW) motif pausing elongating ribosomes. At low polyamine levels, the majority of scanning ribosomes skip this uCC and translate AZIN1; those that infrequently initiate at its AUU start site will translate the uCC, terminate, and become recycled. High polyamine levels, however, interfere with eIF5A function and cause those very few ribosomes translating the uCC to stall at PPW. Subsequently scanning ribosomes, which again mostly skip the AUU start codon, as well as the occasional elongating ribosomes form a queue upstream of the PPW-stalled ribosome. This queue then stimulates uCC initiation at the AUU as scanning ribosomes are jammed all the way up to this site, reinforcing the PPW-imposed elongation stall to efficiently suppress AZIN1 synthesis.

The new layer of *ATF4* translation control that we propose is at least in part analogous to the AZIN1 mechanism. Specifically, the scenario with 80S ribosomes elongating from the AUG of uORF2 and subsequently queuing upstream of SL3 under non-stress conditions (Fig. 2B) – which prevents translation of ATF4 and ultimately also that of uORF2 – is similar to the AZIN1 system at high polyamine levels. This way, the queuing would re-inforce uORF2 initiation and very efficiently limit skipping of its AUG. It would also minimize the leakiness of the whole system to the lowest possible level: at zero demand, there is zero ATF4 synthesis. Upon stress, SL3 could slow down 80S ribosomes elongating from the AUG of ATF4, allowing for the formation of a queue consisting of both scanning 48S PICs and elongating 80S ribosomes (Fig. 2B).

Interestingly and somewhat paradoxically, we also found that removal of a potential modification of A326 upon stress would further stabilize SL3. We suggest that the SL3-imposed queue could then (i) enhance selection of alternative start codon(s) in *ATF4*, and (ii) prevent ATF4 over-induction. This process would involve reduced but still substantial translation of uORF2 even under stress, as has been observed. We further speculate that the near-cognate CUG could serve as another warrant licensing ATF4 synthesis, overcoming the leakiness of the ATF4 start site(s) (Fig. 2B). It seems that the ATF4 queuing system is well-optimized for its complex function, as our data imply that it depends strongly on the precise spacing between SL3 and upstream start codons (either uORF2 or ATF4). When we increased this spacing, either through the uORF2 or ATF4 reading frame, the tight ATF4 control was relaxed.

Importantly, in addition to the reporter analysis, several other observations supported the ribosome queuing hypothesis: 1) the published disome-seq data suggested formation of roughly two disomes (80S couples) in the region spanning the uORF2 AUG and SL3 whose distribution changes upon eIF5A knockdown (Fig. S6E); 2) our ribosome-protection assays indicated a specific protection of 132 nt-long fragment in the same region depending on intact SL3 (Fig. 6 and S6A - C); and 3) analysis of publicly available ribo-seq libraries generated from cells exposed to various ER stressors revealed prominent FP peaks with a strong preference for the CUG placement in the P-site (Fig. S4 and S5). All these results are consistent with a queue of 4 to 5 ribosomal species (either 80S couples or a mixture of 80Ses and 48S PICs) accumulated between uORF2 AUG and SL3.

It is understandable that given the variable length of mRNA FPs of 48S PICs compared to the relatively well-defined FP length of elongating 80S (Archer et al., 2016; Ingolia, 2010; Wagner et al., 2022; Wagner et al., 2020), it is difficult to predict the exact position of the most 5’ ribosomal species. This is made even more difficult because, as suggested by (Ivanov et al., 2018), ribosome queuing forces initiating ribosome to spend more time near the start codon where it can migrate back and forth by ~15 nucleotides, as previously shown (Matsuda and Dreher, 2006). Therefore, the operational space for the last 48S PIC in the queue to find the translation initiation site is relatively wide, increasing the probability for initiation even if the start codon is not in an optimal context. Concordantly, altering the spacing between uORF2 and SL3 cancels their functional interaction, indicating that the defined length of the queue for exerting its role is critical.

### Limitations of the study

One of the limitations of this study is that data obtained with our new reporter system, which we consider to be otherwise robust, are not supported by insertion of key mutations directly into the endogenous *ATF4* locus by for example CRISPR-based gene editing. Although they were not feasible given the scale of this study, this will be addressed by the ongoing experiments. Another issue to consider is that owing to the fact that our reporter mRNAs were highly overexpressed in cells (see SI), we recorded slightly increased ATF4 expression under non-stress conditions even with the wt reporter. This was, however, highly desirable because at least minimal ATF4 expression under non-stress conditions was instrumental for monitoring and comparing effects of our mutations also on the basal level of ATF4 expression. Finally, more experiments are needed to demonstrate that A235 and A326 are indeed modified, possibly by m6A, in human cells and that ATF4 can be stably expressed from the near-cognate CUG codon upstream of SL3.

### General considerations

Based on mounting evidence from the literature, it is now well-established that the precise mode of ISR induction, whether through UPR activation, starvation, or inhibition of tRNA synthetases, likely has very distinct downstream effects that manifest in highly nuanced *ATF4* translational control followed by unique modes of ATF4-mediated transcriptional responses (Neill and Masson, 2023). In light of this high degree of plasticity both in the control of ATF4 expression and its subsequent regulatory effect on its own transcriptional “regulon”, it seems plausible that a complex network of translation regulatory modes, as described here, is required to adjust ATF4 expression to a variety of conditions and tissues. Given the involvement of deregulated ATF4 expression in various pathologies and cancers, the complexity of its translational control should be kept in mind when considering therapies directed against this potent regulator of cell life or death.

## ACKNOWLEDGEMENTS

We are grateful to Dmitry E. Andreev, Pavel Baranov and Nick Guydosh for their timely advice and all past and present members of the Valasek lab for fruitful discussions and patience with this lengthy project. This work was supported by a Grant of Excellence in Basic Research (EXPRO 2019) provided by the Czech Science Foundation (19-25821X) and CZ.02.01.01/00/22_008/0004575 RNA for therapy by ERDF and MEYS (both to L.S.V.), US National Institutes of Health 5R35GM127089 and Chan Zuckerberg Initiative (to C.V), and US National Institutes of Health DK060596 (to M.H.). We also acknowledge Structural Mass Spectrometry Core Facility of CIISB, Instruct-CZ Centre (LM2023042) and European Regional Development Fund-Project “UP CIISB” (CZ.02.1.01/0.0/0.0/18_046/0015974) supported by MEYS CR.

## AUTHOR CONTRIBUTIONS

L.S.V. conceived and designed the project.

A.M.S., V.H., P.M.M., S.G., D.P., P.H., M.Ś., A.H., P.B., J.R., S.C. and K.J. carried out all experiments and analyzed the data.

L.S.V. interpreted results and wrote the paper with input from A.M.S., S.G., and C.V., and partly also from P.M.M., D.P., P.H., J.R., M.Ś., and M.H..

## STAR★Methods

### KEY RESOURCES TABLES

| REAGENT or RESOURCE | SOURCE | IDENTIFIER |
| --- | --- | --- |
| Antibodies |  |  |
| Mouse monoclonal anti-HA tag [HA.C5] | abcam | Cat# ab18181;<br>RRID:AB_444303 |
| Rabbit monoclonal anti-ATF-4 (D4B8) | Cell Signaling | Cat# 11815S;<br>RRID:AB_2616025 |
| Mouse monoclonal anti-ATF4 (B-3) | Santa Cruz | Cat# sc-390063;<br>RRID:AB_2810998 |
| Mouse monoclonal anti-c-Myc tag [9E10] | Dr. Tomáš Vomastek | N/A |
| Bacterial and virus strains |  |  |
| DH5alpha | Invitrogen | Cat# EC0112 |
| DH10B | Invitrogen | Cat# 18290015 |
| Chemicals, peptides, and recombinant proteins |  |  |
| Dimethyl sulfoxide (DMSO) | Sigma | Cat# D2438-5X |
| Thapsigargin | Invitrogen | Cat# T7458 |
| Tunicamycin | Sigma-Aldrich | Cat# T7765-1MG |
| Glo Lysis Buffer | Promega | Cat# E2661 |
| TurboFect | Thermo Fisher | Cat# R0532 |
| RNA blue | Top-Bio | Cat# R013 |
| UltraPure Distilled Water | Thermo Fisher | Cat# 109977-035 |
| Chloroform | VWR BDH Chemicals | Cat# 22711.290 |
| 2-Propanol | VWR BDH Chemicals | Cat# 20842.312 |
| Ethanol 96% vol | VWR BDH Chemicals | Cat# 20822.290 |
| Glycogen | Thermo Fisher | Cat# AM5910 |
| High-capacity cDNA reverse Transcription Kit | Applied Biosystems | Cat# 4368814 |
| RNase Inhibitor | Applied Biosystems | Cat# N8080119 |
| TURBO DNase | Invitrogen | Cat# 2238G |
| 10xTURBO DNase Buffer | Invitrogen | Cat# 8167G |
| DNase Inactivation Reagent | Invitrogen | Cat# 8174G |
| HOT FIREPol EvaGreen qPCR Mix Plus 5x | Solis BioDyne | Cat# 08-25-00020 |
| 10xThermoPol Reaction Buffer | NEB | Cat# B9004S |
| Vent DNA Polymerase | NEB | Cat# M0254L |
| 1x T3 ligation buffer | NEB | Cat# B0535S |
| T3 DNA ligase | NEB | Cat# M0317L |
| Plasmid Buffer P1 | QIAGEN | Cat# 19051 |
| RNase A | QIAGEN | Cat# 19101 |
| Plasmid Buffer P2 | QIAGEN | Cat# 19052 |
| Plasmid Buffer N3 | QIAGEN | Cat# 19064 |
| Plasmid Buffer PE | QIAGEN | Cat# 19065 |
| Plasmid Buffer PB | QIAGEN | Cat# 19066 |
| Buffer QG Solubilization Buffer | QIAGEN | Cat# 1014876 |
| T4 DNA Ligase | Roche | Cat#<br>10716359001 |
| Ligation Buffer, 10x | Roche | Cat#<br>11243292001 |
| Quick CIP | NEB | Cat# M0525S |
| Midori Green Direct | NIPPON Genetics<br>EUROPE | Cat# MG06 |
| BstBI | NEB | Cat# R0519S |
| HindIII-HF | NEB | Cat# R3104S |
| HpaI | NEB | Cat# R0105S |
| EcoRV-HF | NEB | Cat# R3195S |
| PstI-HF | NEB | Cat# R3140S |
| rCutSmart Buffer | NEB | Cat# B6004S |
| Criterion TGX Precast Gels 4-20 % | BIO-RAD | Cat# 5671093 |
| SuperSignal West Femto Maximum Sensitivity<br>Substrate | Thermo Fisher | Cat# 34096 |
| Bio-Rad Protein Assay Dye Reagent<br>Concentrate | BIO-RAD | Cat# 5000006 |
| GeneArt Strings DNA Fragments | Invitrogen | N/A |
| EcoSpin Spin Columns for DNA | Epoch Life<br>Science | Cat# 1910-25 |
| SUPERaseIn RNase inhibitor | Invitrogen | Cat# AM2694 |
| Ambion™ RNase I | Invitrogen | Cat# AM2294 |
| Tissue culture test plate, 6 wells | Techno Plastic<br>Products | Cat# 92406 |
| Critical commercial assays |  |  |
| 12-230 kDa Fluorescence Separation Module | Protein Simple bio-techne | SM-FL004-1 |
| 8xEZ Standard Pack 1 | Protein Simple bio-techne | PS-ST01EZ-8 |
| Anti-Mouse Detection Module | Protein Simple bio-techne | DM-002 |
| Protein Normalization Module | Protein Simple bio-techne | DM-PN02 |
| Deposited data |  |  |
| Analyzed data | (Ichihara et al.,<br>2021) | GSE174329 |
| Analyzed data | (Rendleman et al.,<br>2018) | GSE113171 |
| Analyzed data | (Andreev et al.,<br>2015) | GSE55195 |
| Experimental models: Cell lines |  |  |
| HEK293T | ATCC | CRL-3216;<br>RRID:<br>CVCL_0063 |
| HeLa | ATCC | CCL-2; RRID:<br>CVCL_0030 |
| Oligonucleotides |  |  |
| See Table S25 for a list of oligonucleotides |  |  |
| Recombinant DNA |  |  |
| *pCMV-EGFP-N2* | (Roithova et al.,<br>2020) | N/A |
| wt *ATF4*-HA Tag | This paper | N/A |
| dSt-st | This paper | N/A |
| d1 | This paper | N/A |
| d2 | This paper | N/A |
| d-all | This paper | N/A |
| St-st-only | This paper | N/A |
| uORF1-only | This paper | N/A |
| uORF2-only | This paper | N/A |
| dSt-st-NC1 | This paper | N/A |
| St-st-only-NC1 | This paper | N/A |
| St-st-only-NC2/3 | This paper | N/A |
| dSt-st-SL1 | This paper | N/A |
| St-st-only-SL2-1 | This paper | N/A |
| St-st-only-SL2-2 | This paper | N/A |
| dSt-st-SL2-1 | This paper | N/A |
| dSt-st-SL2-2 | This paper | N/A |
| d-all-SL2-2 | This paper | N/A |
| wt-A235G | This paper | N/A |
| wt-SL3<sup>Mut-1</sup> | This paper | N/A |
| wt-SL3<sup>Mut-2</sup> | This paper | N/A |
| dSt-st-SL3<sup>Mut-1</sup> | This paper | N/A |
| d-all-SL3<sup>Mut-1</sup> | This paper | N/A |
| uORF2-only-SL3<sup>Mut-1</sup> | This paper | N/A |
| wt-AUG3<sup>Mut</sup> (3<sup>rd</sup> *ATF4* AUG) | This paper | N/A |
| dSt-st-AUG3<sup>Mut</sup> (3<sup>rd</sup> *ATF4* AUG) | This paper | N/A |
| wt-UUG<sup>Mut</sup>-AUG3 | This paper | N/A |
| wt-SL3<sup>Mut-1</sup>-AUG3<sup>Mut</sup> | This paper | N/A |
| wt-A326G | This paper | N/A |
| wt-SL3<sup>Mut-1</sup>-A326G<sup>Mut</sup> | This paper | N/A |
| wt-3AUG<sup>Mut</sup> | This paper | N/A |
| d2-3AUG<sup>Mut</sup> | This paper | N/A |
| 3AUG<sup>Mut</sup>-stop | This paper | N/A |
| 3AUG<sup>Mut</sup>-short | This paper | N/A |
| 3AUG<sup>Mut</sup>-short-stop | This paper | N/A |
| uORF2_ins | This paper | N/A |
| d2_ins (uORF2_ins => AUG to AGG) | This paper | N/A |
| uORF2_ins_3AUG<sup>Mut</sup> | This paper | N/A |
| wt-CUG<sup>Mut</sup> | This paper | N/A |
| wt-SL3<sup>Mut-1</sup>-CUG<sup>Mut</sup> | This paper | N/A |
| wt_ins | This paper | N/A |
| wt_ins-SL3<sup>Mut-1</sup> | This paper | N/A |
| d2_ins | This paper | N/A |
| d2_ins-SL3<sup>Mut-1</sup> | This paper | N/A |
| uORF2_ins-SL3<sup>Mut-1</sup> | This paper | N/A |
| uORF2-*ATF4*-HA fusion | This paper | N/A |
| Software and algorithms |  |  |
| Compass for SW ver. 6.3.0 | ProteinSimple | https://www.biotechne.com/resources/instrument-software-download-center |
| GraphPad Prism ver. 10.1.2 | GraphPad Software, San Diego, CA, USA | RRID:SCR_002798 |
| SciPy ver. 1.10.1 | (Virtanen et al., 2020) | RRID:SCR_008058 |
| Python 3.11.3 ver. 8.1.1 | (Rossum and Drake, 2009) | https://www.python.org |
| Matplotlib ver. 3.7.0 | (Hunter, 2007) | RRID:SCR_008624 |
| R ver. 4.2.2: 'pheatmap' package ver. 1.0.12 | (Kolde and Kolde, 2015) | N/A |
| R ver. 4.2.2: 'RiboWaltz' package | (Lauria et al., 2018). | N/A |
| Other |  |  |
| Jess Automated Western Blot System | Protein Simple bio-techne | 004-650 |
| RiboCrypt tool | https://ribocrypt.org |  |
| G-Box imager | Syngene | N/A |
| Mendeley data | http://www.mendeley.com/ | RRID:SCR_002750; DOI: 10.17632/zmdvhwggg8.1 |
| Quantity One Software | Bio-Rad | RRID:SCR_014280 |

### RESOURCE AVAILABILITY

#### Lead contact

Further information and requests for resources and reagents should be directed to and will be fulfilled by the Lead Contact Leoš Shivaya Valášek.

#### Materials availability

Plasmid constructs generated in this study are available upon request.

#### Data and code availability

- The data generated in this paper and used for the preparation of main Figures is available in Tables S1–S23, as well as from the lead contact upon request. Additionally, this paper analyzes existing, publicly available data. These accession numbers for the datasets are listed in the key resources table. Data have been deposited at Mendeley and are publicly available as of the date of publication. Accession numbers are listed in the key resources table.
- This paper does not report original code.
- Any additional information required to reanalyze the data reported in this paper is available from the lead contact upon request.

#### Experimental model and study participant details

##### Human cell line

The HEK293T cell line (ATCC) was used for all experiments in this study; the HeLa cell line (ATCC) was used only for the T3-ligation assay. The source for commercial HEK293T as well as HeLa cell line is reported to be female. Cells were grown in Dulbecco’s modified Eagle’s Medium (DMEM) high glucose medium supplemented with 10% fetal bovine serum (FBS) at 37.0 °C with 5.0% CO_2_ concentration. Cells were seeded into 6-well plates (Techno Plastic Products) to 2.5 ml of medium per well and grown 24 hours prior to transfection at approximately 40-50% confluency.

##### Bacterial strains

Bacterial strains DH5α and DH10B (Invitrogen) were used in this study.

#### Method details

##### Construction of the ATF4-HA tagged plasmids

To create the ATF4-HA tagged reporter plasmids, the hATF4-wt-HA-Tag-3’UTR (NM_182810.2) ordered as GeneArt Strings DNA Fragment (Invitrogen) was cloned into the empty Clontech EGFP-N2 high copy number vector (Roithova et al., 2020), while removing the GFP insert. All hATF4-HA tagged constructs contain the CMV promoter, the full 5’UTR of ATF4, the full CDS of ATF4, and the natural 3’UTR of ATF4. The HA tag is placed at the very C-terminus of the ATF4 CDS just upstream of its stop codon; all these plasmids have the kanamycin resistance. Mutant constructs were generated using either GeneArt Strings (Invitrogen) or PCR using 10×ThermoPol Reaction Buffer (NEB), Vent DNA Polymerase (NEB) and specific primers; for the list of plasmids, details of their cloning, and lists of primers and GeneArt strings please see Supplemental Information (Tables S24 – S26). Plasmid isolation was performed using QIAGEN Plasmid Mini Kit and EconoSpin DNA Spin Columns (Epoch Life Science). QIAGEN QIAquick Gel Extraction Kit and QIAGEN QIAquick PCR Purification Kit (QIAGEN) were used for insert isolation.

##### Preparation of whole cell extracts

HEK293T cells pre-grown for 24 hours as described in the Human cell lines section were transfected with ATF4-HA reporter constructs using TurboFect transfection reagent (Thermo Fisher); 2.5 µg of plasmid DNA was used per one well in the 6-well plate in all experiments. Exactly 8 hours after transfection, cells were treated with either 1 µM thapsigargin (Invitrogen) for 3 hours or Dimethyl sulfoxide (DMSO) (Sigma) as a control. Cells were lysed directly on the plate using 200 µl Glo Lysis Buffer (Promega), which is recommended to be used with the Jess Automated Western Blot System (Protein Simple bio-techne). One half of the lysate was saved for RNA isolation followed by a qPCR control step and the other half of the lysate was subjected to JESS SW assay according to the supplier’s instructions. In the experiments with tunicamycin (Sigma-Aldrich), 8 hours after transfection, HEK293T cells were treated with either 0.5 µM tunicamycin for 4 hours or DMSO as a control. Harvesting of the cell lysates was performed essentially the same as for thapsigargin.

##### RT-qPCR control step

Total RNA was isolated using RNA blue reagent (Top-Bio) according to the manufacturer’s instructions. Subsequently, a TURBO DNase (Invitrogen) cleavage step was performed. For cDNA synthesis, 0.5 µg of total RNA was used for each sample using a High-Capacity cDNA Reverse Transcription kit (Applied Biosystems). At least three 10-fold serial dilutions (as described before (Chiu et al., 2010; Khoshnevis et al., 2014)) of cDNA for each mutant with the wt pair, examined in each individual experiment, were compared using RT-qPCR, and the maximum difference criterion of 1 cycle had to be met for all sample dilutions in all relevant controls (including control for the transfection efficiency) at the time of lysis, otherwise samples were discarded. To compare the reporter levels for all mutants, the reverse primer was designed to be in match with HA tag (to avoid interference with endogenous ATF4). Primers matching the plasmid region carrying neomycin/kanamycin resistance were created (to ensure the same level of transfection efficiency for each pair of samples). Internal controls for RNA isolation efficiency and cDNA synthesis were used as well (SPIKE RNA and the corresponding primers). The qPCR primers are listed in Table S25. For qPCR reactions, HOT FIREPol EvaGreen qPCR Mix Plus 5x (Solis BioDyne) was mixed with 0.8 µM primers and cDNA and run using the following program: 95 °C for 15 sec, followed by 43 cycles of 95 °C for 15 sec, 60 °C for 20 sec, and 72 °C for 20 sec.

##### JESS Simple Western (SW) assay ProteinSimple

JESS SW Assay was used according to the manufacturer’s protocol. In brief, the entire process in the Jess machine can be described as follows: proteins are separated by their molecular weight in a capillary pre-aspirated with separation and stacking matrixes. Once the separation is complete, UV light immobilizes the proteins to the capillary wall. After immobilization and clearing of the matrix from the capillary, the immunoprobing process is initiated, first by incubation with the primary antibody, then with the secondary HRP conjugate, and finally with the chemiluminescent substrate. The emitted chemiluminescent light is then recorded by a CCD camera and automatically quantified. For the experiments conducted as part of this study, the lysate was mixed with 1× Sample Buffer in accordance with the amount calculated in SW Sample Calculator for each sample, and subsequently Fluorescent 5× Master Mix containing DTT solution and 10× Sample Buffer was added (Protein Simple bio-techne). Twenty-four samples were tested in each JESS run. Samples were mixed by vortexing, boiled at 95 °C for 5 min and spun down using a benchtop microcentrifuge. Together with the tested samples, biotinylated ladder was pipetted into 12-230 kDa Pre-filled Plates from Fluorescence Separation Module (Protein Simple bio-techne). Mouse monoclonal anti-HA tag (abcam) or anti-c-Myc antibodies (provided by Dr. Tomáš Vomastek) were pipetted along with Anti-Mouse Detection Module (Protein Simple bio-techne). The plate was centrifuged for 5 minutes at 686 x g at room temperature and inserted into the JESS machine along with Fluorescent Capillary Cartridges. Data analysis and control of the results obtained by the JESS instrument were performed in Compass for SW software (version 6.3.0) (ProteinSimple).

The key point to mention is that the same amount of total protein loaded in all capillaries was pre-estimated using Bio-Rad Protein Assay Dye Reagent Concentrate (BIO-RAD) and BSA titration curve was performed before each JESS run with serial dilutions of BSA in PBS. Afterwards, Protein Normalization Module (Protein Simple bio-techne) was applied for a thorough comparison and finally only up to 20% difference between samples was tolerated using PN Module, otherwise samples were discarded. Please note that the 180 kDa peak, which occurs occasionally in electropherograms, represents (based on the manufacturer’s instructions) a non-specific peak arising from cross-reactions between fluorescence (PN kit) and chemiluminescence (target antibody) reagents during capillary detection. According to the manufacturer, the only occasionally nature of this peak is not known.

##### Human *ATF4* mRNA transcript sequence analysis

The V2 mRNA transcript of human *ATF4* was analyzed using freely available bioinformatics tools designed to predict RNA sequence and structural motifs, such as vsfold5, Hotknot, Pknot, pKiss, RECODE, PRFdb, SCRAMP, and BERMP.

##### Western Blot

All samples were resolved using Criterion TGX Precast Gels 4-20% (BIO-RAD) followed by western blotting. All primary antibodies used in this study are listed in key resources table. The signal was developed using SuperSignal West Femto Maximum Sensitivity Substrate (Thermo Fisher) and detected in a G-Box imager (Syngene) using series of varying exposure time. Data were analysed using Quantity One Software (Bio-Rad) and deposited at Mendeley; accession numbers are listed in the key resources table.

##### Ribosome protection assay

HEK293T cells were seeded in 20 mL of media in the 15 cm dish and grown to a confluency of 60-70% before being treated with either DMSO (control) or thapsigargin (1µM) for 3h. For experiments involving either wt or SL3^Mut-1^*ATF4* reporters (Fig. 7D and S5C), cells were independently transfected with either reporter plasmid at approximately 40-50% cell confluency and – 8 h after transfection – treated with DMSO (Control) or thapsigargin (1µM) for additional 3h. Cells were then treated with formaldehyde (“crosslinked”, incubated with HCHO 0.8% for 5 min at 4°C and the reactions were quenched with 75 mM glycine for 5 min, as described (Wagner et al., 2022; Wagner et al., 2020)) or cycloheximide (“non-crosslinked”; incubated with 100 µg/mL cycloheximide for 1 min at 37°C, as shown before (Herrmannova et al., 2020)), following which cell lysates were prepared. Concentration of cell lysates were measured on NanoDrop at OD_260_ and all samples were divided into two halves. One half (5 AU OD_260_ per 1 ml of a lysate) was treated with Ambion™ RNase I (4U) (Invitrogen) at 30°C for 10 min at 500 rpm to digest all enzyme-accessible RNA. The other half was processed the same way but without the RNase I treatment. Nuclease reactions were stopped with SUPERaseIn RNase inhibitor (8U) (Invitrogen). All samples (RNase I treated and non-treated) were supplemented with 1 ml of RNA blue (Top Bio) and mixed vigorously by inverting the tubes. To reverse crosslinking, formaldehyde-crosslinked samples were incubated at 65°C for 20 min with an intermittent shaking by inverting the tubes. Subsequently, all the samples were cooled down at RT for 5 min and 5µl of SPIKE RNA (*in vitro* transcribed yeast RPL41a) was added as an internal control of RNA isolation to all the samples and RNA was isolated according to the manufacturer’s (Top Bio s.r.o) instructions. The resulting RNA pellets were dissolved in 40 µl of nuclease free water. RNA concentrations were measured using NanoDrop. To remove any DNA contaminants, 2.5 µg of RNA samples were treated with TURBO™ DNase (Invitrogen) according to the manufacturer’s instructions. For cDNA synthesis, 1 µg of DNase-treated RNA was used for each sample using a High-Capacity cDNA Reverse Transcription kit. For qPCR reactions, HOT FIREPol EvaGreen qPCR Mix Plus 5x was mixed with 0.4 µM primers (for the SPIKE primer set) and 0.25 µM (for all Amplicon 1 - 3 primer sets) primers and cDNA, and run using the following program: 95 °C for 15 sec, followed by 40 cycles of 95 °C for 15 sec, 57.4 °C (for SPIKE primer set) or 62 °C (for all Amplicon 1 - 3 primer sets) for 20 sec, and 72 °C for 20 sec. Melting curve was measured between 60-90 °C to assess amplification of the single specific product. qPCR data was analyzed as described in the main text and corresponding figure legends.

##### Total and poly(A) RNA isolation

Total RNA for T3-ligation assay was isolated using the RNA Blue reagent (Top-Bio) according to the manufacturer’s instructions, 5 ml of the reagent was used *per* one 15 cm dish containing HEK293T or HeLa cells grown to approximately 80% confluency. The resulting RNA pellets were resuspended in RNase-free water (Thermo-Fisher) and the concentration was quantified by NanoDrop. Subsequently, poly(A) RNA for T3-ligation assay was purified using the Poly(A)Purist™ MAG kit (Thermo Fisher) according to the manufacturer’s instructions. The poly(A) RNA was stored in 70% ethanol at −80°C and the subsequent steps of precipitation and precipitate resuspension in RNase-free water were finished just before proceeding to the T3-ligation assay.

##### T3-ligation assay and qPCR

The T3-ligation assay was adopted from (Liu et al., 2018a) and (Zhang et al., 2019). Briefly, each ligation reaction mixture A consisted of 20 nM of corresponding probe SGL (Table S25), 20 nM of corresponding probe SGR, 1x T3 ligation buffer (NEB), and ~200-500 ng of poly(A) RNA. The ligation reaction mixture B consisted of T3 DNA ligase (NEB) diluted to 10U/µl with ligation buffer. Mixtures A were heated at 85°C for 3 min and then incubated at 35°C for 10 min, and then the ligation reaction mixtures B were added. The final volumes of the ligation reaction mixtures were 10 μl and contained 10U of T3 DNA ligase. These resultant mixtures were then incubated at 25°C for 5, 15 or 60 min and chilled on ice immediately. qPCR was carried out according to vendor’s instructions (Solis BioDyne) using Bio-Rad CFX384 Real-Time PCR System. The 10 µl qPCR reactions contained 1x HOTFIREPol EvaGreen qPCR Mix Plus, 200 nM of each adapter primer (SGadapterFOR and SGadapterREV) and 1 μl of individual ligation reactions and were run using the following program: 95°C for 15 min followed by 44 cycles of 95°C for 15 sec, 60°C for 20 sec, and 72°C for 20 sec. Melting curves were analyzed between 65 and 90°C. Results were analyzed using Bio-Rad CFX Manager. Cycle differences between thapsigargin-treated and mock-treated samples were calculated for each individually analyzed region and the resultant ratios were then used for the normalization of each of the investigated regions (containing either *ATF4* A_235_ or A_326_) to both control regions (containing *ATF4* A_267_ and A_311_).

##### Ribosome profiling datasets analysis

Riboseq fastq files of untreated or sodium arsenite (GSE17432, GSE55195) or tunicamycin (GSE113171) treated samples from previously published datasets (Andreev et al., 2015; Ichihara et al., 2021; Rendleman et al., 2018) were used for analysis in this study. Reads devoid of rRNA and tRNA sequences were aligned to the human reference genome and transcriptome (GRCh38.p13, annotation release 109) by STAR aligner (2.7.10a) (Dobin et al., 2013) using local alignment with 10% mismatches, and with indels and soft clipping allowed for transcriptomic alignment. Transcriptomic alignments were further filtered to obtain unique alignments using custom programs (Herrmannová et al., 2023). The Unique transcriptomic alignments and MANE project (v. 1.0) annotation were used to generate ribosome foot-print coverage and P-site analysis plots of ATF4 transcript by RiboWaltz package (Lauria et al., 2018).

##### Quantification and statistical analysis

Variables were tested for normality using the Shapiro-Wilk normality test. Based on the normality test, the differences between experimental groups were tested by the t-test or by Mann–Whitney test (stated in the Figure’s legend). Variables are presented as mean ± SD and *p* values of < 0.05 were considered statistically significant. GraphPad Prism statistical software (ver. 10.1.2, GraphPad Software, San Diego, CA, USA, RRID:SCR_002798) and SciPy (ver. 1.10.1) (Virtanen et al., 2020), Matplotlib (ver. 3.7.0) (Hunter, 2007) libraries in Python (ver. 3.11.3) (Rossum and Drake, 2009) were used for statistical analyses and visualization. The heatmaps showing the differences in the relative expression of mutant *versus* WT constructs were created in R (ver. 4.2.2) using the ‘pheatmap’ package (ver. 1.0.12) (Kolde and Kolde, 2015). For the ribosome protection assay, statistical significance was assessed using unpaired, two-sided, *t*-test with Bonferroni correction.

##### Additional resources

This work does not use additional resources.

## SUPPLEMENTAL INFORMATION

### SUPPLEMENTAL RESULTS

#### Validation of the new ATF4-HA reporter system

To understand what lies behind the “background” ATF4 expression under non-stress conditions and to validate that our new reporter system faithfully mimics endogenous *ATF4* regulation, we used traditional western blot analysis. We reasoned that the non-stress peak could originate from the DNA transfection which is known to be stressful to cells (Andreev et al., 2016) and/or from CMV promoter-driven transcription generating a high level of the *ATF4-HA* mRNA, which could magnify the normally negligible leakiness of *ATF4* repression under non-stress conditions. As shown in Figure S1B, transfection of our wt reporter using turbofect, but not turbofect or DMEM alone, resulted in a mild ATF4 induction even without Tg treatment (lanes 2 and 8 *vs.* lanes 3, 4 and 9, 10). This result could suggest that DNA transfections trigger mild stress as reported before, however, control transfections of an empty vector did not trigger such a response (Fig. S1C). Therefore, we concluded that a high level of our CMV-driven reporter mRNA (>100 times higher than endogenous *ATF4* mRNA levels, as demonstrated below) resulted in detectable basal levels of ATF4 protein in otherwise unstressed cells, as also observed by others using the CMV-driven ATF4 expression (Iwawaki et al., 2017; Wang et al., 2022). Nonetheless, clear Tg-mediated induction of the endogenous ATF4 in lanes 6 and 7 in the “ATF4 (anti-ATF4)” panel of Figure S1B (compared to lanes 3 and 4), and of our ATF4-HA reporter in lane 5 in the “ATF4-HA (anti-HA)” panel (compared to lane 2), demonstrate that both ATF4 variants respond to Tg treatment in the same manner. Due to the absence of signal in lanes 3 and 4 we could not quantify the level of induction of the endogenous ATF4 alone. Nonetheless, the level of induction of ATF4-HA calculated from JESS (5.2±0.5) or western blots (3.9±1.3) and the level of induction of combined endogenous ATF4 and ATF4-HA (lanes 5 *versus* 2 in the “ATF4 (anti-ATF4)” panel showing the cumulative amounts of both HA-tagged and endogenous proteins that anti-ATF4 antibodies recognize) calculated from JESS (5.5±0.6) or western blots (3.4±0.2) correspond nicely to what we expected (Fig. S1D). Given the fully quantitative nature of JESS measurements, we consider them to be the most reliable.

Importantly, the observed induction pattern is solely attributable to *ATF4* translational control, as the mRNA levels of both *ATF4-HA* and endogenous *ATF4* (measured separately using highly specific reverse primers matching the *ATF4* stop codon region, by which these two alleles differ due to the HA tag sequence) remained virtually unchanged under non-stress *vs.* stress conditions (Fig. S2A). Collectively, these results document that our reporter mirrors the behavior of endogenous ATF4 regulation.

As aforementioned, when we compared *ATF4* mRNA levels under non-stress conditions using primer sets A1 and A2 downstream of SL3 (Amplicons 2 and 3 defined in the results chapter “Ribosome queuing contributes to the overall translational control of *ATF4*“ and Fig. S6), we revealed over 100-fold more *ATF4* mRNA in transfected (expressing both endo and exogenous *ATF4* mRNA) than in non-transfected (expressing endogenous *ATF4* mRNA alone) (Fig. S6D; for raw data see Supplementary Excel File 1).

#### St-st modestly inhibits ATF4 expression, uORF1 allows for 50% downstream REI, and a solitary, inhibitory uORF2 allows for high ATF4 stress-inducibility

Keeping St-st as the only element in the *ATF4* leader resulted in ~20.6-fold increase in ATF4 expression under non-stress conditions and ~3.1-fold increase under Tg stress over wt (Fig. S3A and B, and Table S3 and S4). The former value is only modestly but reproducibly lower than that of d-all (reduced by ~10%) supporting the idea that St-st acts as a general repressive element, i.e., a roadblock, as observed before (Rendleman et al., 2023; Wagner et al., 2020).

The uORF1-only construct, where only the AUG of uORF1 is preserved, led to 12.2-fold and 2.7-fold upregulation in non-stress and Tg stress conditions, respectively, while also showing no stress inducibility compared to wt (Fig. S3A - C, and Table S3 and S4). These results clearly confirm the stress-independent, REI-permissive nature of this uORF and indicate that more than 50% of ribosomes that translate uORF1 (~12-fold induction of uORF1-only over wt [Fig. S3A - row 1] divided by ~23-fold induction of d-all [Fig. S3A - row 1]) undergo partial recycling and reinitiate downstream upon reacquisition the TC.

According to the original delayed reinitiation model, uORF2 serves as a barrier capturing most, if of not all, 40S ribosomes that resumed scanning after termination at uORF1 under non-stress conditions, thus preventing ATF4 expression because its sequence extends into the *ATF4* CDS out of frame. Under stress conditions, characterized by low TC levels, its expression should be dramatically reduced, allowing ATF4 expression. In the wt set-up, uORF2 might still be expressed to some degree because not 100% of ribosomes skip its AUG even under decreased TC levels, but the amount of those that do should be substantially higher when compared to non-stress conditions. In any case, in the uORF2-only set-up, this strong barrier should theoretically not allow any significant inducibility of the reporter under stress conditions, because once the ribosome initiates at uORF2 as the first and only upstream uORF, the effect on ATF4 is independent of the TC levels.

Accordingly, we found that the uORF2-only construct showed an ~80% drop in reporter expression compared to the wt under both normal and Tg stress conditions indicating dramatically reduced initiation at *ATF4* (Fig. S3A and B – rows 1 and 2, and Table S3 and S4). However, in stark contrast to the above logic, comparing stress with non-stress values of uORF2-only (“fold-induction” – row 3) revealed an unexpected ~3.2-fold increase in ATF4 translation under stress that could not be explained by the original model. We argue that this large increase cannot be explained purely by the proposed increased leaky scanning over uORF2 under stress (Starck et al., 2016).

Next, we constructed mutants in which we eliminated one upstream element at a time and compared the resulting constructs with the wt set to 1 (Fig. S3B and D, and Table S3 and S4). Eliminating St-st (in dSt-st) showed no significant effect under both conditions. Nonetheless, we repeatedly observed higher, but not significant, stress induction of dSt-st compared to wt (increased by ~10% from wt ~5.2-fold to ~5.8-fold; Fig. S3C *vs.* S3D – row3, and Table S3 and S4), which argues for the St-st’s role as a stress-independent roadblock reducing processivity of the scanning ribosome. These results, and the fact that mouse and possum *ATF4* lacks the St-st, indicate that uORFs 1 and 2 are largely sufficient to ensure stress-induced translation of ATF4.

While removal of the REI-permissive uORF1 (in d1) significantly reduced ATF4 expression by ~70% under both non-stress and Tg stress, eliminating the inhibitory uORF2 (in d2) dramatically increased ATF4 expression (~12.1-fold) under non-stress conditions and slightly (~2.4-fold) under stress (Fig. S3B and D – rows 1 and 2, and Table S3 and S4), all as expected from the original model. The latter result nicely corroborates that ~50% of ribosomes terminating at uORF1 reinitiate downstream (~12-fold induction of d2 over wt [Fig. S3D - row 1] divided by ~23-fold induction of d-all [Fig. S3A - row 1]), as we observed with the uORF1-only construct (Fig. S3A).

Comparison of the stress induction potentials of each construct showed that d2 is very similar to uORF1-only and d1 to uORF2-only (Fig. S3D and S3A, “fold-induction” plots in the respective columns), except that the presence of the St-st further magnified the uORF2 inducibility (from ~3.2-fold to ~5.8-fold; “fold-induction” plots in Fig. S3A – uORF2-only *versus* Fig. S3D – d1). This result is again in line with St-st serving as a ribosome barrier to maintain low basal levels of ATF4 expression.

Taken together, our results confirmed major aspects of the original model but raised the question as to the mechanism behind the unexpected stress induction of the uORF2-only reporter.

#### Sequence analysis of the 5’ UTR of the human ATF4 mRNA reveals additional elements potentially contributing to ATF4 translational control

Here we asked if there are some so-far unidentified regulatory elements in the ATF4 mRNA leader besides St-st and both uORFs. We had previously predicted and experimentally validated the existence of two stem loops (SL1 and 2) preceding uORF1 (Fig. 2A), the first of which cooperates with eIF3 to unleash the full REI potential of uORF1 (Hronova et al., 2017). SL1 also contains one near-cognate CUG start site (NC1) that is immediately followed by two overlapping GUG near-cognates (NC2/3) that are no longer part of SL1.

Notably, the SL1 and its NC1, and NC2/3 are not universally conserved in eukaryotes (Fig. S1A). The St-st is conserved in most mammals examined, except mouse a few others. The SL2 is conserved in primates and, interestingly, its formation is even slightly more favorable in mouse and rat than in human. Indeed, the presence of uORF1 and uORF2 is well conserved.

The next element, predicted by our analysis, is another stem-loop (SL3), with ΔG = −15.40 kcal/mol, which could potentially be inhibitory and is highly conserved among vertebrates (Fig. S1A). It is located roughly in the middle of the uORF2/ATF4 overlap (Fig. 2A) and has a hairpin structure with the highest free energy predicted for this particular sequence region.

Further, our analysis predicted a total of four individual sites of m6A methylation within well-defined motifs (Fig. 2A); DRACH1 (with A225) is located in the non-overlapping, likely unstructured region of uORF2, followed by RRACH1 (A235) in the same region, which is homologous to the predicted methylated A in mouse mRNA (Zhou et al., 2018); RRACH (R = G or A; H = A, C or U), DRACH (D = A, G or U). The DRACH2 (A286) motif occurs in the uORF2/ATF4 overlap and covers the ATF4 main initiation codon with a modifiable adenine located just downstream of the AUG. Thus, its modification could interfere with AUG recognition during start site selection. Finally, RRACH2 (A326) locates to the stem of SL3; therefore, it could potentially affect its stability. Although the DRACH1 and 2 motifs are not universally conserved in eukaryotes, RRACH1 and 2 motifs are highly conserved in mammals (Fig. S1A) and could therefore play an important role in ATF4 regulation.

The ATF4 gene begins with two nearly consecutive AUGs followed by a third in close proximity, all in frame (Fig. 2A, in dark green). Both the canonical AUG1 and AUG2 have a medium Kozak initiation context; AUG1 overlaps with DRACH2, and AUG2 is located two codons downstream. AUG3 has a weak Kozak initiation context and is exposed in the open loop of SL3. It represents the 17th codon downstream of AUG1. Both AUG2 and AUG3 are highly conserved among vertebrates (Fig. S1A).

The uORF2/ATF4 overlap also contains two putative conserved sliding sequences (FS-A and -B) and a so-called C-tract motif (Penn et al., 2020), which could potentially prompt the elongating ribosome to switch the reading frame from uORF2 to ATF4. We investigated this hypothesis by examining the frame of ribosome footprints from a dataset collected from HeLa cells subjected to stress (Rendleman et al., 2018). Specifically, we mapped ribosome protected fragments in the region of the uORF2/ATF4 overlap and found that, during acute ER stress, reads continued to locate to the uORF2 reading frame (frame 0) and perhaps even increased, while there was no coverage of the ATF4’s frame (frame 1)(Fig. S2B). In fact, ATF4’s translation appeared to initiate downstream of the predicted SL3 after 4 hours of Tunicamycin stress.

Sliding region FS-A involves 3 consecutive glycine codons, and because of its overlap with SL3, it can only function if SL3 is not formed. Switching from uORF2 to the ATF4 reading frame could be mediated by either a +1 or −2 programmed ribosomal frameshifting (PRF). Of these two possibilities, the latter is much more likely because both types of glycine tRNAs, with GGG in the P-site and GGA in the A-site, are well positioned for re-pairing after a −2 shift (Penn et al., 2020). In support of this hypothesis, 8 nucleotides downstream from FS-A occurs the C-nucleotide-rich C-tract motif (CCCCCUUCGACC) with high similarity to the inhibitory C-motif promoting −2 PRF in arteviruses (Pakos-Zebrucka et al., 2016). Since 8 nucleotides is considered the optimal distance for the C-motif to act as the inhibitory element that can cooperate with the upstream sliding sequence to promote −2 PRF, we designed constructs to examine this option (see below).

FS-B is located just upstream of the uORF2 stop codon, thereby partially fulfilling the +1 PRF criteria (Penn et al., 2020). Yet, neither of its glycine codons had the potential to slow or stop ribosomes, as they belong to the high codon usage category. Since FS-B also did not meet the −2 PRF criteria, it likely has no role in ATF4 translational control.

Given this knowledge, we created an array of mutants for all these predicted elements (Fig. 2A), and first evaluated them in a large-scale screen. It resulted in no observable effect for mutations in NC1 to NC3 in SL1, both DRACH motifs, mutations unfolding SL1 or SL2 (specifically with respect to the St-st function), and mutations of AUG3 inside the ATF4 coding region (data not shown). Therefore, we focused our analysis on mutations in other elements that had observable effects.

#### uORF2 translation under stress is more prevalent than expected

To investigate characteristics of uORF2 translation further, we next tested for evidence of uORF2 to ATF4 frameshifting. To do so, we mutated the first 3 AUGs of ATF4 (Fig. S7A, in 3AUGMut), which would eliminate all ATF4 variants except for the potential frameshifted fusion protein. However, we observed that the peak corresponding to the 53 kDa wt protein did not vanish but shifted to 52 kDa (Fig. S7A) and its relative expression level decreased by ~5.5-fold under both conditions when the 3 AUGs were mutated (Fig. S7A, bottom electropherogram, Table S11 and Table S12. This 52 kDa peak in 3AUGMut remained inducible under Tg stress, indicating that despite the lack of these three ATF4 AUGs, the regulatory system remained fully responsive. Further, we observed two additional, clearly discernible peaks at 44 and 34 kDa, which were consistently also inducible upon stress. Careful inspection of the ATF4 sequence suggested that the 52 kDa peak may correspond to the almost full-length protein initiated either on the CUG or on five other nearby near-cognate codons (Fig. S8A), whereas the 44 and 34 peaks corresponded to shorter ATF4 variants initiated on internal AUGs further downstream.

These results clearly suggest that ATF4 might be translated from several alternative start sites, and the variants are induced by uORF2. However, since none of these peaks occurred at a higher molecular weight than the original ATF4 protein, the idea of frameshifting resulting in a longer ATF4 variant was not supported. Concordantly, shortening the ATF4 sequence by removing 103 amino acid residues from the C-terminal segment of its CDS shifted all three peaks upward, exactly according to the corresponding loss in molecular weight (Fig. S8B – 3AUGMut versus Fig. S8C – 3AUGMut-short). Furthermore, inserting 2 consecutive stops immediately downstream of the uORF2/ATF4 overlap (in the ATF4 reading frame) eliminated only the heaviest peak but not the other two peaks, in both the full-length and shortened 3AUGMut constructs (Fig. S8D – 3AUGMut-stop, Fig. S8E – 3AUGMut-short-stop), strongly suggesting that the initiation codon for the 52-kDa protein lied within the overlap region, but not in the uORF2 frame. Consistently, combining 3AUGMut with an AUG to AGG mutation of uORF2 had no effect on the size and distribution of the aforementioned 3 protein products (Fig. S9A, in d2-3AUGMut). Indeed, inserting 2 consecutive stops into the wt construct at the same place as in case of the 3AUGMut construct completely eliminated the signal (data not shown).

The final piece of evidence for the absence of frameshifting arose from an insertion of a 10x c-Myc tag into the uORF2 frame exactly 21 nt downstream of its AUG (Fig. S7B, in uORF2_ins). This mutant resulted in only one ATF4-HA peak corresponding to ~53 kDa, with inducibility unchanged compared to the wt (Fig. S7B, upper electropherogram). Note that the expected size of the fusion protein is ~62 kDa but it runs at ~57 kDa, as verified experimentally (Fig. S9B, Table S13 and Table S14). Probing with c-Myc antibodies revealed a single peak at ~40 kDa corresponding to the uORF2-c-Myc fusion protein, whose presence was strictly depended on the presence of the AUG of uORF2 (Fig. S7B; bottom panel, uORF2_ins => AUG to AGG in red) and whose intensity remained unexpectedly high (reduced only by ~40%) even under Tg stress (Fig. S7B, bottom electropherogram; see Table S11 and Table S12). Consistently, combining uORF2_ins with the triple AUG mutation of ATF4 produced three Tg-inducible peaks of the same size as observed with the 3AUGMut alone (Fig. S9C [uORF2_ins_3AUGMut] versus Fig. S8B [3AUGMut]). Finally, mutations of the computer-predicted slippery sequences (Fig. 2A) showed no effect (data not shown). Altogether, these findings clearly ruled out possible uORF2 to ATF4 frameshifting and strongly supported the idea that uORF2 was translated even under stress and at a much higher level than could be expected based on the original model.

### SUPPLEMENTAL MATERIALS AND METHODS

#### Construction of bacterial plasmids

List of all plasmids, PCR primers and GeneArt Strings DNA Fragments (Invitrogen) used throughout this study can be found in Tables S24 - S26.

wt ATF4-HA Tag was created by inserting *Hind*III-*Not*I digested h*ATF4*-wt-HA-Tag (GeneArt String DNA Fragment; Invitrogen) into *Hind*III-*Not*I digested *pCMV-EGFP-N2* high copy vector.

dSt-st was created by inserting the *Hind*III-*Pst*I digested fusion PCR product obtained with primers h*ATF4*-St-st_AGG-F and h*ATF4*-St-st_AGG-R using wt ATF4-HA Tag as a template into *Hind*III-*Pst*I digested wt ATF4-HA Tag.

d1 was created by inserting the *Hind*III-*Pst*I digested fusion PCR product obtained with primers h*ATF4*-uORF1_AGG-F and h*ATF4*-uORF1_AGG-R primers using wt ATF4-HA Tag as a template into *Hind*III-*Pst*I digested wt ATF4-HA Tag.

d2 was created by inserting the *Pst*I-*EcoR*V digested fusion PCR product obtained with primers h*ATF4*-uORF2_AGG and SW *ATF4* d120 SphI R using wt ATF4-HA Tag as a template into *Pst*I-*EcoR*V digested wt ATF4-HA Tag.

d-all was created by inserting *Hind*III-*Not*I digested h*ATF4*-d-all-HA-Tag (GeneArt String DNA Fragment; Invitrogen) into *Hind*III-*Not*I digested *pCMV-EGFP-N2* high copy vector.

St-st-only was created using fusion PCR with wt ATF4-HA Tag serving as a template and the following combination of primers: (i) h*ATF4*-uORF1_AGG-F + h*ATF4*-uORF1_AGG-R and (ii) h*ATF4*-uORF2_AGG and SW *ATF4* d120 SphI R. The resulting PCR product was digested with *Hind*III and *EcoR*V and inserted into *Hind*III-*EcoR*V digested wt ATF4-HA Tag.

uORF1-only was created using fusion PCR with wt ATF4-HA Tag serving as a template and the following combination of primers: (i) h*ATF4*-St-st_AGG-F + h*ATF4*-St-st_AGG-R and (ii) h*ATF4*-uORF2_AGG and SW *ATF4* d120 SphI R. The resulting PCR product was digested with *Hind*III and *EcoR*V and inserted into *Hind*III-*EcoR*V digested wt ATF4-HA Tag.

uORF2-only was created using fusion PCR with wt ATF4-HA Tag serving as a template and the following combination of primers: (i) h*ATF4*-St-st_AGG-F + h*ATF4*-St-st_AGG-R and (ii) h*ATF4*-uORF1_AGG-F + h*ATF4*-uORF1_AGG-R. The resulting PCR product was digested with *Hind*III and *Pst*I and inserted into *Hind*III-*Pst*I digested wt ATF4-HA Tag.

dSt-st-NC1 was created by inserting the *Hind*III-*Pst*I digested PCR product obtained with primers h*ATF4*-dSt-st_NC1 and h*ATF4*-PstI-R using dSt-st as a template into *Hind*III-*Pst*I digested dSt-st.

St-st-only-NC1 was created by inserting the *Hind*III-*Pst*I digested PCR product obtained with primers h*ATF4*-St-st-only_NC1 and h*ATF4*-PstI R using St-st-only as a template into *Hind*III-*Pst*I digested St-st-only.

St-st-only-NC2/3 was created by inserting the *Hind*III-*Pst*I digested PCR product obtained with primers h*ATF4*-St-st-only_NC2/3 and h*ATF4*-PstI-R using St-st-only as a template into *Hind*III-*Pst*I digested St-st-only.

dSt-st-SL1 was created by inserting the *Hind*III-*Pst*I digested PCR product obtained with primers h*ATF4*-dSt-st-SL1 and h*ATF4*-PstI-R using dSt-st as a template into *Hind*III-*Pst*I digested dSt-st.

St-st-only-SL2-1 was created by inserting the *Hind*III-*Pst*I digested fusion PCR product obtained with primers h*ATF4*-SL2-1-F and h*ATF4*-SL2-1-R using St-st-only as a template into *Hind*III-*Pst*I digested St-st-only.

St-st-only-SL2-2 was created by inserting the *Hind*III-*Pst*I digested fusion PCR product obtained with primers h*ATF4*-SL2-2-F and h*ATF4*-SL2-2-R using St-st-only as a template into *Hind*III-*Pst*I digested St-st-only.

dSt-st-SL2-1 was created by inserting the *Hind*III-*Pst*I digested fusion PCR product obtained with primers h*ATF4*-SL2-1-F and h*ATF4*-SL2-1-R using dSt-st as a template into *Hind*III-*Pst*I digested dSt-st.

dSt-st-SL2-2 was created by inserting the *Hind*III-*Pst*I digested fusion PCR product obtained with primers h*ATF4*-SL2-2-F and h*ATF4*-SL2-2-R using dSt-st as a template into *Hind*III-*Pst*I digested dSt-st.

d-all-SL2-2 was created by inserting the *Hind*III-*Pst*I digested fusion PCR product obtained with primers h*ATF4*-SL2-2-F and h*ATF4*-SL2-2-R using d-all as a template into *Hind*III-*Pst*I digested d-all.

wt-A235G was created by inserting the *Pst*I-*EcoR*V digested fusion PCR product obtained with primers h*ATF4*-A235G-F and h*ATF4*-A235G-R using wt ATF4-HA Tag as a template into *Pst*I-*EcoR*V digested wt ATF4-HA Tag.

wt-SL3^Mut-1^ was created by inserting the *Pst*I-*EcoR*V digested fusion PCR product obtained with primers h*ATF4*-SL3_CAAAA-F and h*ATF4*-SL3_CAAAA-R using wt ATF4-HA Tag as a template into *Pst*I-*EcoR*V digested wt ATF4-HA Tag.

wt-SL3^Mut-2^ was created by *Pst*I-*EcoR*V digested h*ATF4*-wt-SL3^Mut-2^ (GeneArt String DNA Fragment; Invitrogen) into *Pst*I-*EcoR*V digested wt ATF4-HA Tag.

dSt-st-SL3^Mut-1^ was created by inserting the *Pst*I-*EcoR*V digested fusion PCR product obtained with primers h*ATF4*-SL3_CAAAA-F and h*ATF4*-SL3_CAAAA-R using dSt-st as a template into *Pst*I-*EcoR*V digested dSt-st.

d-all-SL3^Mut-1^ was created by inserting the *Pst*I-*EcoR*V digested fusion PCR product obtained with primers h*ATF4*-SL3_CAAAA-F and h*ATF4*-SL3_CAAAA-R using d-all as a template into *Pst*I-*EcoR*V digested d-all.

uORF2-only-SL3^Mut-1^ was created by inserting the *Pst*I-*EcoR*V digested fusion PCR product obtained with primers h*ATF4*-SL3_CAAAA-F and h*ATF4*-SL3_CAAAA-R using uORF2-only as a template into *Pst*I-*EcoR*V digested uORF2-only.

wt-AUG3^Mut^ (3rd *ATF4* AUG) was created by inserting the *Pst*I-*EcoR*V digested fusion PCR product obtained with primers h*ATF4*-3rdAGG-F and h*ATF4*-3rdAGG-R wt ATF4-HA Tag as a template into *Pst*I-*EcoR*V digested wt ATF4-HA Tag.

dSt-st-AUG3^Mut^ (3rd *ATF4* AUG) was created by inserting the *Pst*I-*EcoR*V digested fusion PCR product obtained with primers h*ATF4*-3rdAGG-F and h*ATF4*-3rdAGG-R dSt-st as a template into *Pst*I-*EcoR*V digested dSt-st.

wt-UUG^Mut^-AUG3 was created by inserting *Pst*I-*EcoR*V digested h*ATF4*-wt-UUG^Mut^_AUG3 (GeneArt String DNA Fragment; Invitrogen) into *Pst*I-*EcoR*V digested wt ATF4-HA Tag.

wt-SL3^Mut-1^-AUG3^Mut^ was created by inserting *Pst*I-*EcoR*V digested h*ATF4*-wt-SL3^Mut-1^-AUG3^Mut^ (GeneArt String DNA Fragment; Invitrogen) into *Pst*I-*EcoR*V digested wt ATF4-HA Tag.

wt-A326G was created by inserting *Pst*I-*EcoR*V digested h*ATF4*-wt-A326G (GeneArt String DNA Fragment; Invitrogen) into *Pst*I-*EcoR*V digested wt ATF4-HA Tag.

wt-SL3^Mut-1^-A326G was created by *Pst*I-*EcoR*V digested h*ATF4*-wt-SL3^Mut-1^-A326G (GeneArt String DNA Fragment; Invitrogen) into *Pst*I-*EcoR*V digested wt ATF4-HA Tag.

wt-3AUG^Mut^ was created by inserting *Pst*I-*EcoR*V digested h*ATF4*-wt-3AUG^Mut^ of *ATF4* (GeneArt String DNA Fragment; Invitrogen) into *Pst*I-*EcoR*V digested wt ATF4-HA Tag.

d2-3AUG^Mut^ was created by inserting *Pst*I-*EcoR*V digested h*ATF4*-d2-3AUG^Mut^ of *ATF4* (GeneArt String DNA Fragment; Invitrogen) into *Pst*I-*EcoR*V digested wt ATF4-HA Tag.

3AUG^Mut^-stop was created by inserting *Pst*I-*EcoR*V digested h*ATF4*-3AUG^Mut^-stop (GeneArt String DNA Fragment; Invitrogen) into *Pst*I-*EcoR*V digested wt ATF4-HA Tag.

3AUG^Mut^-short was created by inserting *EcoR*V-*Not*I digested h*ATF4*-3AUG^Mut^-short (GeneArt String DNA Fragment; Invitrogen) into *EcoR*V-*Not*I digested wt-3AUG^Mut^.

3AUG^Mut^-short-stop was created by inserting *EcoR*V-*Not*I digested h*ATF4*-3AUG^Mut^-short (GeneArt String DNA Fragment; Invitrogen) into *EcoR*V-*Not*I digested 3AUG^Mut^-stop.

uORF2_ins was created by inserting *Pst*I-*EcoR*V digested h*ATF4*-uORF2_ins (GeneArt String DNA Fragment; Invitrogen) into PstI-EcoRV digested wt ATF4-HA Tag.

d2_ins (uORF2_ins => AUG to AGG) was created by inserting *Pst*I-*EcoR*V digested h*ATF4*-d2_ins (GeneArt String DNA Fragment; Invitrogen) into *Pst*I-*EcoR*V digested wt ATF4-HA Tag.

uORF2_ins_3AUG^Mut^ was created by inserting *Pst*I-*EcoR*V digested h*ATF4*-uORF2_ins_3AUG^Mut^ (GeneArt String DNA Fragment; Invitrogen) into *Pst*I-*EcoR*V digested wt ATF4-HA Tag.

wt-CUG^Mut^ was created by inserting *Pst*I-*EcoR*V digested h*ATF4*-wt-CUG^Mut^ (GeneArt String DNA Fragment; Invitrogen) into *Pst*I-*EcoR*V digested wt ATF4-HA Tag.

wt-SL3^Mut-1^-CUG^Mut^ was created by inserting *Pst*I-*EcoR*V digested h*ATF4*-wt— SL3^Mut-1^-CUG^Mut^ (GeneArt String DNA Fragment; Invitrogen) into *Pst*I-*EcoR*V digested wt ATF4-HA Tag.

wt_ins was created by inserting *Pst*I-*EcoR*V digested h*ATF4*-wt_ins (GeneArt String DNA Fragment; Invitrogen) into *Pst*I-*EcoR*V digested wt ATF4-HA Tag.

wt_ins-SL3^Mut-1^ was created by inserting *Pst*I-*EcoR*V digested h*ATF4*-wt_ins-SL3^Mut-1^ (GeneArt String DNA Fragment; Invitrogen) into *Pst*I-*EcoR*V digested wt ATF4-HA Tag.

d2_ins was created by inserting *BstB*I digested d2 into *BstB*I digested wt_ins. d2_ins-SL3^Mut-1^ was created by inserting *BstB*I digested d2 into *BstB*I digested wt_ins-SL3^Mut-1^.

uORF2_ins-SL3^Mut-1^ was created by inserting *BstB*I digested uORF2_ins into *BstB*I digested wt-SL3^Mut-1^.

uORF2-*ATF4*-HA fusion was created by inserting *Pst*I-*EcoR*V digested h*ATF4*-uORF2-*ATF4*-HA fusion (GeneArt String DNA Fragment; Invitrogen) into *Pst*I-*EcoR*V digested wt ATF4-HA Tag.

### SUPPLEMENTARY FIGURES AND FIGURE LEGENDS

**Figure S1.**
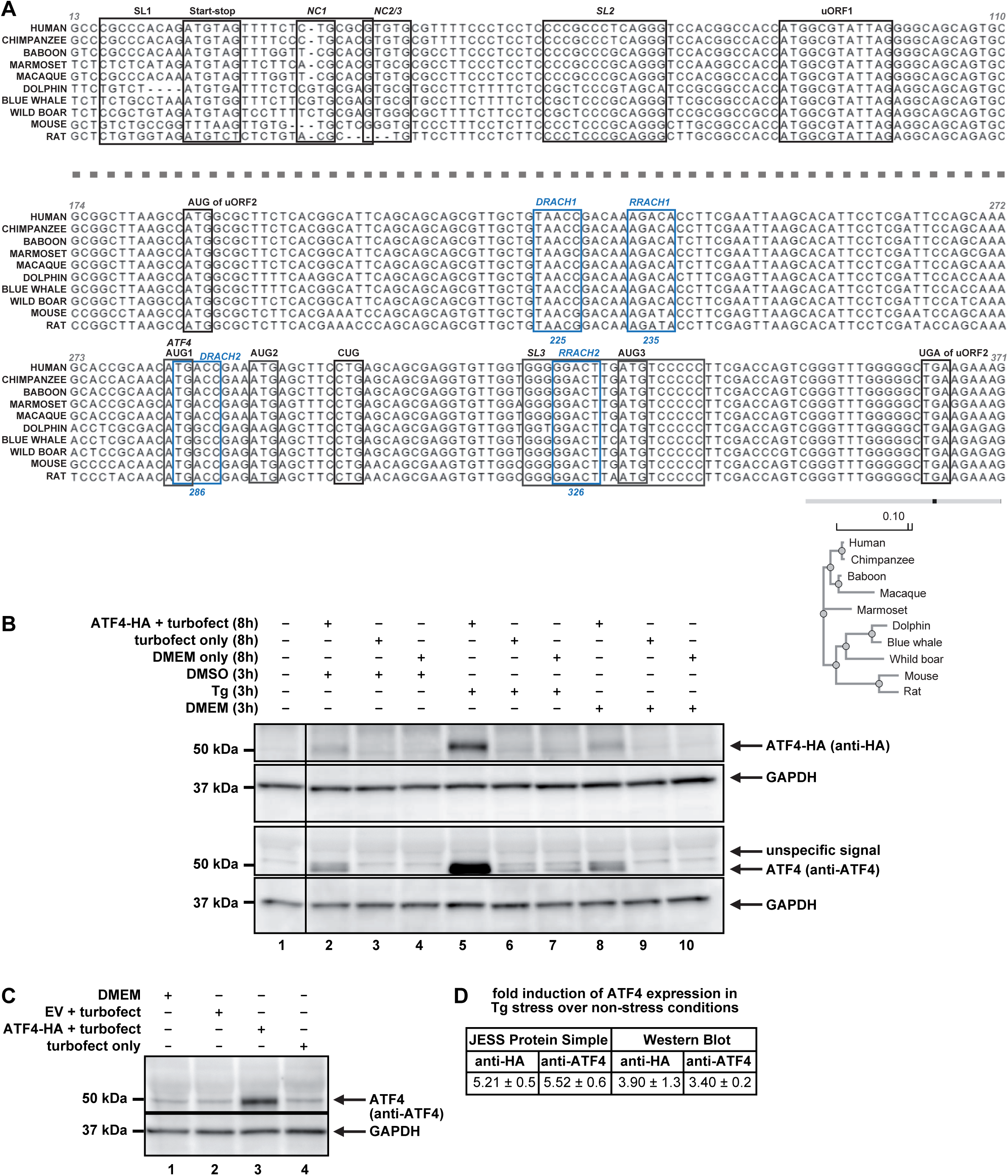
The SL3 and CUG are conserved in evolution. (A) Multiple Sequence Alignment of 10 mammalian species using MAFFT (Multiple Alignment using Fast Fourier Transform) is shown for the 5′ UTR of ATF4 mRNA beginning 12 nucleotides upstream of the AUG of Start-stop and ending six nucleotides past the stop codon of uORF2. The ATF4‘s mRNA-specific features under study and their conservation comparisons are outlined in bold boxes. Genomic alignments of mammalian species were compared using Job Dispatcher EMBL-EBI website MAFFT tool and Madeira et al. (Madeira et al., 2022). The Phylogram based on Phylogenetic Tree scores reflects sequence differences between the *ATF4* gene among given species. (B) Our ATF4-HA reporter system faithfully recapitulates the endogenous *ATF4* regulation as demonstrated by traditional western blot analysis. See SI for further details. Results are representative of three independent experiments. (C) Transfection of an empty vector (EV) did not increase ATF4 expression under non-stress conditions as demonstrated by traditional western blot analysis. See text for further details. Results are representative of three independent experiments. (D) Fold induction of ATF4 expression in Tg stress *versus* non-stress conditions is comparable over different methods and antibodies. All quantifications were done from a minimum of three independent experiments (JESS anti-HA n=17; JESS anti-ATF4 n=3; WB anti-HA n=3; WB anti-ATF4 n=4). For western blots, only exposures with non-saturated signals were used for quantifications by Quantity One software.

**Figure S2.**
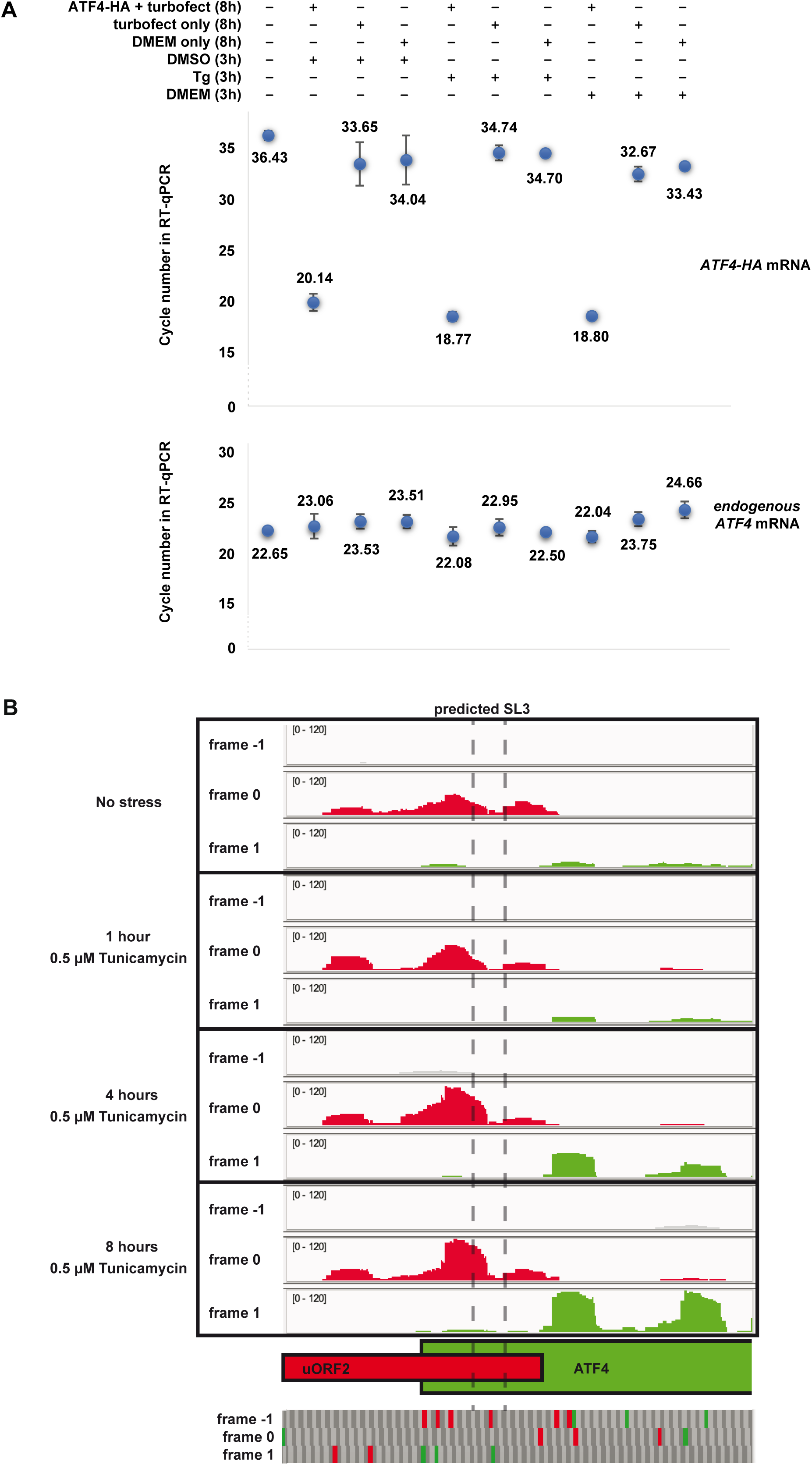
Our ATF4-HA reporter system faithfully recapitulates the endogenous *ATF4* regulation; does uORF2 to *ATF4* frameshifting occurs during acute stress? (A) mRNA levels of both *ATF4-HA* and endogenous *ATF4* (measured separately using highly specific reverse primers matching the *ATF4* stop codon region, by which these two alleles differ due to the HA tag sequence) remain virtually unchanged under non-stress *vs.* stress conditions. See text for further details. (B) Ribosome footprints from HeLa cells undergoing acute ER stress (Rendleman et al., 2018) were mapped to the *ATF4* exon 2 based on the reading frame engaged. Reading frame assignment was determined by the +12 position in reads 28-30 bp long, corresponding to the P-site of ribosomes. ER stress was induced by 0.5 µM Tunicamycin and cells were harvested at 0, 1, 4, and 8 hours post-treatment, as described in (Rendleman et al., 2018). Predicted stem-loop 3 (SL3) start and end positions are indicated by the dashed line. Below, canonical start and stop codons within frames −1, 0, and 1 are indicated in green and red, respectively.

**Figure S3.**
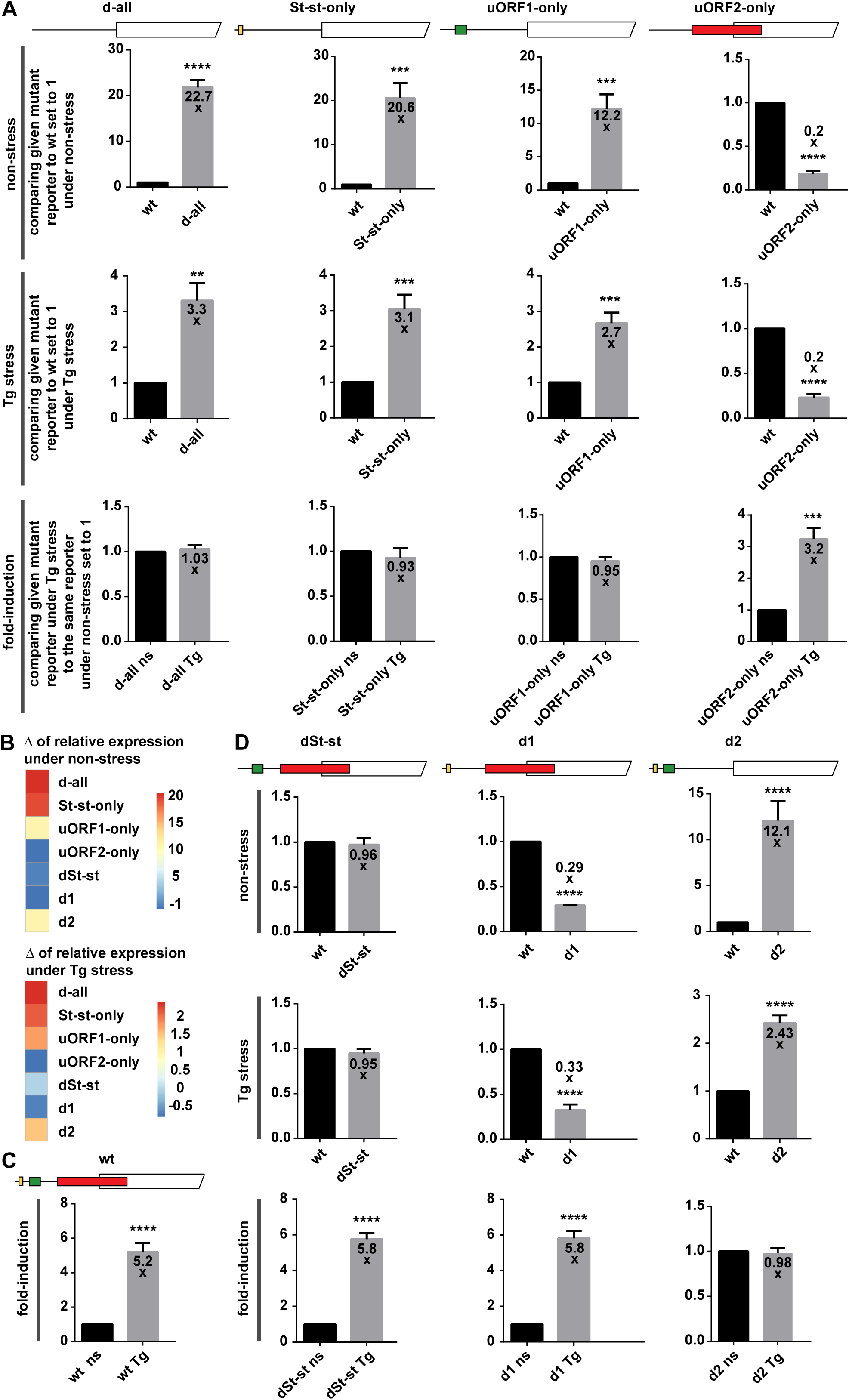
Revisiting the delayed translation reinitiation mechanism of *ATF4* translational control – uORF2 shows an inducible nature under Tg stress. (A) Schematics at the top of the corresponding panels indicate the *ATF4* mutant constructs subjected to JESS analyses. Relative ATF4-HA protein expression levels were plotted as ratios of values obtained with mutants *versus* wt set to 1 under “non-stress” and “Tg stress” conditions; “fold-induction” plots depict ratios of Tg stress *versus* non-stress values obtained with mutant constructs. Statistical analyses were carried out as in Figure 1E. (B) Heatmaps, created in R (ver. 4.2.2) using the ‘pheatmap’ package (ver. 1.0.12) (Kolde and Kolde, 2015), show the differences (Δ) in the relative ATF4-HA protein expression of individual mutant *versus* wt constructs that were obtained under non-stress (top panel) and Tg stress conditions (bottom panel). The maximal expression levels are shown in red (for the control d-all construct), whereas the minimal levels are depicted in blue (for the uORF2-only construct). (C) Same as Figure 1E for better comparison. (D) Same as panel A with different constructs, depicted at the top, under study.

**Figure S4.**
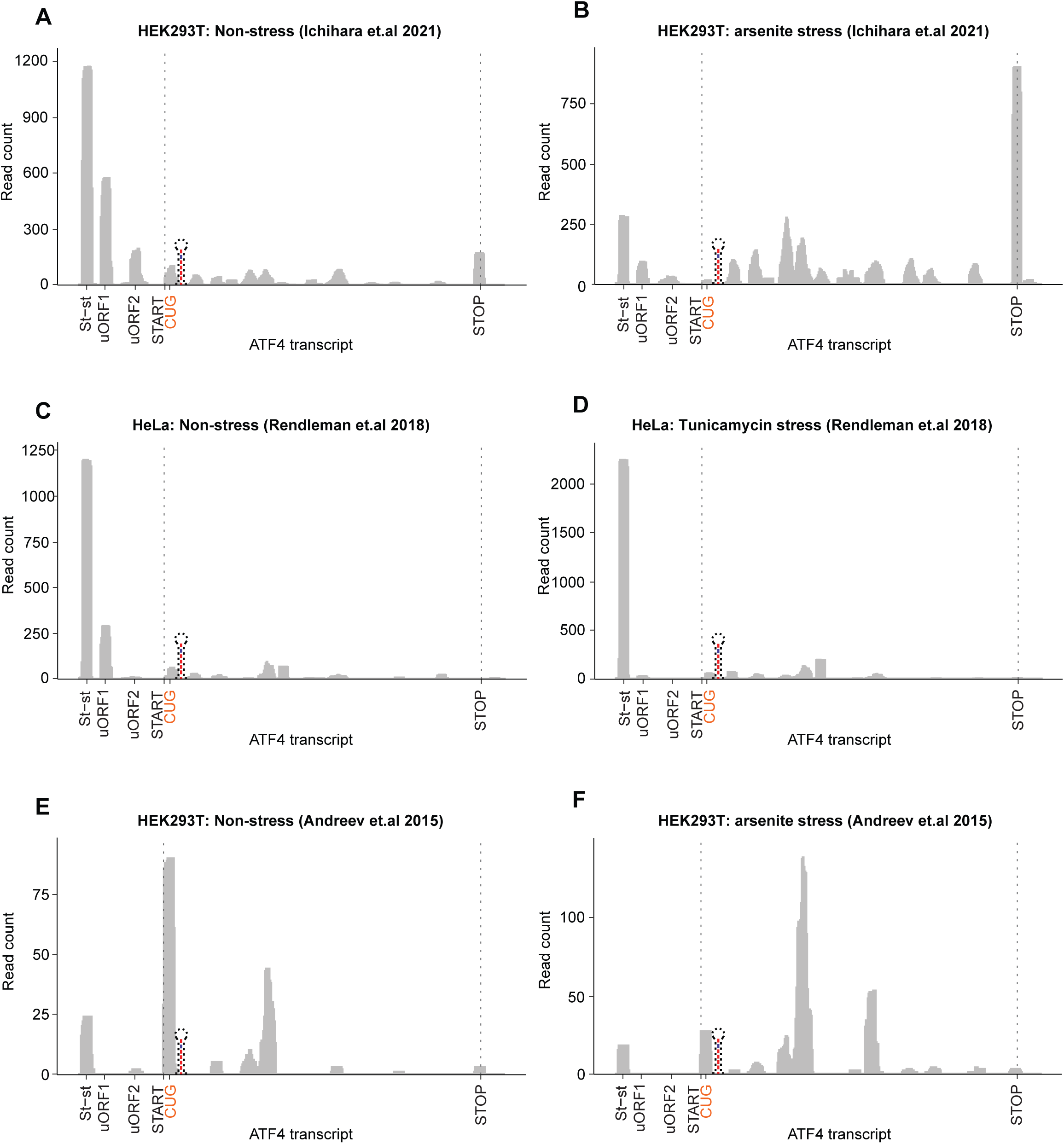
SL3 stalls the ribosome in the uORF2/*ATF4* overlap region. The profiles show ribosome foot-print coverage on the *ATF4* mRNA generated by RiboWaltz (Lauria et al., 2018) package using the datasets obtained from three different studies examining HEK293T cells treated with DMSO (A and E) or Sodium arsenite (B and F) or HeLa cells treated with tunicamycin for 0 (C) or 4 h (D) (Andreev et al., 2015; Ichihara et al., 2021; Rendleman et al., 2018). The *ATF4* transcript with its *cis*-acting features indicated is plotted on the X-axis. Raw read count is plotted on the Y-axis. The dotted lines indicate the 1st nucleotide of canonical start and stop codons of the *ATF4* ORF; the near cognate “CUG” codon is highlighted in red. The schematic stem-loop indicates the location of the predicted RNA secondary structure designed SL3.

**Figure S5.**
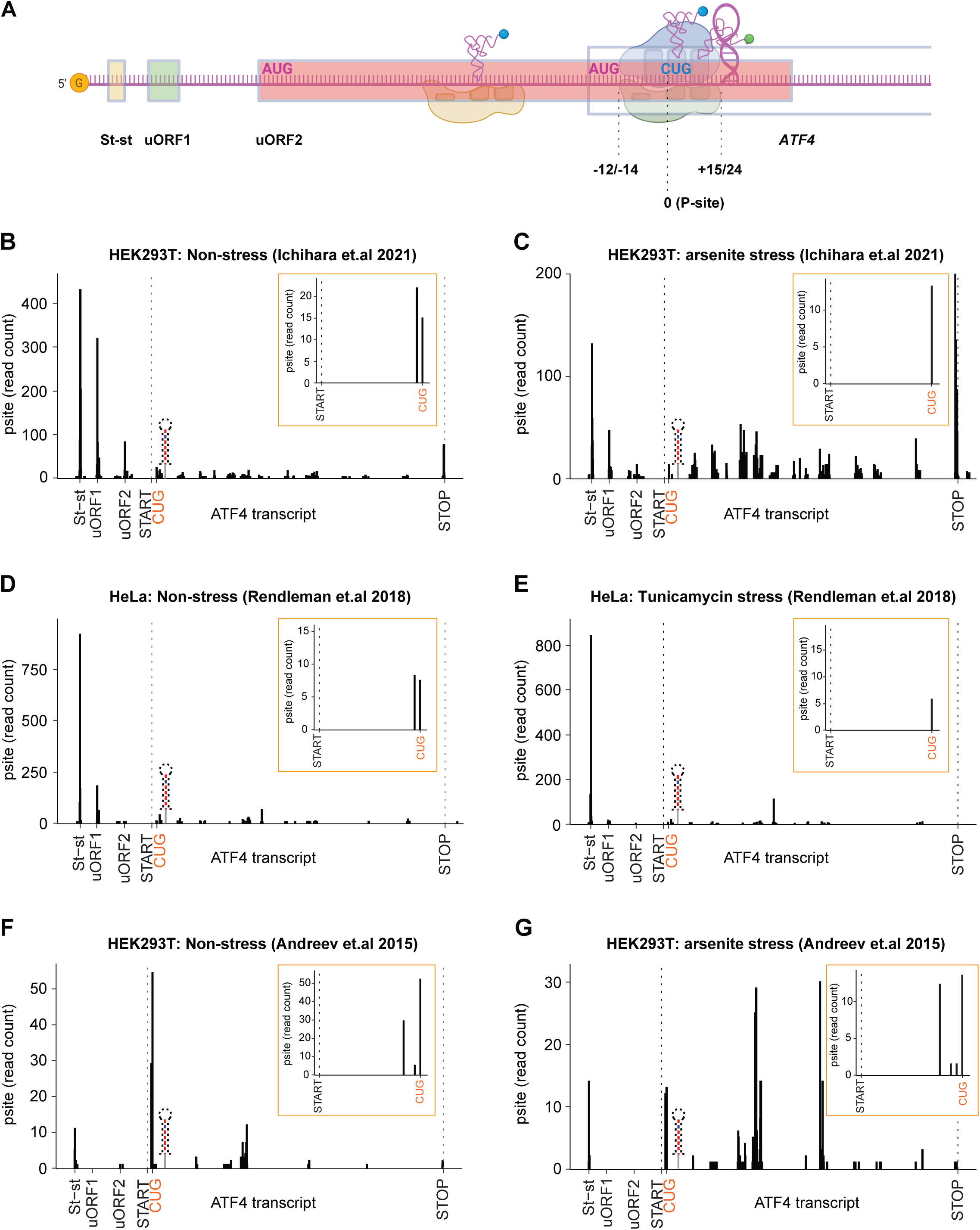
SL3 may promote placement of the near-cognate “CUG” in the ribosomal P-site. (A) Schematic illustrating that the ribosome paused by SL3 on *ATF4* mRNA is in an ideal distance to position the near-cognate CUG in its P site to initiate translation. (B – G) The profiles display the P-site occupancy of Ribosome Protected Fragments (RPFs) on the *ATF4* mRNA generated by RiboWaltz (Lauria et al., 2018) package using the datasets obtained from three different studies examining HEK293T cells treated with DMSO (B and F) or Sodium arsenite (C and G) or HeLa cells treated with tunicamycin for 0 (D) or 4 h (E) (Andreev et al., 2015; Ichihara et al., 2021; Rendleman et al., 2018). The corresponding zoomed-in views of the P-site occupancy at the near-cognate “CUG” codon are shown in orangish boxes. The *ATF4* transcript with its *cis*-acting features indicated is plotted on the X-axis. Raw read count is plotted on the Y-axis. The dotted lines indicate the 1st nucleotide of canonical start and stop codons of the *ATF4* ORF; the near cognate “CUG” codon is highlighted in red. The schematic of the stem-loop indicates the location of the predicted RNA secondary structure designated as SL3.

**Figure S6.**
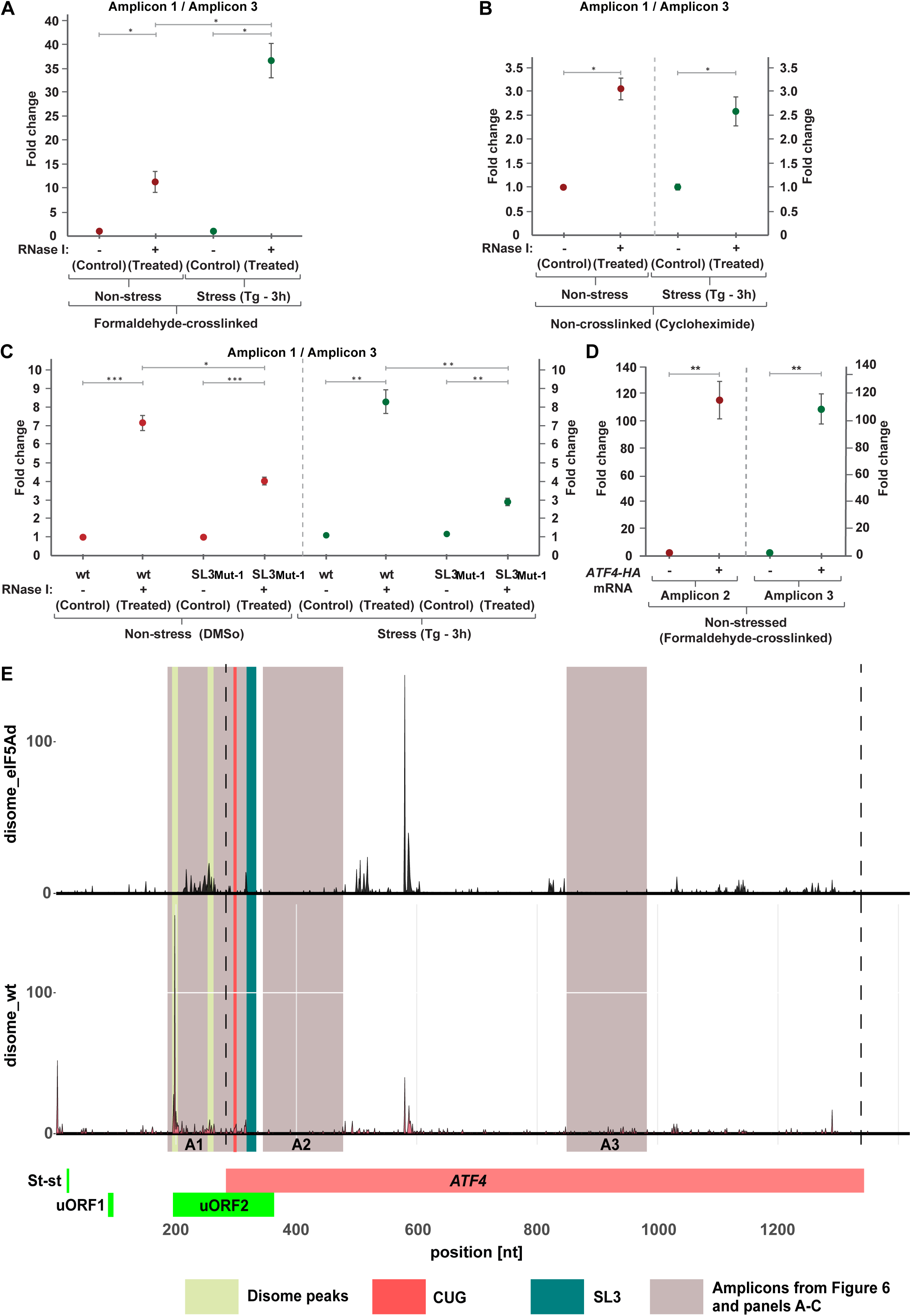
Further evidence supporting ribosome queuing hypothesis. (A) HEK293T cells were cross-linked with formaldehyde (HCHO) and then subjected to the ribosome-protection assay as described in panel B of Figure 7. qPCR product levels of the recovered putative queuing region (amplicon A1) are normalized to the region in the middle of the *ATF4* CDS (amplicon A3), as well as to the internal RNA isolation control (SPIKE) with non-stress values set to 1 (*p<0.01). Results are representative of three independent experiments. (B) HEK293T cells were treated with cycloheximide (non-crosslinking agent) then subjected to ribosome-protection assay as described in panel C of Figure 7. Results from three independent experiments were analyzed as described in panel B of Figure 7 with non-stress values set to 1 (*p<0.01). (C) HEK293T cells were transiently transfected with plasmids carrying either wt or SL3-mutated (in SL3^Mut-1^) *ATF4* reporters and treated as described in panel B of Figure 7. Results three independent experiments were analyzed as described in panel B of Figure 7 with the wt values set to 1 (*p<0.01, **p<0.001, ***p<0.0001). (D) HEK293T cells were either not transfected or transiently transfected with wt *ATF4* reporter, treated with DMSO (control) for 3 h and total RNA was isolated. qPCR product levels of the *ATF4* mRNA obtained from Amplicon 2 (left panel) or Amplicon 3 (right panel) were normalized to the internal RNA isolation control (SPIKE) and compared between non-transfected and transiently transfected cell with non-transfection values set to 1 (*p<0.01). Results are representative of three independent experiments. (E) Disome-seq coverage along the *ATF4* locus analyzed by RiboCrypt tool (https://ribocrypt.org). Mapped reads were reduced to the 5‘ end of the reads to increase resolution and facilitate identification of the individual peaks. Top track: eIF5A depletion; bottom track: wild type. “Disome peaks” indicates the 5’ ends of disomes; “CUG” and “SL3” indicate their respective positions; “Amplicons” depict the regions subjected to RT-qPCR in Figure 6 and panels A – C.

**Figure S7.**
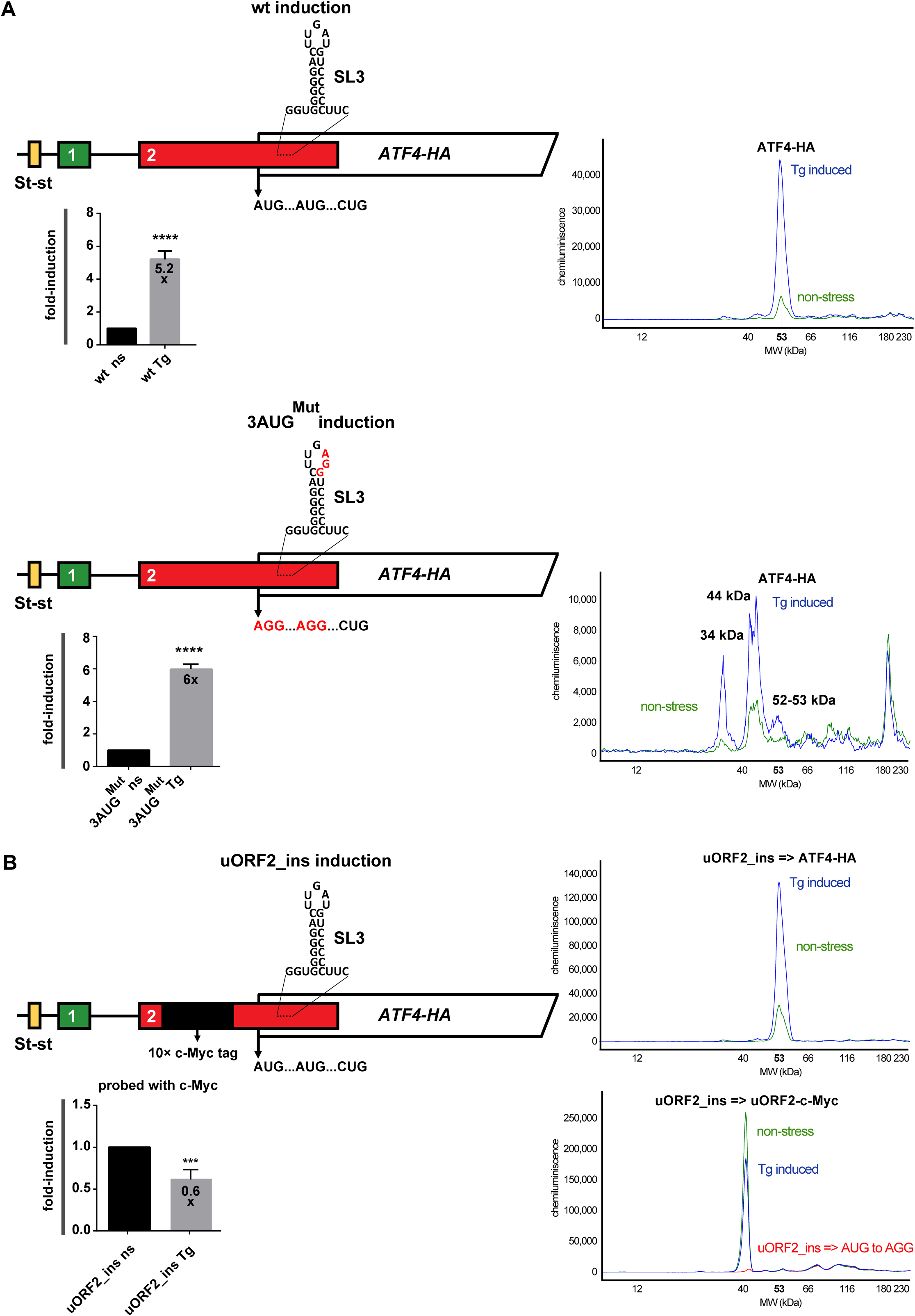
The *ATF4* mRNA expression is not subject to frameshifting; uORF2 is translated even under stress. (A) Same as Figure 1E for better comparison except that the 3AUG^Mut^ construct (bottom panel; depicted at its the top) was also subjected to JESS analyses. The electropherograms of the wt (top panel) and the 3AUG^Mut^ mutant construct (bottom panel) under Tg stress (in blue) compared to non-stress conditions (in green) probed with anti-HA antibodies are shown. For details, see the main text. (B) Same as in Figure S3A except that the 10x c-Myc tag insertion in-frame with uORF2, depicted at the top of the panel, was subjected to JESS analyses. The level of uORF2 induction under stress determined by anti-c-Myc antibodies is plotted. The electropherograms of the construct bearing the 10x c-Myc tag insertion in-frame with uORF2 under Tg stress (in blue) compared to non-stress conditions (in green) probed with the anti-HA (top panel) and anti-c-Myc (bottom panel) antibodies are shown. The electropherogram of the control construct bearing the 10x c-Myc tag insertion in-frame with uORF2, the AUG of which was mutated to AGG, is shown in red. For details, see the main text.

**Figure S8.**
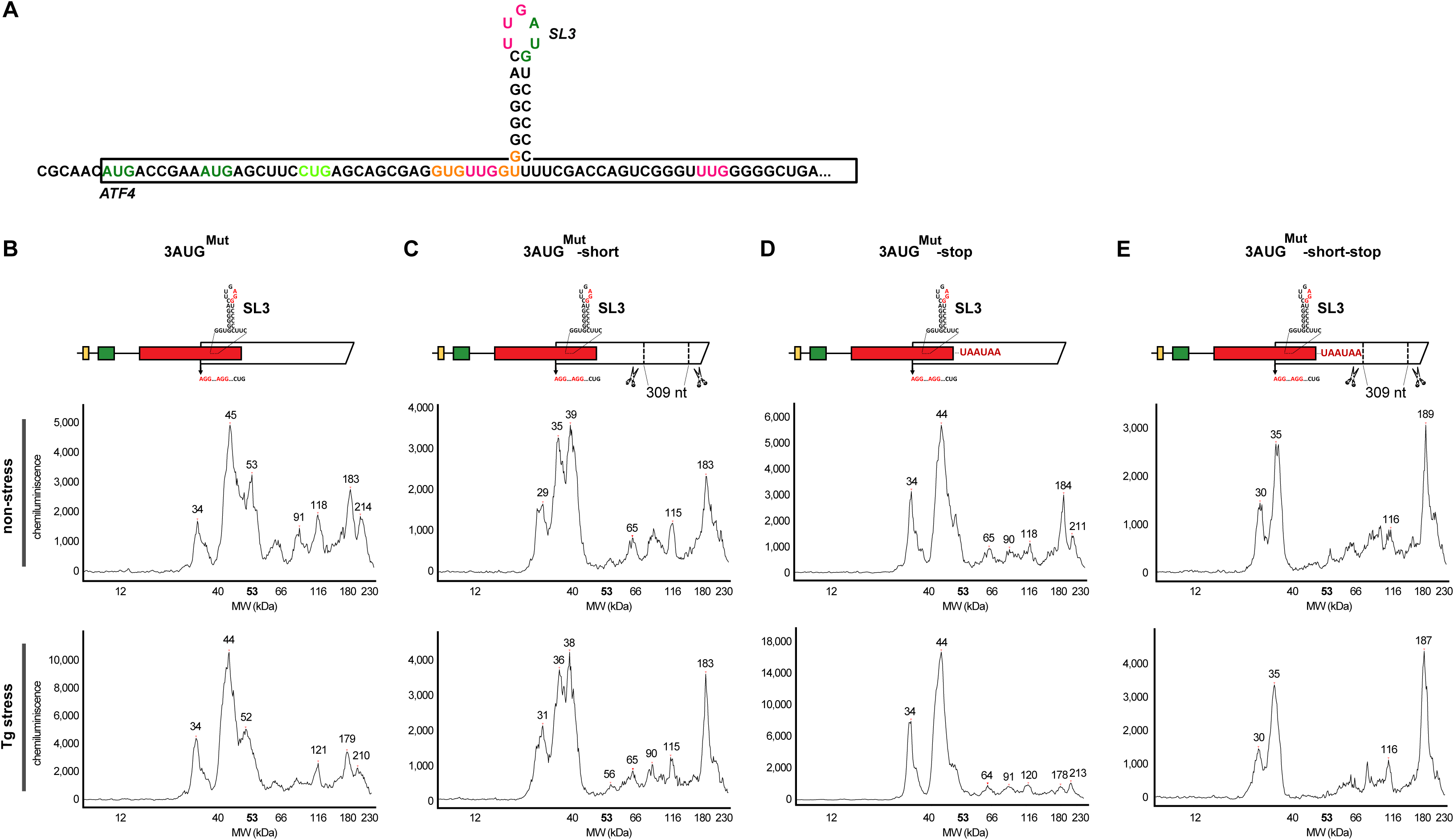
The *ATF4* mRNA expression is not subject to frameshifting; the canonical AUG1 translation start site of ATF4 is substantial leaky scanned. (A) Schematic representation of the *ATF4* sequence showing the annotated AUG translation start site (TSS) of the ATF4 full-length protein, two alternative canonical TSSs, near-cognate CUG and five other near-cognate codons in uORF2/*ATF4* overlap. (B – E) The electropherograms of the full length (B) or C-terminally shortened (C) *ATF4* constructs bearing 3AUG^Mut^ without (B – C) or with 2 consecutive UAA stops inserted immediately downstream of the uORF2/*ATF4* overlap (D – E), depicted at the top of the corresponding panels, are shown.

**Figure S9.**
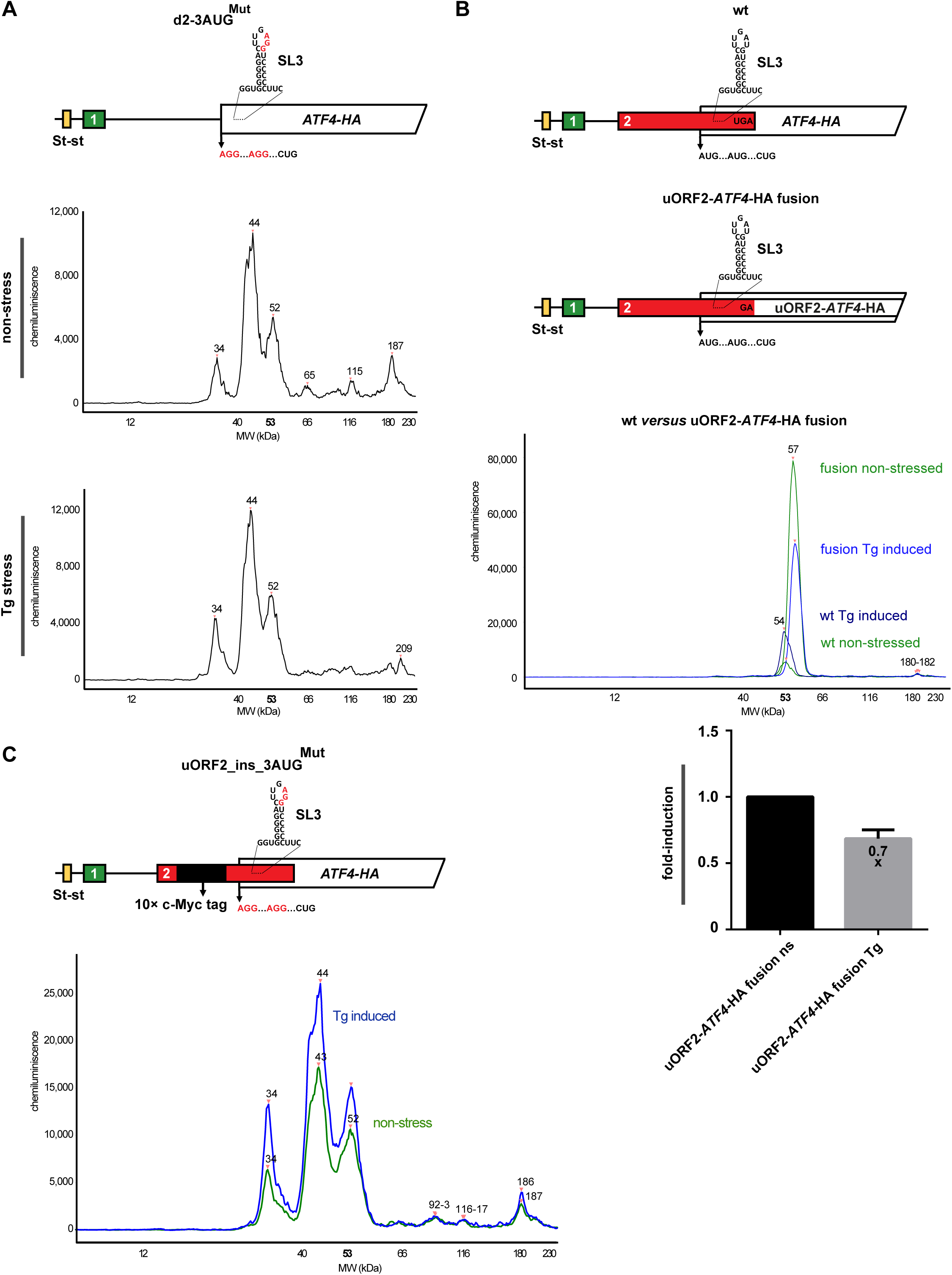
Supporting mutational analysis of the *ATF4* mRNA leader. (A) Combining 3AUG^Mut^ with the AUG to AGG mutation of uORF2 shows no impact on the size and distribution of the ATF4 protein variants expressed with 3AUG^Mut^ alone. The electropherogram of the 3AUG^Mut^ mutation combined with the AUG to AGG mutation of uORF2 in the same *ATF4* construct, depicted at the top, is shown. (B) The artificially created uORF2-*ATF4*-HA fusion product further confirms sustained uORF2 expression even under stress conditions. Schematics at the top depict wt and engineered uORF2-*ATF4*-HA fusion constructs with the electropherograms indicating their expression under non-stress (shades of green) *versus* stress (shades of blue) conditions. Fold-induction values of the fusion construct were plotted (bottom panel) further confirming uORF2 expression even under stress conditions. (C) Further evidence that the *ATF4* mRNA expression is not subject to frameshifting and that uORF2 is expressed even under stress conditions. The schematic at the top depicts the *ATF4*-HA mutant construct where 3AUG^Mut^ was combined with 10x c-Myc tag insertion in-frame with uORF2 with the electropherograms indicating their expression under non-stress (green) *versus* stress (blue) conditions.

**Figure S10.**
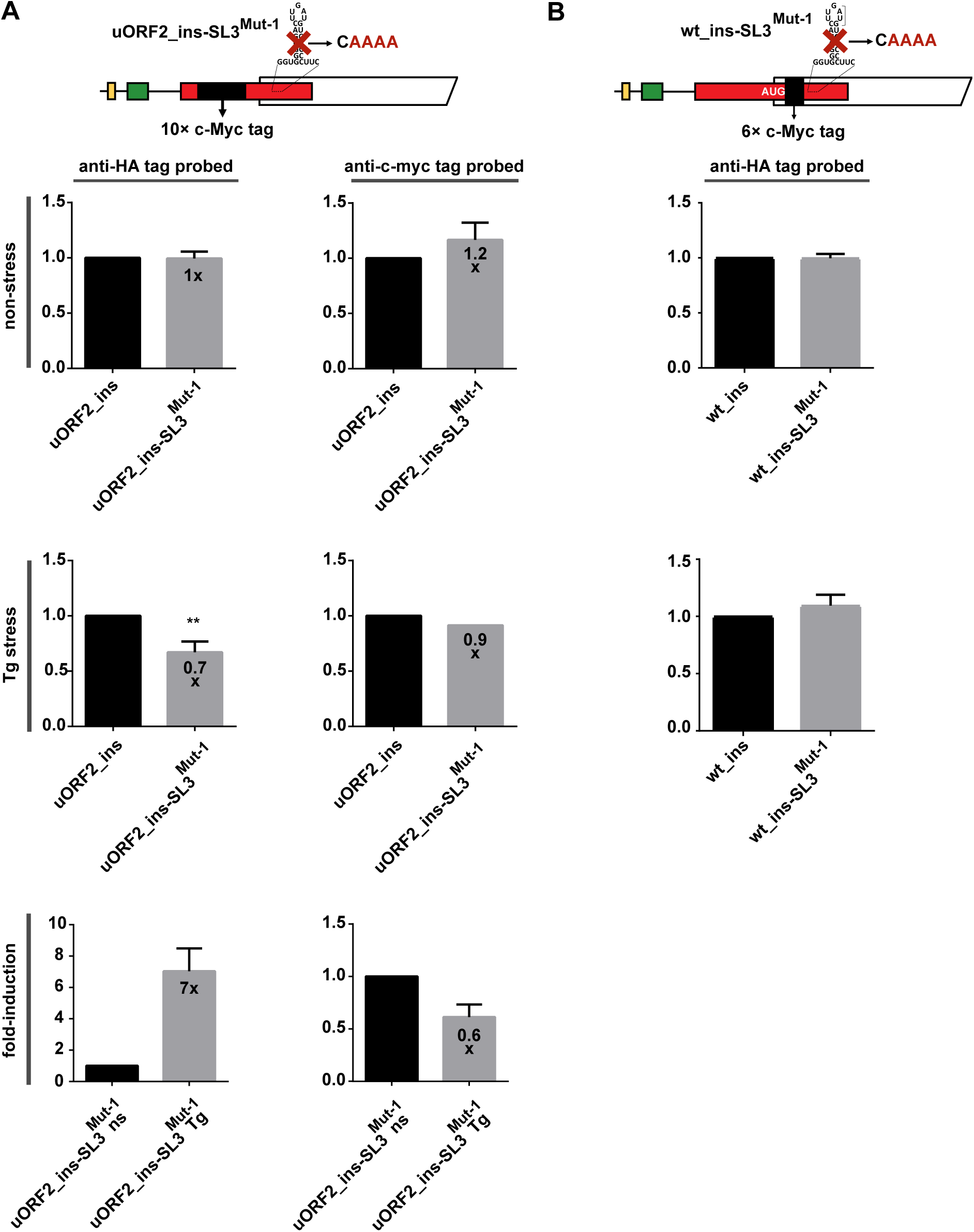
Precise placement of SL3 inside of uORF2 with an exactly defined length expands the existing delayed REI model to a more complex model that includes ribosome queuing. (A) The 10x c-Myc tag insertion in-frame with uORF2 was combined with the SL3^Mut-1^ mutation, as depicted at the top, and subjected to JESS analyses as described in Figure 1E. Probing with both anti-HA (left plots) and anti-c-Myc (right plots) antibodies are shown. (B) The 6x c-Myc tag insertion in-frame with ATF4 was combined with the SL3^Mut-1^ mutation, as depicted at the top, and subjected to JESS analyses as described in Figure 1E.

### SUPPLEMENTAL TABLES

**Table S1.**
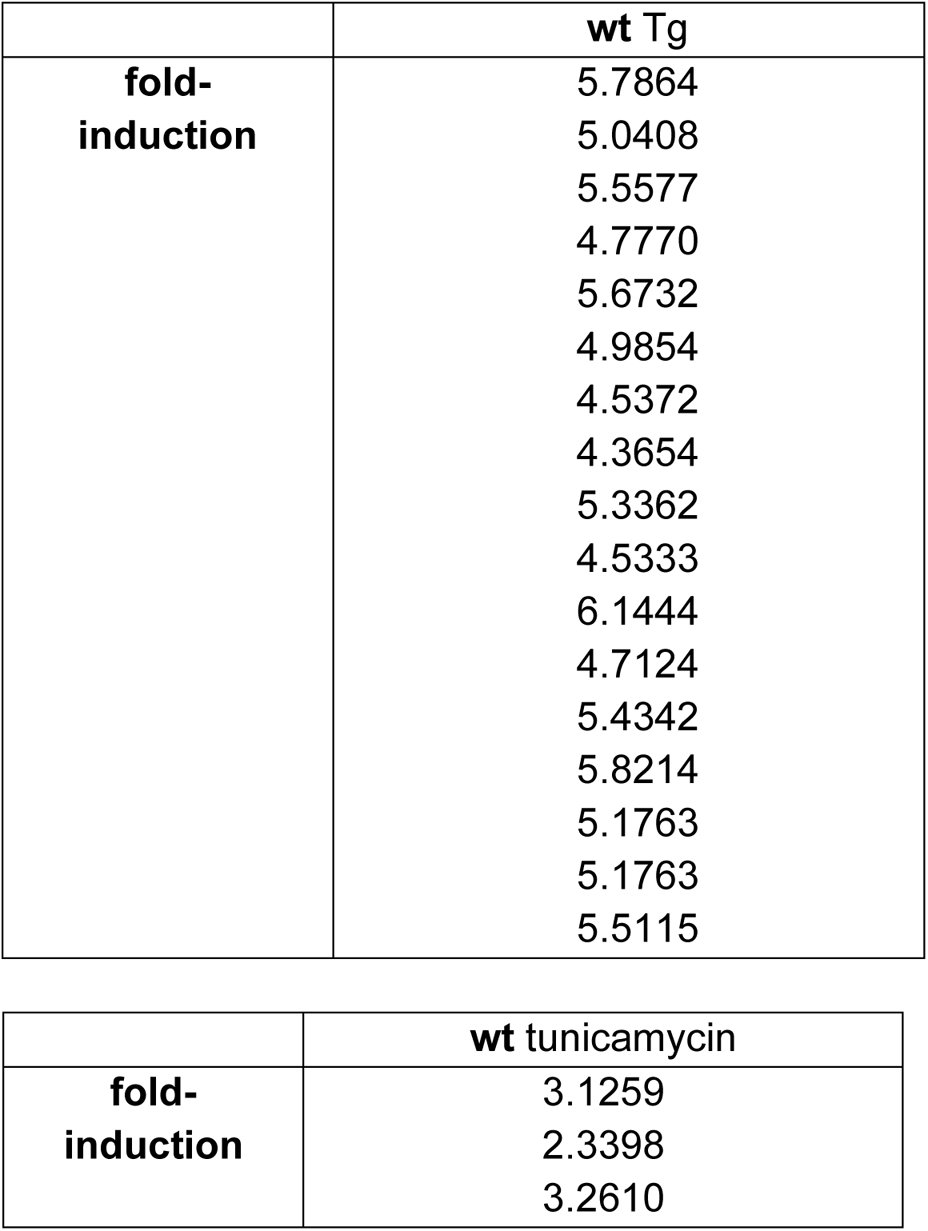
Reference for Figure 1E and 1F; relative wt ATF4-HA protein expression under 3 hours of thapsigargin (Tg) or 4 hours of tunicamycin stress compared to wt ATF4-HA under non-stress (ns) set to 1.

**Table S2.**
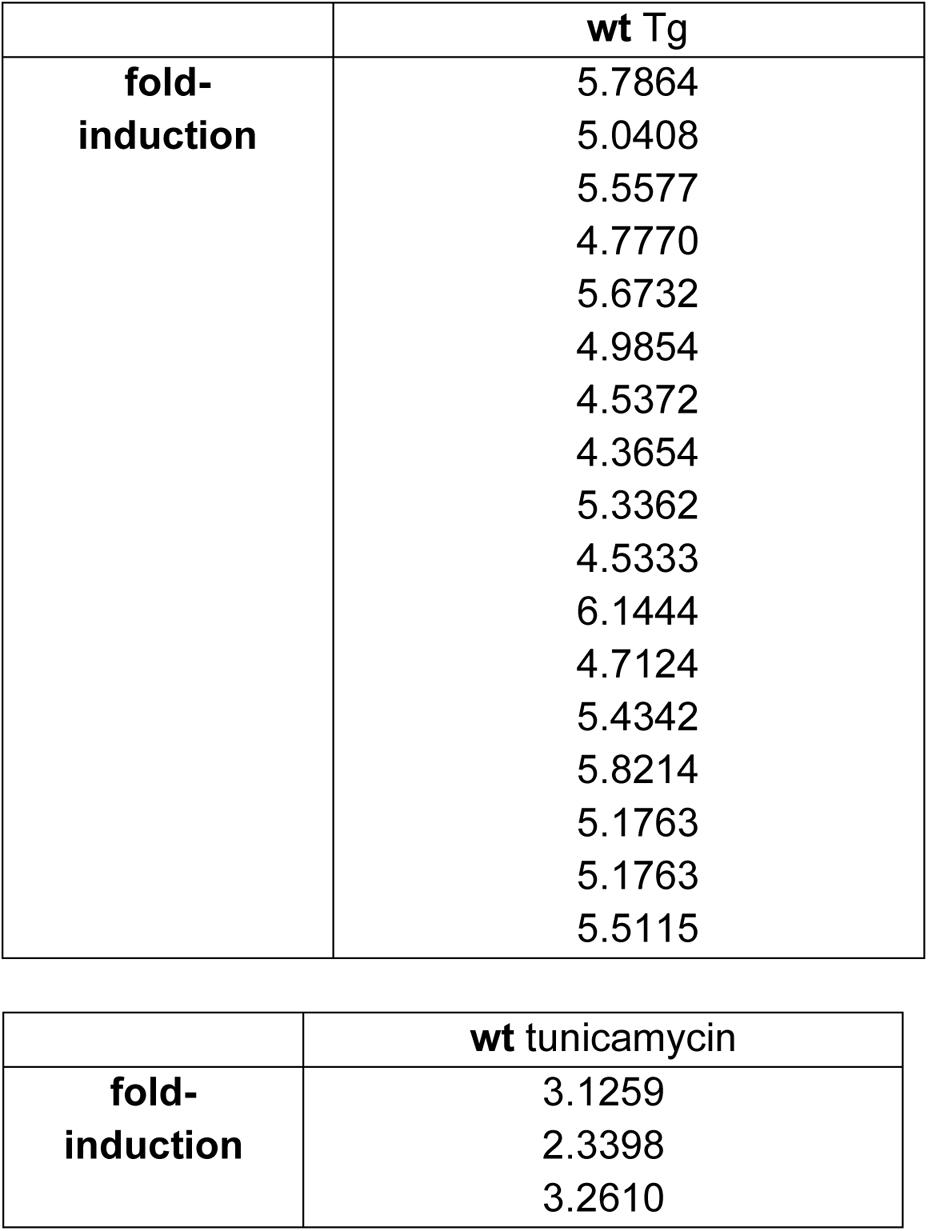
Reference for Figure 1E and 1F and Table S1; relative wt ATF4-HA protein expression under 3 hours of thapsigargin (Tg) or 4 hours of tunicamycin stress compared to wt ATF4-HA under non-stress (ns) set to 1 – individual values.

**Table S3.** Reference for Figure S3; relative ATF4-HA protein expression with wt set to 1. For wt ATF4-HA tag fold-induction see Table S1.

|  | **d-all (N, p-value)** |
| --- | --- |
| **non-stress** | 22.68 ± 2.45 (5, <0.0001 ****) |
| **Tg stress** | 3.31 ± 0.49 (3, 0.0012**) |
| **fold-induction** | 1.03 ± 0.04 (3, 0.3357 ns) |
|  | **St-st-only (N, p-value)** |
| **non-stress** | 20.56 ± 3.43 (3, 0.0006 ***) |
| **Tg stress** | 3.05 ± 0.41 (3, 0.0010 ***) |
| **fold-induction** | 0.93 ± 0.11 (3, 0.3064 ns) |
|  | **u1-only (N, p-value)** |
| **non-stress** | 12.22 ± 2.17 (3, 0.0009 ***) |
| **Tg stress** | 2.67 ± 0.29 (3, 0.0006 ***) |
| **fold-induction** | 0.95 ± 0.04 (3, 0.1438 ns) |
|  | **u2-only (N, p-value)** |
| **non-stress** | 0.19 ± 0.03 (4, <0.0001 ****) |
| **Tg stress** | 0.23 ± 0.04 (3, <0.0001 ****) |
| **fold-induction** | 3.24 ± 0.34 (3, 0.0003 ***) |
|  | **dSt-st (N, p-value)** |
| **non-stress** | 0.96 ± 0.11 (6, 0.3801 ns) |
| **Tg stress** | 0.95 ± 0.05 (3, 0.1221 ns) |
| **fold-induction** | 5.77 ± 0.33 (3, <0.0001 ****) |
|  | **d1 (N, p-value)** |
| **non-stress** | 0.29 ± 0.01 (3, <0.0001 ****) |
| **Tg stress** | 0.33 ± 0.06 (3, <0.0001 ****) |
| **fold-induction** | 5.81 ± 0.41 (3, <0.0001 ****) |
|  | **d2 (N, p-value)** |
| **non-stress** | 12.08 ± 2.15 (7, <0.0001 ****) |
| **Tg stress** | 2.43 ± 0.16 (6, <0.0001 ****) |
| **fold-induction** | 0.98 ± 0.06 (4, 0.5065 ns) |

**Table S4.** Reference for Figure S3 and Table S3; relative ATF4-HA protein expression with wt set to 1 – individual values. For wt ATF4-HA tag fold-induction see Table S2.

|  | **d-all** | **St-st-only** | **uORF1-only** | **uORF2-only** |
| --- | --- | --- | --- | --- |
| **non-stress** | 20.2108<br>20.1493<br>24.9716<br>25.1497<br>22.9260 | 19.2065<br>24.4600<br>18.0205 | 11.7912<br>14.5744<br>10.2913 | 0.2182<br>0.1407<br>0.2058<br>0.1765 |
| **Tg stress** | 2.8184<br>3.3113<br>3.7961 | 2.5967<br>3.1636<br>3.3866 | 2.3353<br>2.8305<br>2.8535 | 0.2745<br>0.2188<br>0.1998 |
| **fold-induction** | 0.9906<br>1.0773<br>1.0164 | 0.8076<br>0.9861<br>0.9930 | 0.9920<br>0.9045<br>0.9633 | 2.8828<br>3.2869<br>3.5573 |
|  | **dSt-st** | **d1** | **d2** |  |
| **non-stress** | 0.9732<br>0.8915<br>0.8889<br>0.8518<br>1.0173<br>1.1380 | 0.2875<br>0.2954<br>0.2919 | 14.0385<br>15.4196<br>8.8531<br>11.3319<br>10.7310<br>12.0588<br>12.1492 |  |
| **Tg stress** | 0.9235<br>1.0011<br>0.9185 | 0.3767<br>0.3440<br>0.2564 | 2.5079<br>2.4676<br>2.5781<br>2.2401<br>2.5556<br>2.2038 |  |
| **fold-induction** | 6.1441<br>5.5343<br>5.6200 | 6.2766<br>5.6519<br>5.5149 | 0.9584<br>0.9709<br>1.0598<br>0.9330 |  |

**Table S5.** Reference for Figure 3B; relative ATF4-HA protein expression with wt (or parental d2) constructs set to 1. For wt ATF4-HA tag fold-induction see Table S1, for d2 fold-induction – Table S3.

|  | **wt-SL3<sup>Mut-1</sup> (N, p-value)** | **wt- SL3<sup>Mut-2</sup> (N, p-value)** |
| --- | --- | --- |
| **non-stress** | 1.31 ± 0.13 (5, 0.0007 ***) | 1.72 ± 0.16 (4, 0.0001 ***) |
| **Tg stress** | 1.53 ± 0.28 (6, 0.0022 **) | 1.84 ± 0.22 (3, 0.0027 **) |
| **fold-induction** | 5.69 ± 0.59 (6, < 0.0001 ****) | 5.87 ± 0.18 (3, < 0.0001 ****) |
|  | **d2-SL3<sup>Mut-1</sup> (N, p-value)** |  |
| **non-stress** | 1.69 ± 0.33 (6, 0.0022 **) |  |
| **Tg stress** | 1.40 ± 0.21 (5, 0.0027 **) |  |
| **fold-induction** | 0.99 ± 0.12 (7, 0.8315 ns) |  |

**Table S6.** Reference for Figure 3B and Table S5; relative ATF4-HA protein expression with wt (or parental d2) construct set to 1 – individual values. For wt ATF4-HA tag fold-induction see Table S2, for d2 fold-induction – Table S4.

|  | **wt-SL3<sup>Mut-1</sup>** | **d2-SL3<sup>Mut-1</sup>** | **wt-SL3<sup>Mut-2</sup>** |
| --- | --- | --- | --- |
| **non-stress** | 1.3549 | 1.3556 | 1.5097 |
|  | 1.3636 | 1.9470 | 1.6794 |
|  | 1.1961 | 1.3943 | 1.7911 |
|  | 1.1578 | 1.4266 | 1.8890 |
|  | 1.4705 | 2.0385 |  |
|  |  | 1.9983 |  |
| **Tg stress** | 1.4009 | 1.1336 | 1.9355 |
|  | 1.5382 | 1.5362 | 1.9997 |
|  | 1.2837 | 1.6662 | 1.5906 |
|  | 1.4387 | 1.3437 |  |
|  | 1.4448 | 1.3058 |  |
|  | 2.0767 |  |  |
| **fold-induction** | 4.7580 | 0.8509 | 5.6622 |
|  | 5.2583 | 0.8849 | 5.9618 |
|  | 6.2353 | 1.0014 | 5.9843 |
|  | 5.6282 | 0.9475 |  |
|  | 6.2162 | 0.9471 |  |
|  | 6.0593 | 1.1995 |  |
|  |  | 1.0983 |  |

**Table S7.** Reference for Figure 4B; relative ATF4-HA protein expression with wt set to 1. For wt ATF4-HA tag fold-induction see Table S1; for average values of wt-SL3^Mut-1^ construct see Table S5.

|  | **wt-CUG<sup>Mut-1</sup> (N, p-value)** | **wt-SL3<sup>Mut-1</sup>-CUG<sup>Mut</sup> (N, p-value)** |
| --- | --- | --- |
| **non-stress** | 0.78 ± 0.07 (3, 0.0072 **) | 1.04 ± 0.07 (3, 0.4482 ns) |
| **Tg stress** | 0.81 ± 0.06 (4, 0.0009 ***) | 1.13 ± 0.11 (3, 0.1065 ns) |
| **fold-induction** | 5.56 ± 0.59 (3, 0.0002 ***) | 5.40 ± 0.33 (3, <0.0001****) |

**Table S8.** Reference for Figure 4B and Table S7; relative ATF4-HA protein expression with wt set to 1 – individual values. For wt ATF4-HA tag fold-induction see Table S2; for individual values of wt-SL3^Mut-1^ construct see Table S6.

|  | **WT-CUG<sup>Mut</sup>**<br>**8h** | **WT-SL3<sup>Mut-1</sup>-**<br>**CUG<sup>Mut</sup>**<br>**8h** |
| --- | --- | --- |
| **non-stress** | 0.8477<br>0.8005<br>0.7022 | 0.9834<br>1.0037<br>1.1208 |
| **Tg stress** | 0.8775<br>0.7303<br>0.8225<br>0.8214 | 1.0397<br>1.0960<br>1.2442 |
| **fold-induction** | 6.1537<br>5.5542<br>4.9730 | 5.7668<br>5.2947<br>5.1425 |

**Table S9.** Reference for Figure 5; relative ATF4-HA protein expression with wt (or parental, i.e., wt-SL3^Mut^ or d2, or d2-SL3^Mut^) constructs set to 1. For wt ATF4-HA tag fold-induction see Table S1; for d2 – Table S3; for wt-SL3^Mut^ and d2-SL3^Mut^ constructs – Table S5.

|  | wt_ins (N, p-value) | d2_ins (N, p-value) |
| --- | --- | --- |
| **non-stress** | 1.39 ± 0.14 (5, 0.0002 ***) | 1.81 ± 0.28 (4, 0.0011 **) |
| **Tg stress** | 1.34 ± 0.22 (4, 0.0219 *) | 1.43 ± 0.31 (6, 0.0067 **) |

|  | wt_ins_SL3<sup>Mut-1</sup> (N, p-value) | d2_ins_SL3<sup>Mut-1</sup> (N, p-value) |
| --- | --- | --- |
| **non-stress** | 0.99 ± 0.10 (4, 0.7780 ns) | 1.59 ± 0.30 (6, 0.0007 ***) |
| **Tg stress** | 1.02 ± 0.11 (4, 0.7349 ns) | 1.26 ± 0.32 (8, 0.0002 ***) |

**Table S10.**
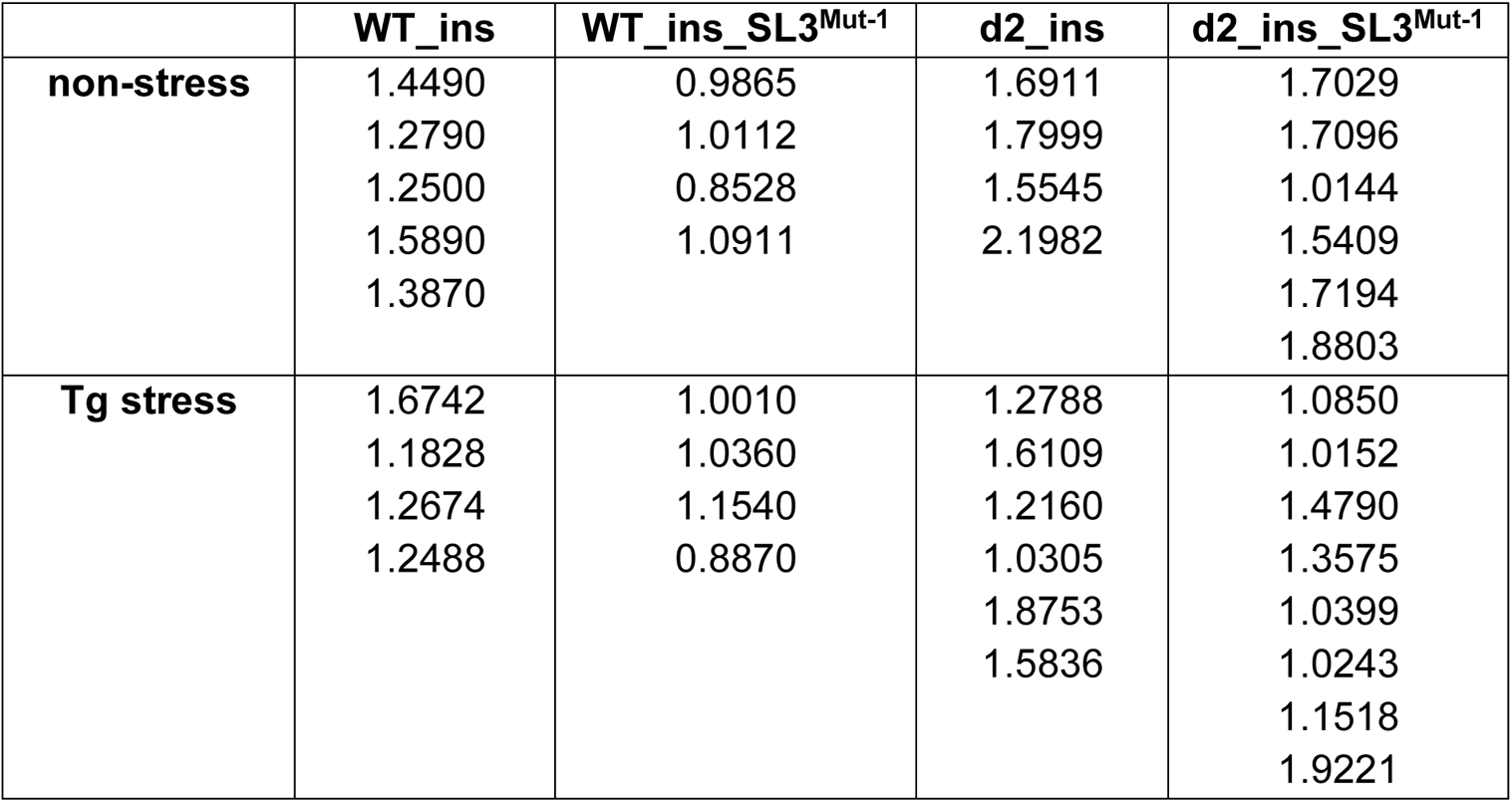
Reference for Figure 5 and Table S9; relative ATF4-HA protein expression with wt (or parental, i.e., wt-SL3^Mut^ or d2, or d2-SL3^Mut^) constructs set to 1 – individual values. For wt ATF4-HA tag fold-induction see Table S2; for d2 – Table S4; for wt-SL3^Mut^ or d2-SL3^Mut^ constructs see Table S6.

**Table S11.** Reference for Figure S7; relative 3AUG^Mut^ (or uORF2_ins) protein expression under 3 hours of thapsigargin (Tg) stress compared to 3AUG^Mut^ (or uORF2_ins, respectively) under non-stress (ns) set to 1. For wt ATF4-HA tag fold-induction see Table S1.

|  | 3AUG<sup>Mut</sup> (N, p-value) | uORF2_ins (N, p-value) |
| --- | --- | --- |
| **fold-induction** | 5.98 ± 0.31 (3, <0.0001 ****) | 0.62 ± 0.12 (4, 0.0006 ***) |

**Table S12.** Reference for Figure S7 and Table S11; relative 3AUG^Mut^ (or uORF2_ins) protein expression under 3 hours of thapsigargin (Tg) stress compared to 3AUG^Mut^ (or uORF2_ins, respectively) under non-stress (ns) set to 1 – individual values. For wt ATF4-HA tag fold-induction see Table S2.

|  | **3AUG<sup>Mut</sup>** | **uORF2_ins** |
| --- | --- | --- |
| **fold-induction** | 5.7269 | 0.7550 |
|  | 6.3253 | 0.6555 |
|  | 5.8946 | 0.4766 |
|  |  | 0.6090 |

**Table S13.**
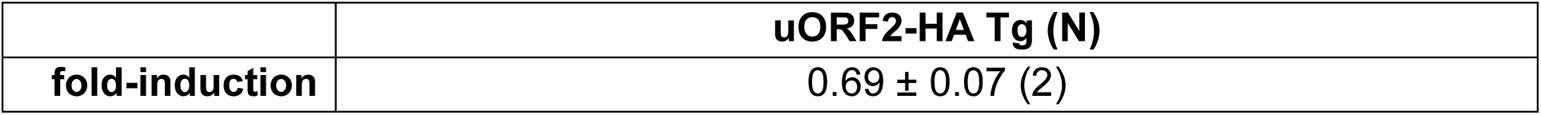
Reference for Figure S9B; relative uORF2-HA fusion protein expression under 3 hours of thapsigargin (Tg) stress compared to uORF2-HA under non-stress (ns) set to 1.

|  | **uORF2-HA Tg (N)** |
| --- | --- |
| **fold-induction** | 0.69 ± 0.07 (2) |

**Table S14.**
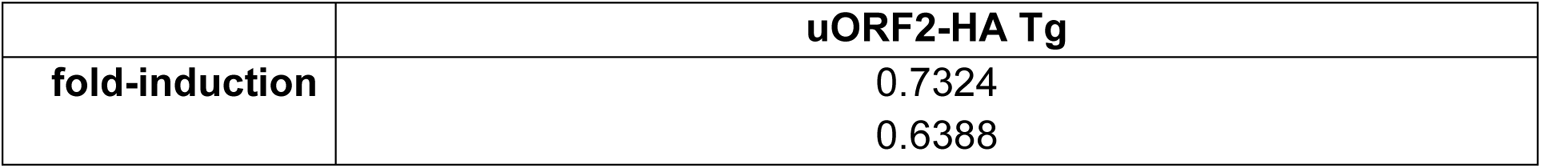
Reference for Figure S9B and Table S13; relative uORF2-HA fusion protein expression under 3 hours of thapsigargin (Tg) stress compared to uORF2-HA under non-stress (ns) set to 1 – individual values.

**Table S15.**
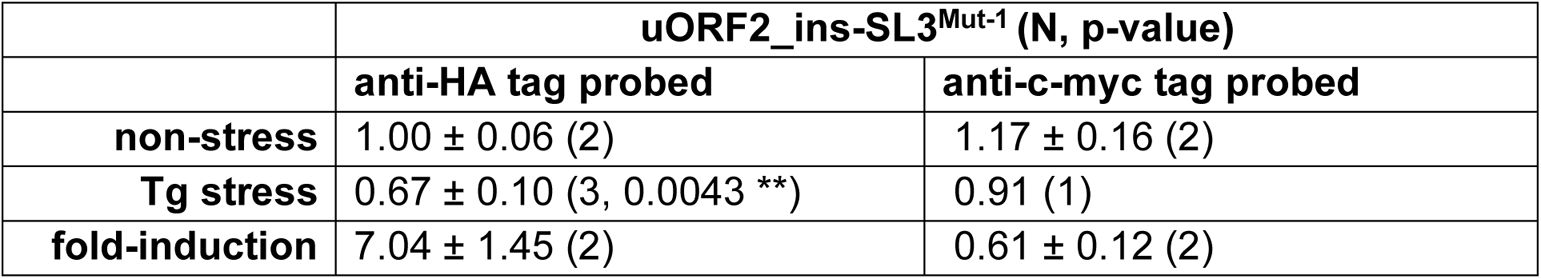
Reference for Figure S10A; relative ATF4-HA protein expression with uORF2_ins set to 1. For uORF2_ins fold-induction see Table S11.

|  | **uORF2_ins-SL3<sup>Mut-1</sup> (N, p-value)** |  |
| --- | --- | --- |
|  | **anti-HA tag probed** | **anti-c-myc tag probed** |
| **non-stress** | 1.00 ± 0.06 (2) | 1.17 ± 0.16 (2) |
| **Tg stress** | 0.67 ± 0.10 (3, 0.0043 **) | 0.91 (1) |
| **fold-induction** | 7.04 ± 1.45 (2) | 0.61 ± 0.12 (2) |

**Table S16.** Reference for Figure S10A and Table S15; relative ATF4-HA protein expression with uORF2_ins set to 1 – individual values. For uORF2_ins fold-induction see Table S11.

|  | **uORF2_ins-SL3<sup>Mut-1</sup>** |  |
| --- | --- | --- |
|  | **anti-HA tag probed** | **anti-c-myc tag probed** |
| **non-stress** | 1.0382 | 1.0564 |
|  | 0.9510 | 1.2776 |
| **Tg stress** | 0.5718 | 0.9147 |
|  | 0.6703 |  |
|  | 0.7681 |  |
| **fold-induction** | 6.0108 | 0.6978 |
|  | 8.0632 | 0.5295 |

**Table S17.**
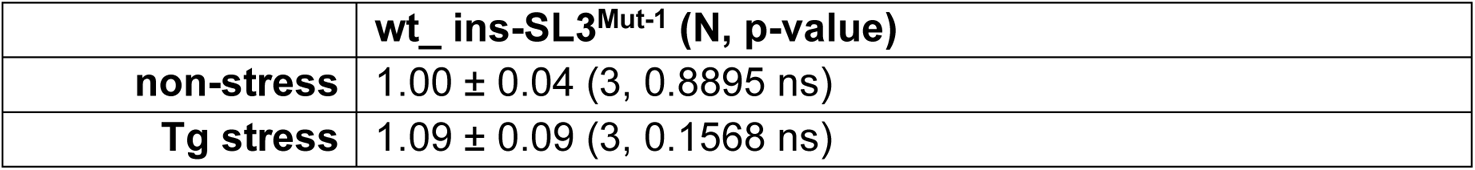
Reference for Figure S10B; relative ATF4-HA protein expression with wt_ins set to 1.

**Table S18.** Reference for Figure S10B and Table S17; relative ATF4-HA protein expression with wt_ins set to 1 – individual values.

|  | **wt_ins-SL3<sup>Mut-1</sup>** |
| --- | --- |
| **non-stress** | 0.9781 |
|  | 0.9699 |
|  | 1.0419 |
| **Tg stress** | 0.9963 |
|  | 1.1847 |
|  | 1.1039 |

**Table S19.** Reference for Figure 7B (left and middle panels); relative ATF4-HA protein expression with wt set to 1. For wt ATF4-HA tag fold-induction see Table S1.

|  | **wt-A\*235G (N, p-value)** | **wt-A\*326G (N, p-value)** |
| --- | --- | --- |
| **non-stress** | 1.34 ± 0.06 (4, < 0.0001 ****) | 0.78 ± 0.09 (4, 0.0032 **) |
| **Tg stress** | 1.10 ± 0.15 (3, 0.2960 ns) | 0.83 ± 0.16 (4, 0.0681 ns) |
| **fold-induction** | 6.22 ± 0.57 (4, < 0.0001 ****) | 4.83 ± 0.55 (4, < 0.0001 ****) |

**Table S20.**
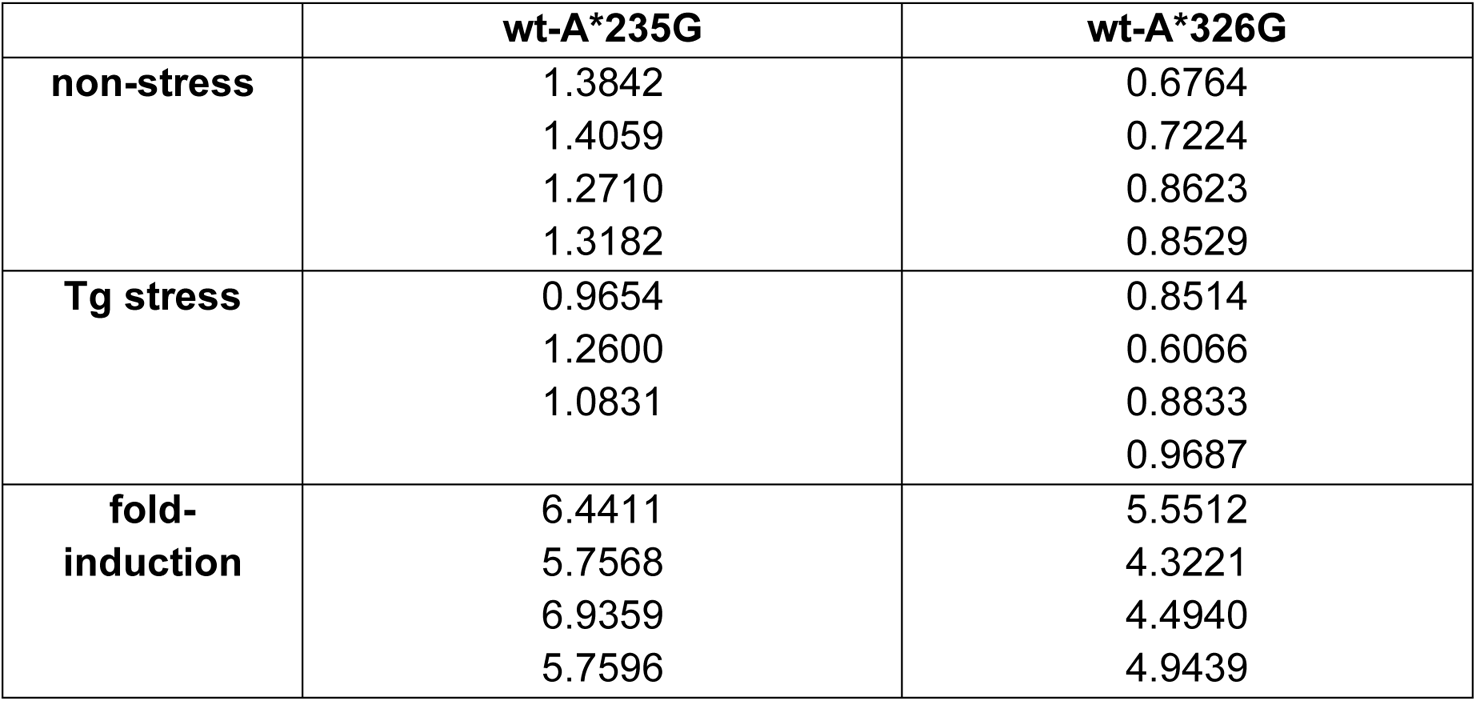
Reference for Figure 7B (left and middle panels) and Table S17; relative ATF4-HA protein expression with wt set to 1 – individual values. For wt ATF4-HA tag fold-induction see Table S2.

**Table S21.** Reference for Figure 7C; relative levels of amplification efficiencies of *ATF4* transcript regions containing either A_235_ or A_326_ in mock treated *ATF4* mRNA with Tg treated set to 1.

| **HEK293T** | **A<sub>235</sub> (N, p-value)** | **A<sub>326</sub> (N, p-value)** |
| --- | --- | --- |
| **5 min** | 0.5025 ± 0.0990 (4, 0.0032 **) | 0.8725 ± 0.0192 (4, 0.0014 **) |
| **15 min** | 0.5275 ± 0.0629 (4, 0.001 ***) | 0.92 ± 0.0604 (4, 0.1056 ns) |
| **60 min** | 0.95 ± 0.2152 (4, 0.7144 ns) | 0.8475 ± 0.0675 (4, > 0.0298 *) |
| **HeLa** | **A<sub>235</sub> (N, p-value)** | **A<sub>326</sub> (N, p-value)** |
| **5 min** | 0.4325 ± 0.1229 (4, 0.0041 **) | 0.89 ± 0.2410 (4, 0.4870 ns) |
| **15 min** | 0.3013 ± 0.1298 (4, 0.0026 **) | 0.85 ± 0.2178 (4, 0.3187 ns) |
| **60 min** | 0.5025 ± 0.3338 (4, > 0.0817 ns) | 0.995 ± 0.1265 (4, > 0.9498 ns) |

**Table S22.** Reference for Figure 7B (right panel); relative ATF4-HA protein expression with wt set to 1.

|  | **wt-SL3<sup>Mut-1</sup>-A\*326G (N, p-value)** |
| --- | --- |
| **non-stress** | 0.86 ± 0.04 (3, 0.0044 **) |
| **Tg stress** | 1.59 ± 0.16 (3, 0.0032 **) |
| **fold-induction** | 6.48 ± 0.34 (3, < 0.0001 ****) |

**Table S23.** Reference for Figure 7B (right panel) and Table S2; relative ATF4-HA protein expression with wt set to 1 – individual values.

|  | **wt-SL3<sup>Mut-1</sup>-A\*326G** |
| --- | --- |
| **non-stress** | 0.8557<br>0.8275<br>0.9078 |
| **Tg stress** | 1.7344<br>1.4163<br>1.6309 |
| **fold-induction** | 6.8769<br>6.2719<br>6.3050 |

**Table S24.** Plasmids used in this study.

| **Plasmid** | **Description** | **Source of reference** |
| --- | --- | --- |
| *pCMV-EGFP-N2* | high copy Clontech empty vector *EGFP-N2* with enhanced green fluorescent protein gene | (Roithova et al., 2020) |
| wt *ATF4*-HA Tag | high copy h*ATF4*-wt-HA-Tag plasmid containing 5'UTR of *ATF4* and 3'UTR of *ATF4* in high copy *pCMV-EGFP-N2* vector | This study |
| dSt-st | high copy h*ATF4*-dSt-st in wt *ATF4*-HA Tag plasmid containing point mutation in St-st AUG/AGG | This study |
| d1 | high copy h*ATF4*-d1 in wt *ATF4*-HA Tag plasmid containing point mutation in uORF1 AUG/AGG | This study |
| d2 | high copy h*ATF4*-d2 in wt *ATF4*-HA Tag plasmid containing point mutation in uORF2 AUG/AGG | This study |
| d-all | high copy h*ATF4*-d-all-HA-Tag plasmid containing 5'UTR of *ATF4* and 3'UTR of *ATF4*, carrying point mutation in St-st, uORF1 and uORF2 – AUG/AGG, in high copy *pCMV-EGFP-N2* vector | This study |
| St-st-only | high copy h*ATF4*-St-st-only in wt *ATF4*-HA Tag plasmid containing point mutation in uORF1 and uORF2 – AUG/AGG | This study |
| uORF1-only | high copy h*ATF4*-uORF1-only in wt *ATF4*-HA Tag plasmid containing point mutation in St-st and uORF2 – AUG/AGG | This study |
| uORF2-only | high copy h*ATF4*-uORF2-only in wt *ATF4*-HA Tag plasmid containing point mutation in St-st and uORF1 – AUG/AGG | This study |
| dSt-st-NC1 | high copy h*ATF4*-dSt-st-NC1 in dSt-st plasmid containing substitution of CUG near-cognate +7 to +9 downstream of St-st by CUA | This study |
| St-st-only-NC1 | high copy h*ATF4*-St-st-only-NC1 in St-st-only plasmid containing substitution of CUG near-cognate +7 to +9 downstream of St-st by CUA | This study |
| St-st-only-NC2/3 | high copy h*ATF4*-St-st-only-NC2/3 in St-st-only plasmid containing substitution of CUG near-cognate +7 to +9 downstream of | This study |
|  | St-st by CUA, and GUGUG +13 to +17 downstream of St-st – by GUACG |  |
| dSt-st-SL1 | high copy h*ATF4*-dSt-st-SL1 in dSt-st plasmid containing point mutations C/A, C/A, -3 and -7 upstream of St-st, respectively, plus point mutation in St-st AUG/AGG | This study |
| St-st-only-SL2-1 | high copy h*ATF4*-St-st-only-SL2-1 in St-st-only plasmid containing substitution of GGG -15 to -13 upstream of uORF1 by TTG | This study |
| St-st-only-SL2-2 | high copy h*ATF4*-St-st-only-SL2-2 in St-st-only plasmid containing substitution of CGC -23 to -21 upstream of uORF1 by TGC, and GGG -15 to -13 upstream of uORF1 by GTG | This study |
| dSt-st-SL2-1 | high copy h*ATF4*-dSt-st-SL2-1 in dSt-st plasmid containing substitution of GGG -15 to -13 upstream of uORF1 by TTG | This study |
| dSt-st-SL2-2 | high copy h*ATF4*-dSt-st-SL2-2 in dSt-st plasmid containing substitution of CGC -23 to -21 upstream of uORF1 by TGC, and GGG -15 to -13 upstream of uORF1 by GTG | This study |
| d-all-SL2-2 | high copy h*ATF4*-d-all-SL2-2 in d-all plasmid containing substitution of CGC -23 to -21 upstream of uORF1 by TGC, and GGG -15 to -13 upstream of uORF1 by GTG | This study |
| wt-A235G | high copy h*ATF4*-wt-A235G in wt *ATF4*-HA Tag plasmid containing point mutation +47 downstream of AUG of uORF2 – A/G | This study |
| wt-SL3<sup>Mut-1</sup> | high copy h*ATF4*-wt-SL3<sup>Mut-1</sup> in wt *ATF4*-HA Tag plasmid containing substitution of the sequence CCCCC +50 to +54 downstream of first AUG of ATF4 by CAAAA | This study |
| wt-SL3<sup>Mut-2</sup> | high copy h*ATF4*-wt-SL3<sup>Mut-2</sup> in wt *ATF4*-HA Tag plasmid containing substitution of the sequence GGG +36 to +38 downstream of first AUG of ATF4 by AUC | This study |
| dSt-st-SL3<sup>Mut-1</sup> | high copy hATF4-dSt-st-SL3<sup>Mut-1</sup> in dSt-st plasmid containing substitution of the sequence CCCCC +50 to +54 downstream of first AUG of ATF4 by CAAAA | This study |
| d-all-SL3<sup>Mut-1</sup> | high copy hATF4-d-all-SL3<sup>Mut-1</sup> in d-all plasmid containing substitution of the sequence CCCCC +50 to +54 downstream of first AUG of ATF4 by CAAAA | This study |
| uORF2-only-SL3<sup>Mut-1</sup> | high copy hATF4-uORF2-only-SL3<sup>Mut-1</sup> in uORF2-only plasmid containing substitution of the sequence CCCCC +50 to +54 downstream of first AUG of ATF4 by CAAAA | This study |
| wt-AUG3<sup>Mut</sup> (3rd ATF4 AUG) | high copy hATF4-wt-AUG3<sup>Mut</sup> in wt ATF4-HA Tag plasmid containing point mutation of 3rd AUG of ATF4, located +46 to +48 downstream of 1st AUG – AUG/AGG | This study |
| dSt-st-AUG3<sup>Mut</sup> (3rd ATF4 AUG) | high copy hATF4-dSt-st-AUG3<sup>Mut</sup> in dSt-st plasmid containing point mutation of 3rd AUG of ATF4, located +46 to +48 downstream of 1st AUG – AUG/AGG | This study |
| wt-UUG<sup>Mut</sup>-AUG3 | high copy hATF4-wt-UUG<sup>Mut</sup>-AUG3 in wt ATF4-HA Tag plasmid containing substitution in amino acid preceding 3rd AUG of ATF4 – UUG by GCA | This study |
| wt-SL3<sup>Mut-1</sup>-AUG3<sup>Mut</sup> | High copy hATF4-wt-SL3<sup>Mut-1</sup>-AUG3<sup>Mut</sup> in wt ATF4-HA Tag plasmid containing mutations of SL3<sup>Mut</sup> and AUG3<sup>Mut</sup> (3rd ATF4 AUG) | This study |
| wt-A326G | high copy hATF4-wt-A326G in wt ATF4-HA Tag plasmid containing substitution in potential methylation motif in SL3: A -5 upstream of third AUG of ATF4 by G | This study |
| wt-SL3<sup>Mut-1</sup>-A326G | high copy hATF4-wt-SL3<sup>Mut-1</sup>-A326G in wt ATF4-HA Tag plasmid containing substitution in potential methylation motif in SL3: A -5 upstream of third AUG of ATF4 by G as well as substitution of the sequence CCCCC +50 to +54 downstream of first AUG of ATF4 by CAAAA | This study |
| wt-3AUG<sup>Mut</sup> | high copy hATF4-wt-3AUG<sup>Mut</sup> in wt ATF4-HA Tag plasmid containing point mutation in the first AUG of ATF4, in the second AUG | This study |
|  | of ATF4 (located +7 to +9 downstream of the first), and in the third AUG of ATF4 (located in SL3, +46 to +48 downstream of the first AUG) – AUG/AGG |  |
| d2-3AUG<sup>Mut</sup> | high copy hATF4-d2-3AUG<sup>Mut</sup> in wt ATF4-HA Tag plasmid containing point mutation uORF2 AUG/AGG, then in the first AUG of ATF4, in the second AUG of ATF4 (located +7 to +9 downstream of the first), and in the third AUG of ATF4 (located in SL3, +46 to +48 downstream of the first AUG) – AUG/AGG | This study |
| 3AUG<sup>Mut</sup>-stop | high copy hATF4-3AUG<sup>Mut</sup>-stop in wt ATF4-HA Tag plasmid containing insertion of two Stop codons UAAUAA in ATF4 sequence after uORF2 Stop codon +109 to +114 downstream of first AUG of ATF4 | This study |
| 3AUG<sup>Mut</sup>- short | high copy hATF4-3AUG<sup>Mut</sup>-short in wt ATF4-HA Tag plasmid with the shortened ATF4: the sequence +712 to +1020 downstream of first AUG of ATF4 was removed | This study |
| 3AUG<sup>Mut</sup>- short-stop | high copy hATF4-3AUG<sup>Mut</sup>-short in 3AUG<sup>Mut</sup>-stop plasmid – 3AUG<sup>Mut</sup>-stop and 3AUG<sup>Mut</sup>- short mutations were combined | This study |
| uORF2_ins | high copy hATF4-uORF2_ins in wt ATF4-HA Tag plasmid with the extension of uORF2 sequence by a stretch of 10 copies of c-Myc Tag, inserted +22 downstream of AUG of uORF2, in uORF2 frame | This study |
| d2_ins (uORF2_ins => AUG to AGG) | high copy hATF4-d2_ins in wt ATF4-HA Tag plasmid containing point mutation in uORF2 AUG/AGG, then the extension of uORF2 sequence by a stretch of 10 copies of c-Myc Tag, inserted +22 downstream of AUG of uORF2, in uORF2 frame | This study |
| uORF2_ins_3AUG<sup>Mut</sup> | high copy hATF4-uORF2_ins_3AUG<sup>Mut</sup> in 3AUG<sup>Mut</sup> plasmid – 3AUG<sup>Mut</sup> and uORF2_ins mutations were combined | This study |
| wt-CUG<sup>Mut</sup> | high copy hATF4-wt-CUG<sup>Mut</sup> in wt ATF4-HA Tag plasmid containing substitution of CUG near-cognate +16 to +18 downstream of the first AUG of ATF4 by CUA | This study |
| Wt-SL3<sup>Mut-1</sup>-CUG<sup>Mut</sup> | high copy h*ATF4*-wt-SL3<sup>Mut-1</sup>-CUG<sup>Mut</sup> in wt *ATF4*-HA Tag plasmid – CUG<sup>Mut</sup> and SL3<sup>Mut-1</sup> mutations were combined | This study |
| wt_ins | high copy h*ATF4*-wt_ins in wt *ATF4*-HA Tag plasmid containing the extension of *ATF4* sequence by a stretch of 6 copies of c-Myc Tag, inserted +1 downstream of the first AUG of *ATF4* | This study |
| wt_ins-SL3<sup>Mut-1</sup> | high copy h*ATF4*-wt_ins-SL3<sup>Mut-1</sup> in wt *ATF4*-HA Tag plasmid – wt_ins and SL3<sup>Mut-1</sup> mutations were combined | This study |
| d2_ins | high copy h*ATF4*-d2_ins in wt_ins plasmid – combination of d2 and wt_ins mutations | This study |
| d2_ins-SL3<sup>Mut-1</sup> | high copy h*ATF4*-d2_ins-SL3<sup>Mut-1</sup> in wt_ins-SL3<sup>Mut-1</sup> plasmid – combination of d2 and wt_ins-SL3<sup>Mut-1</sup> mutations | This study |
| uORF2_ins-SL3<sup>Mut-1</sup> | high copy h*ATF4*-uORF2_ins-SL3<sup>Mut-1</sup> in wt_ins-SL3<sup>Mut-1</sup> plasmid – combination of uORF2_ins and wt-SL3<sup>Mut-1</sup> mutations | This study |
| uORF2-*ATF4*-HA fusion | high copy h*ATF4*-uORF2-*ATF4*-HA fusion in wt *ATF4*-HA Tag plasmid containing mutation in uORF2 Stop codon UGA to GA | This study |

**Table S25.**
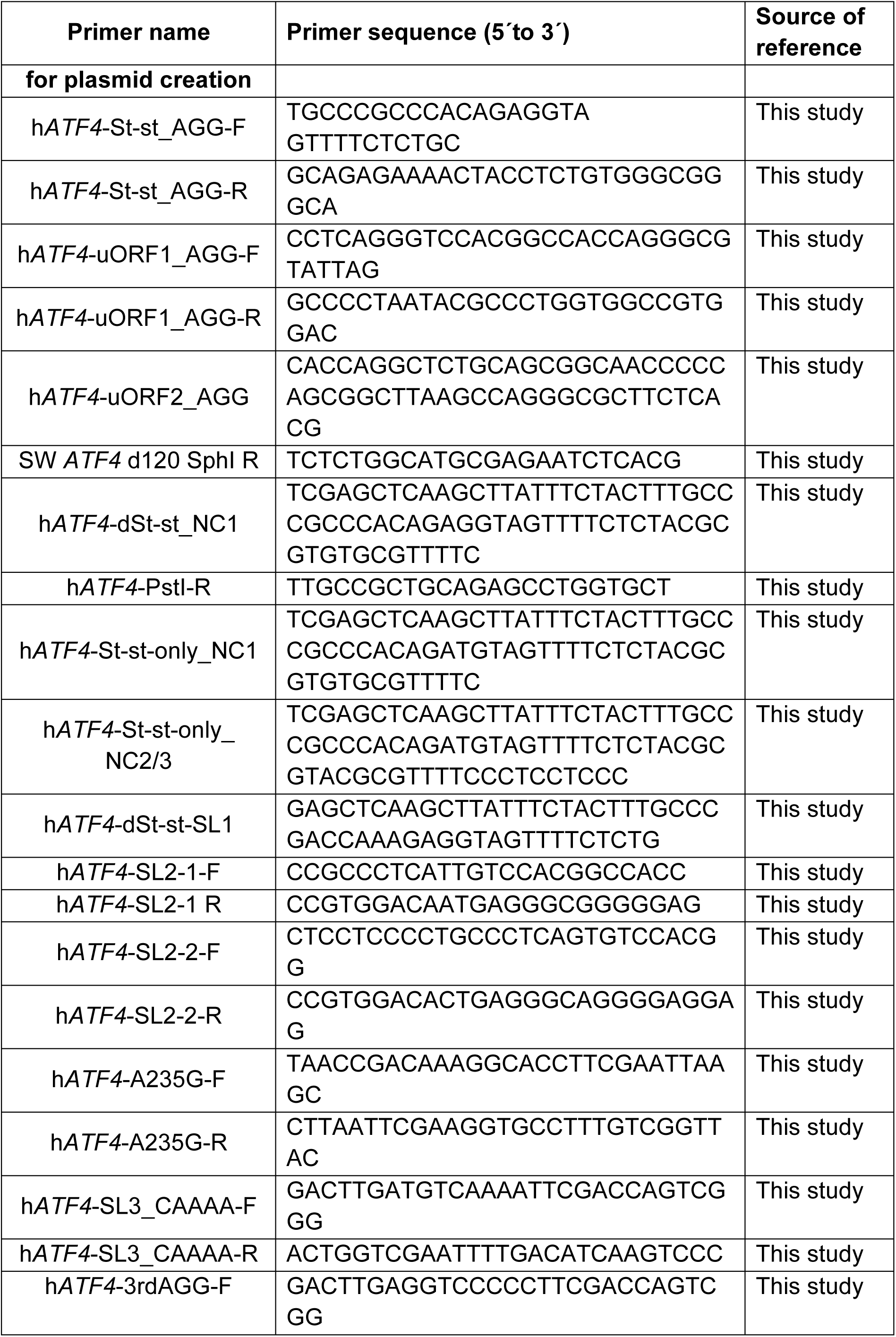
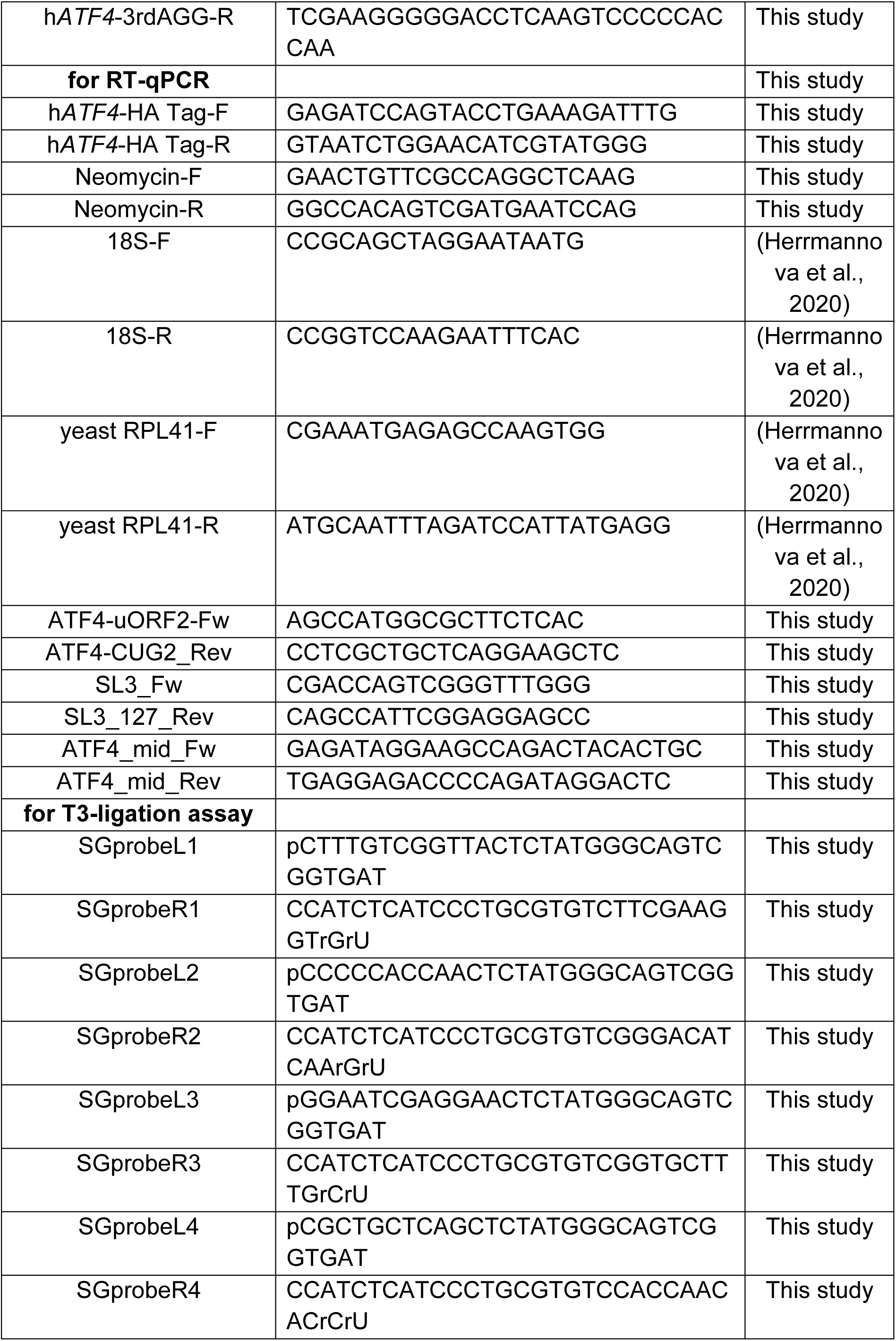
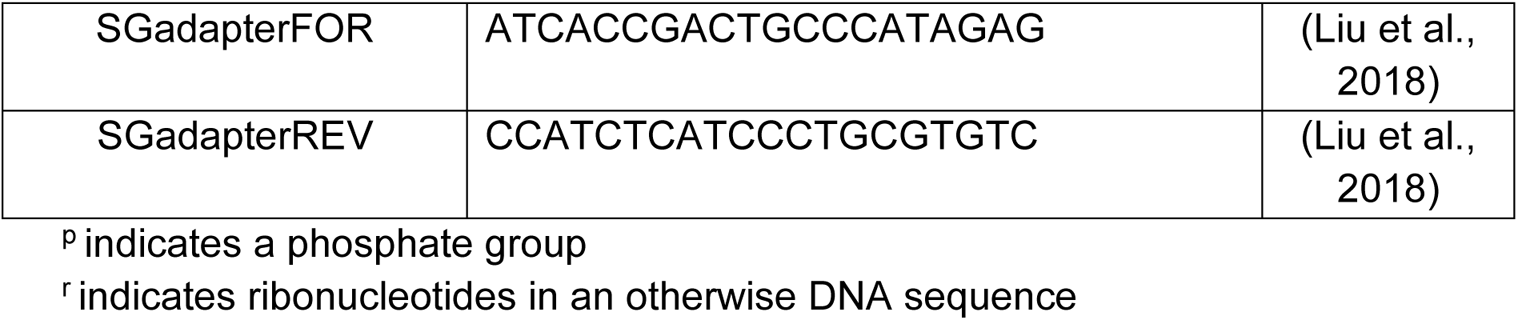
Primers (Eurofins Genomics) used in this study.

**Table S26.**
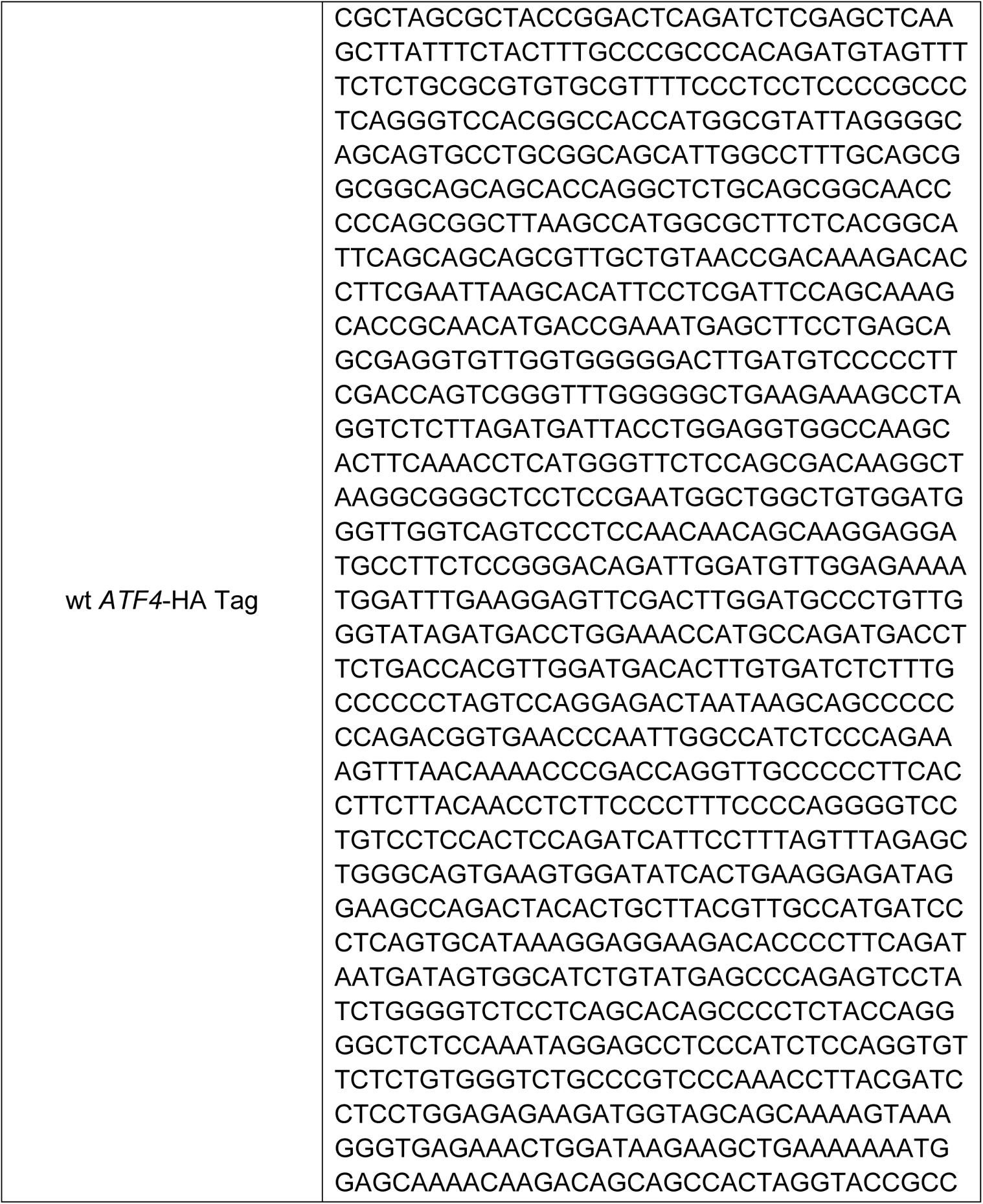
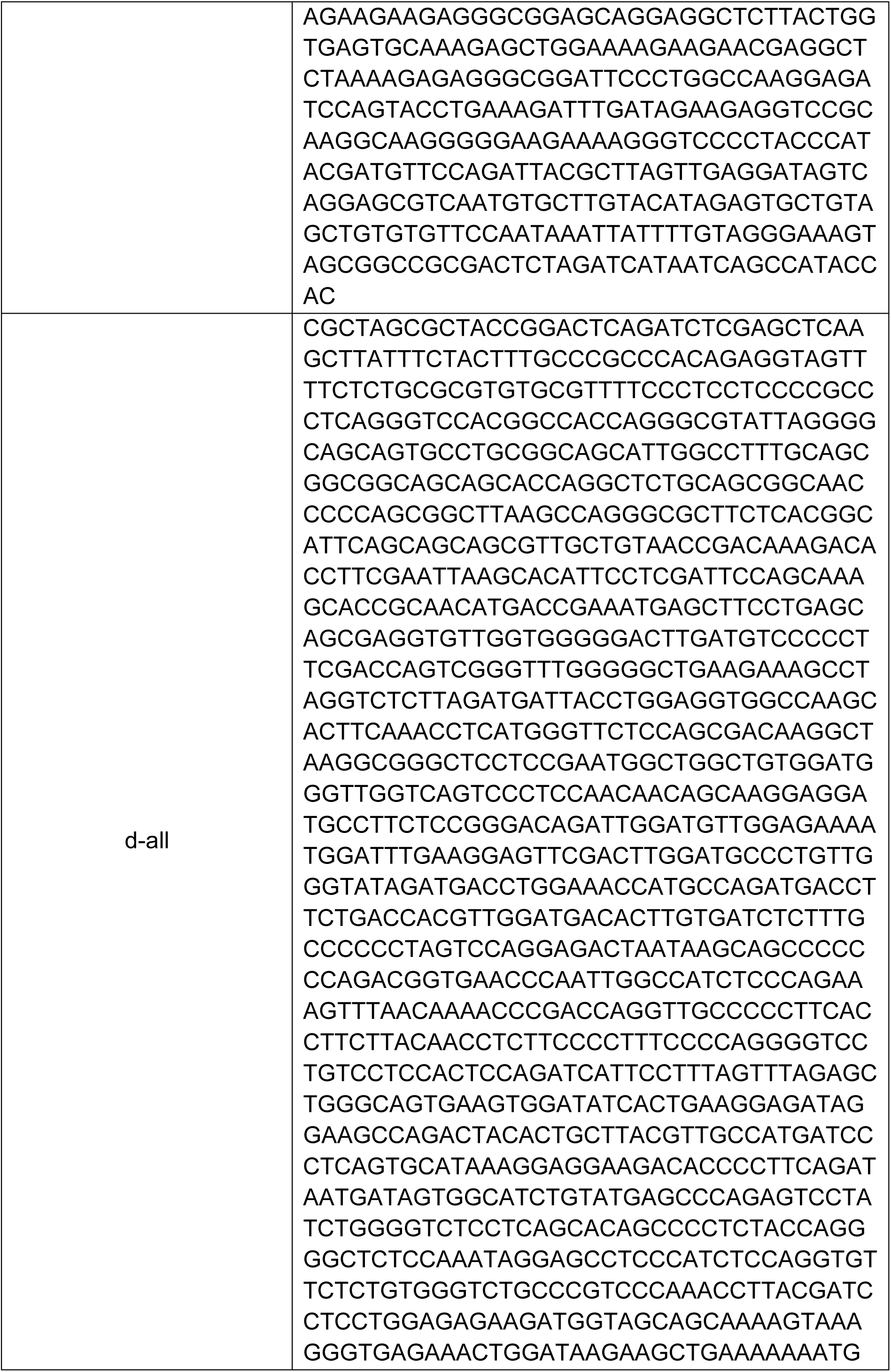
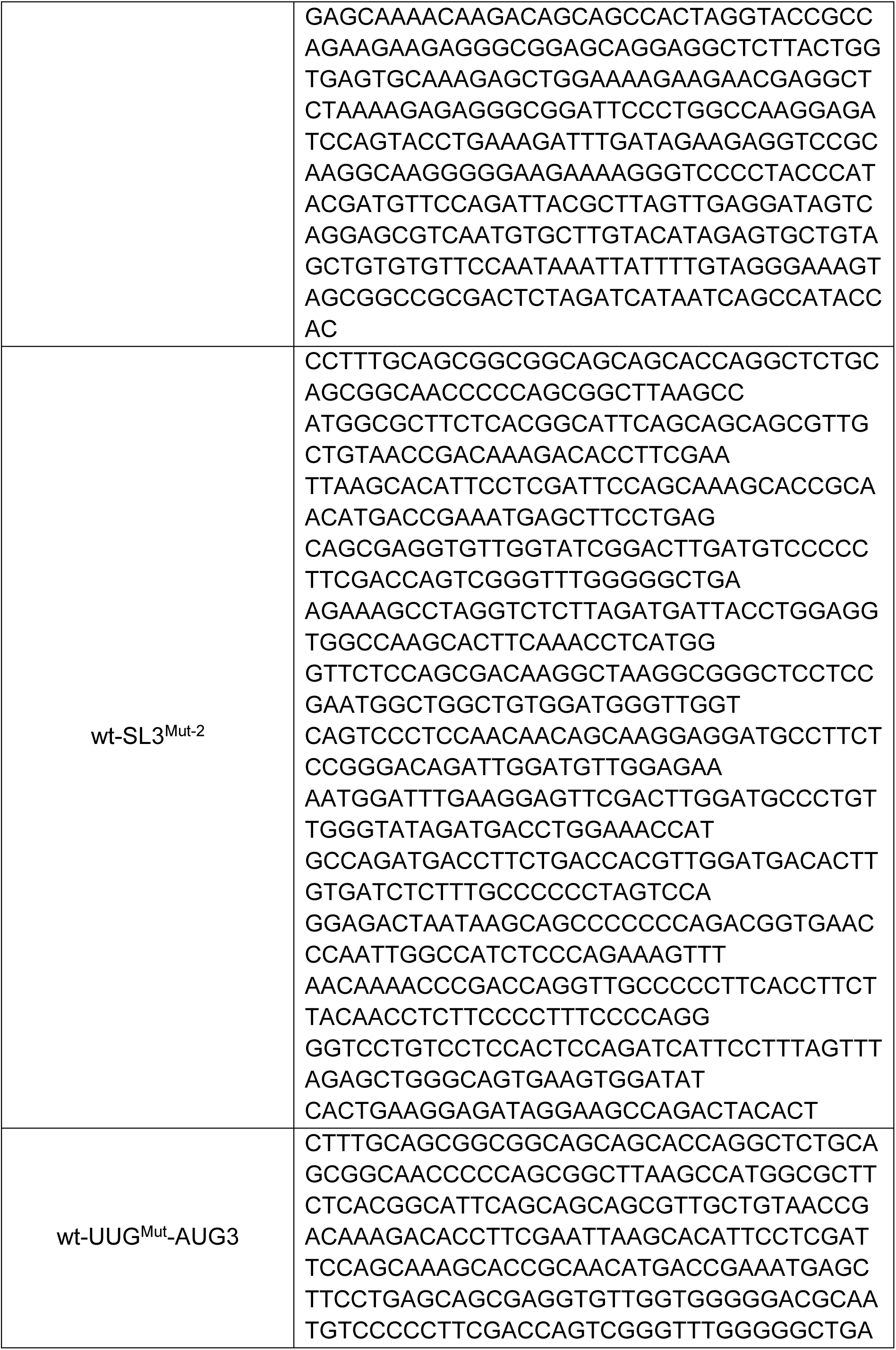
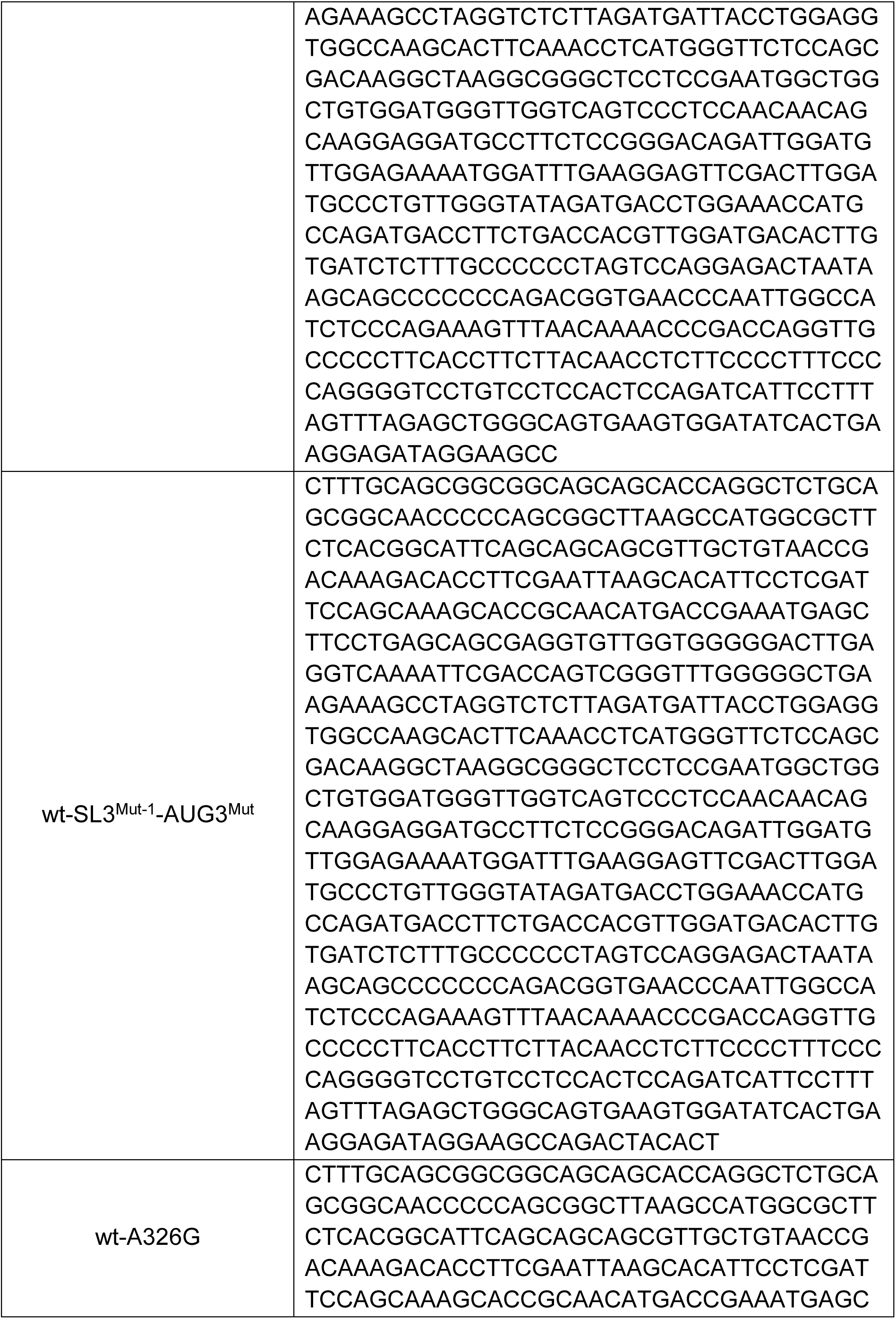
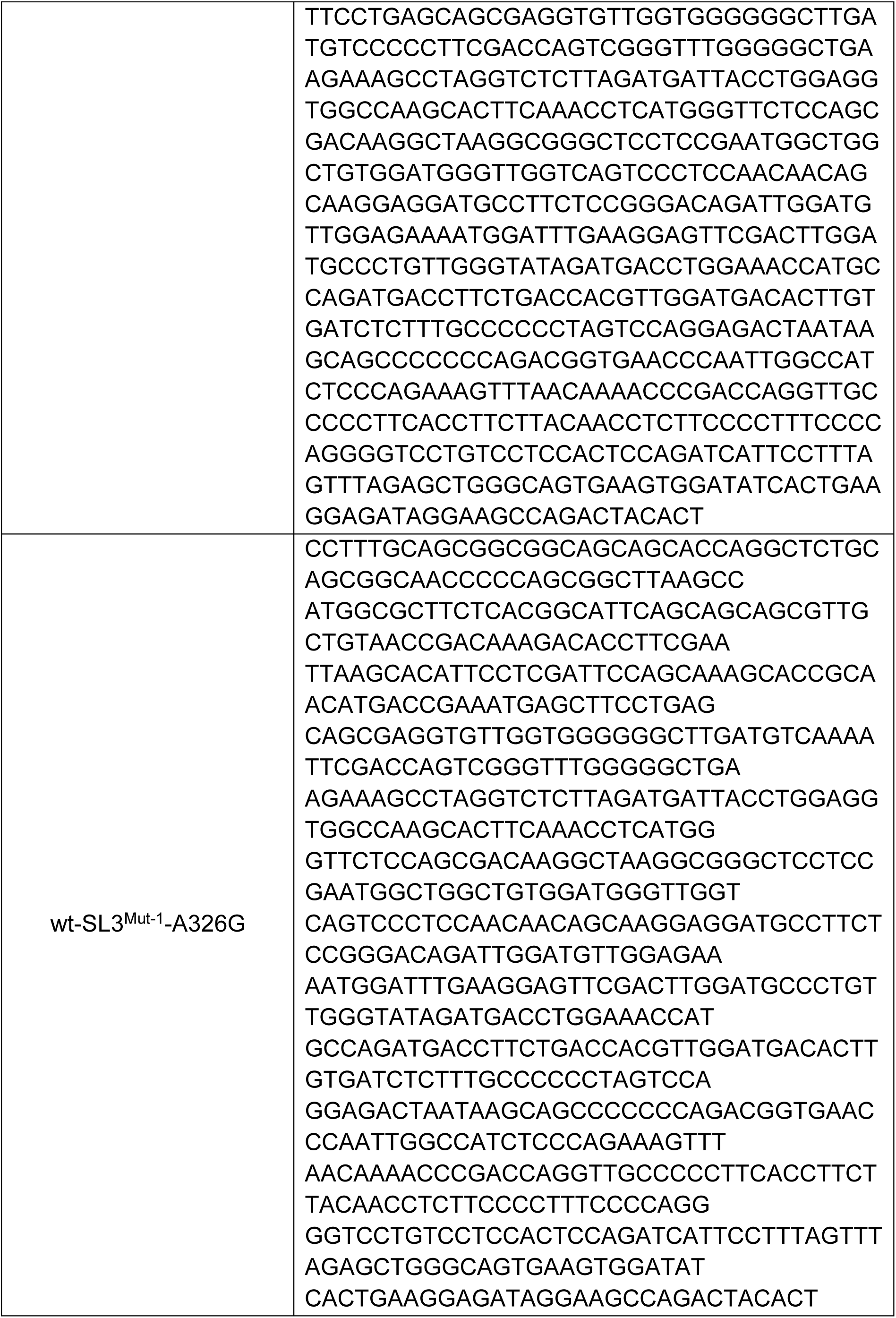
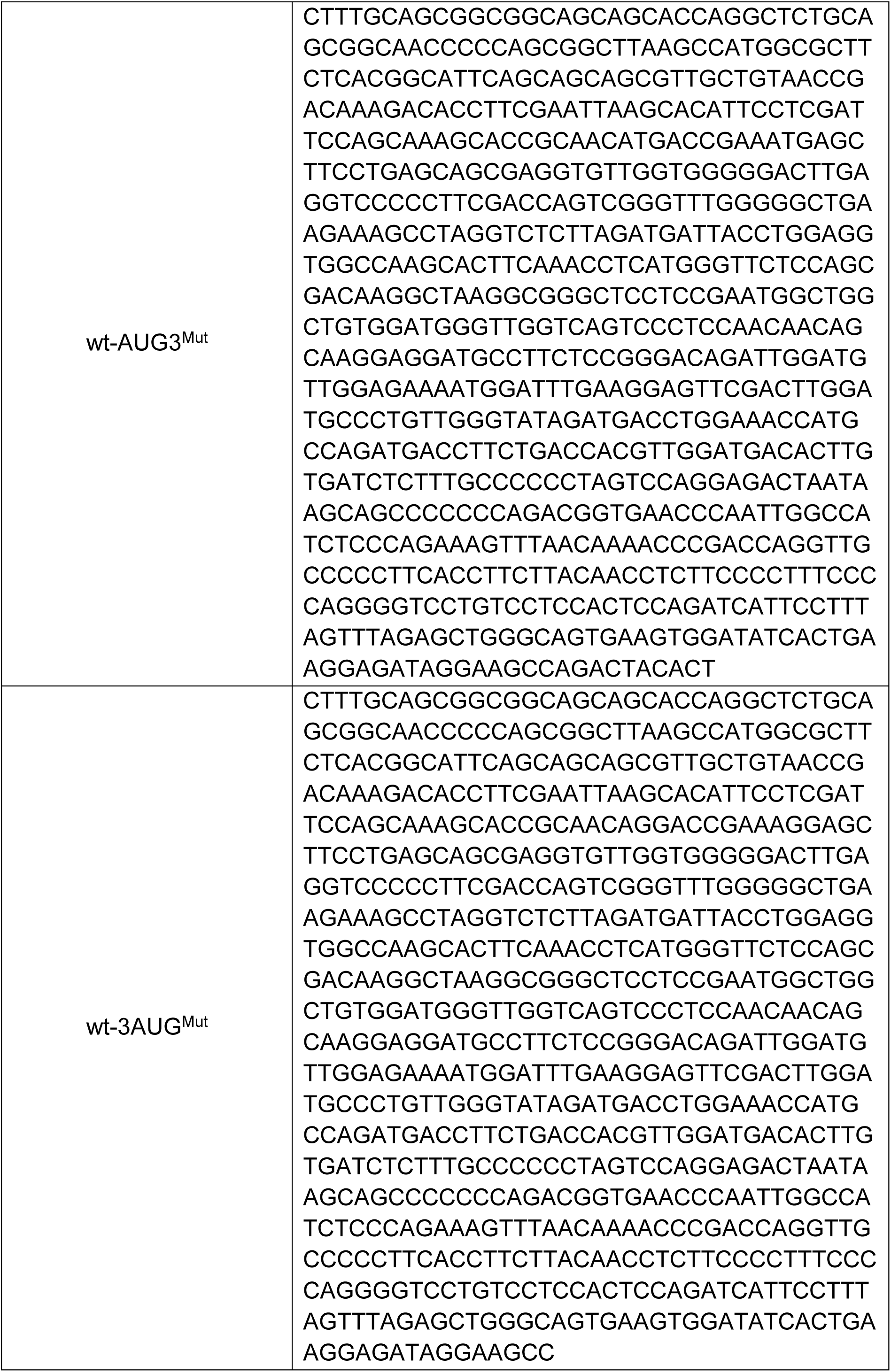
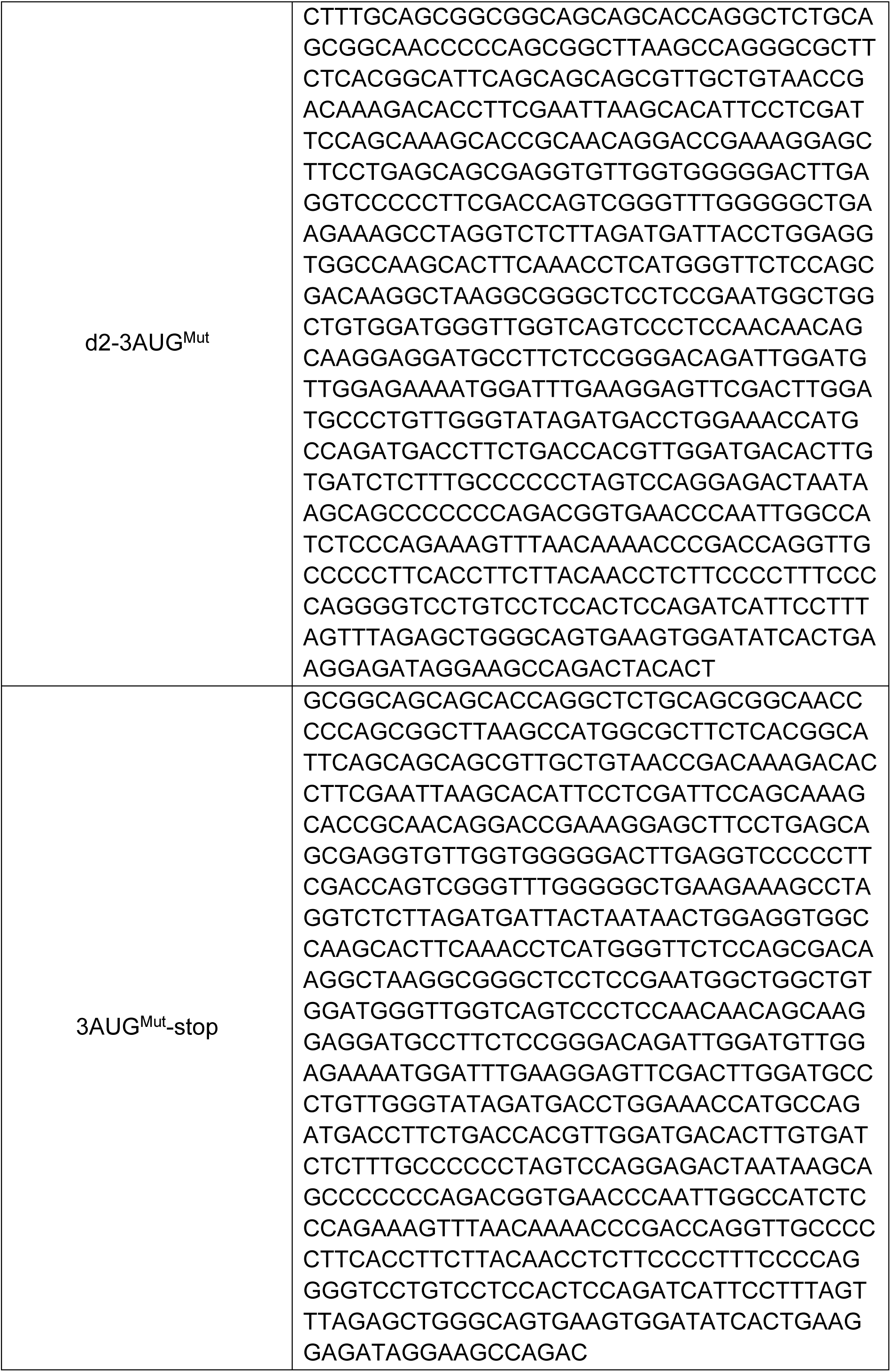
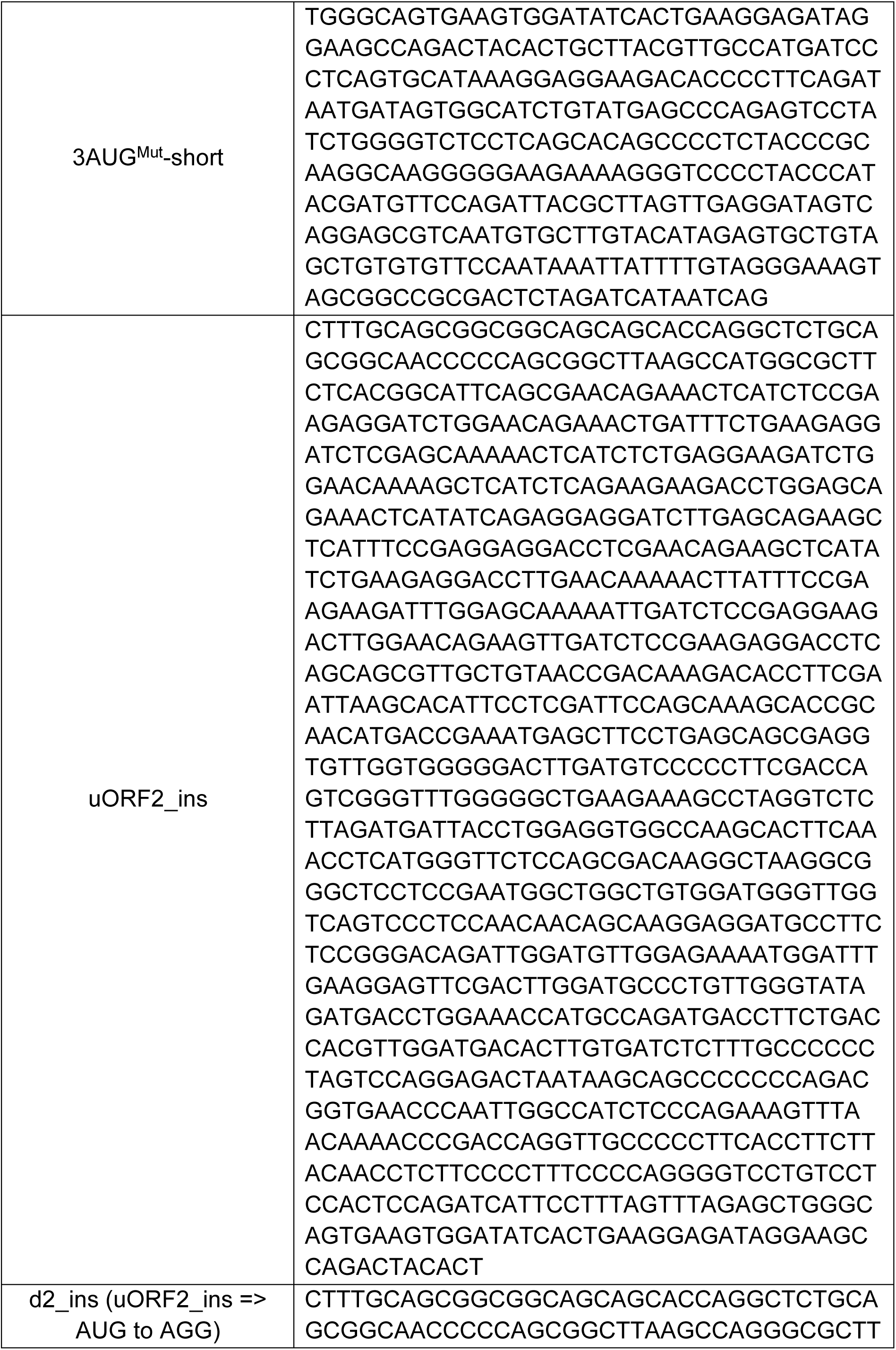
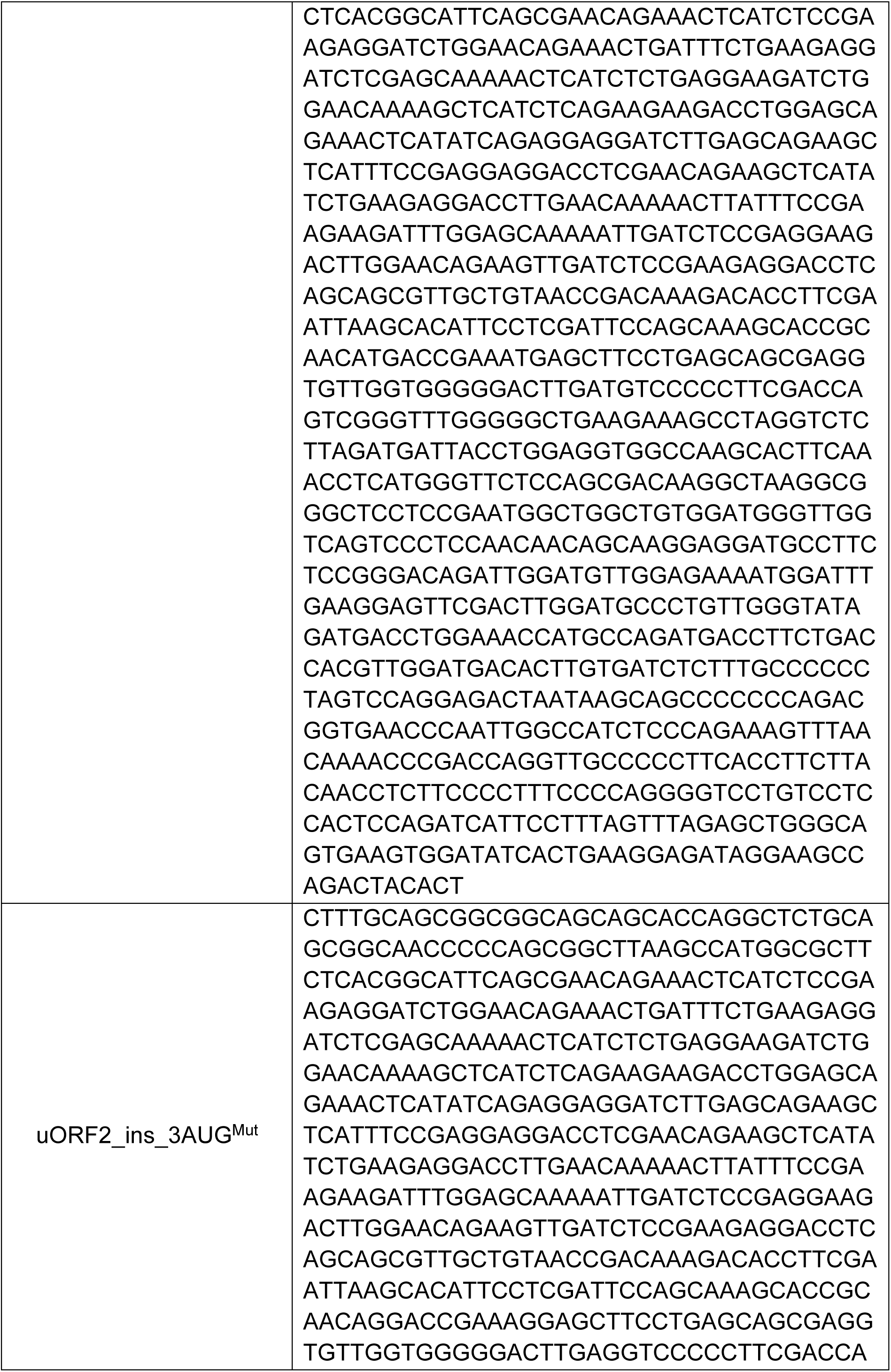
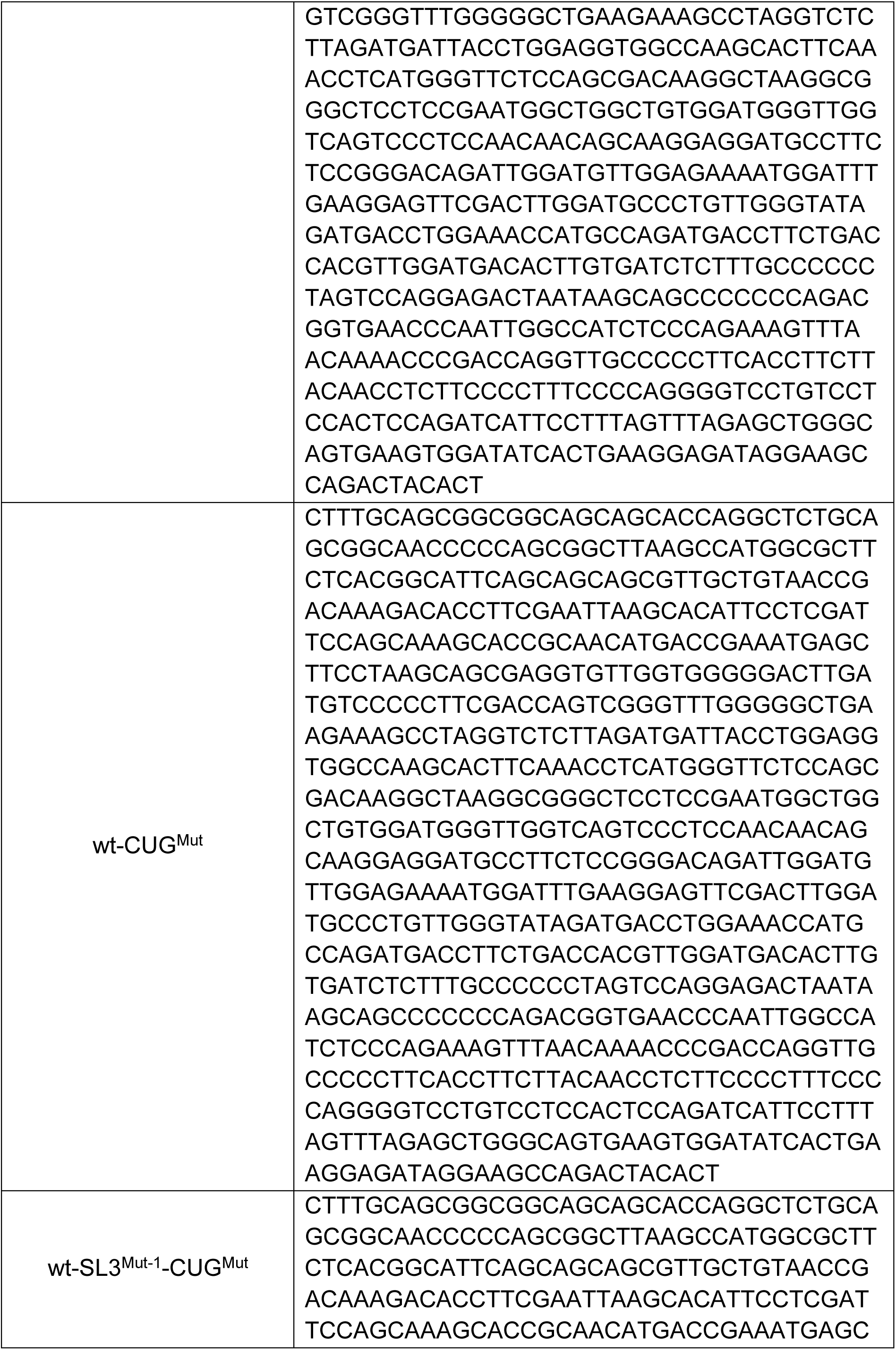
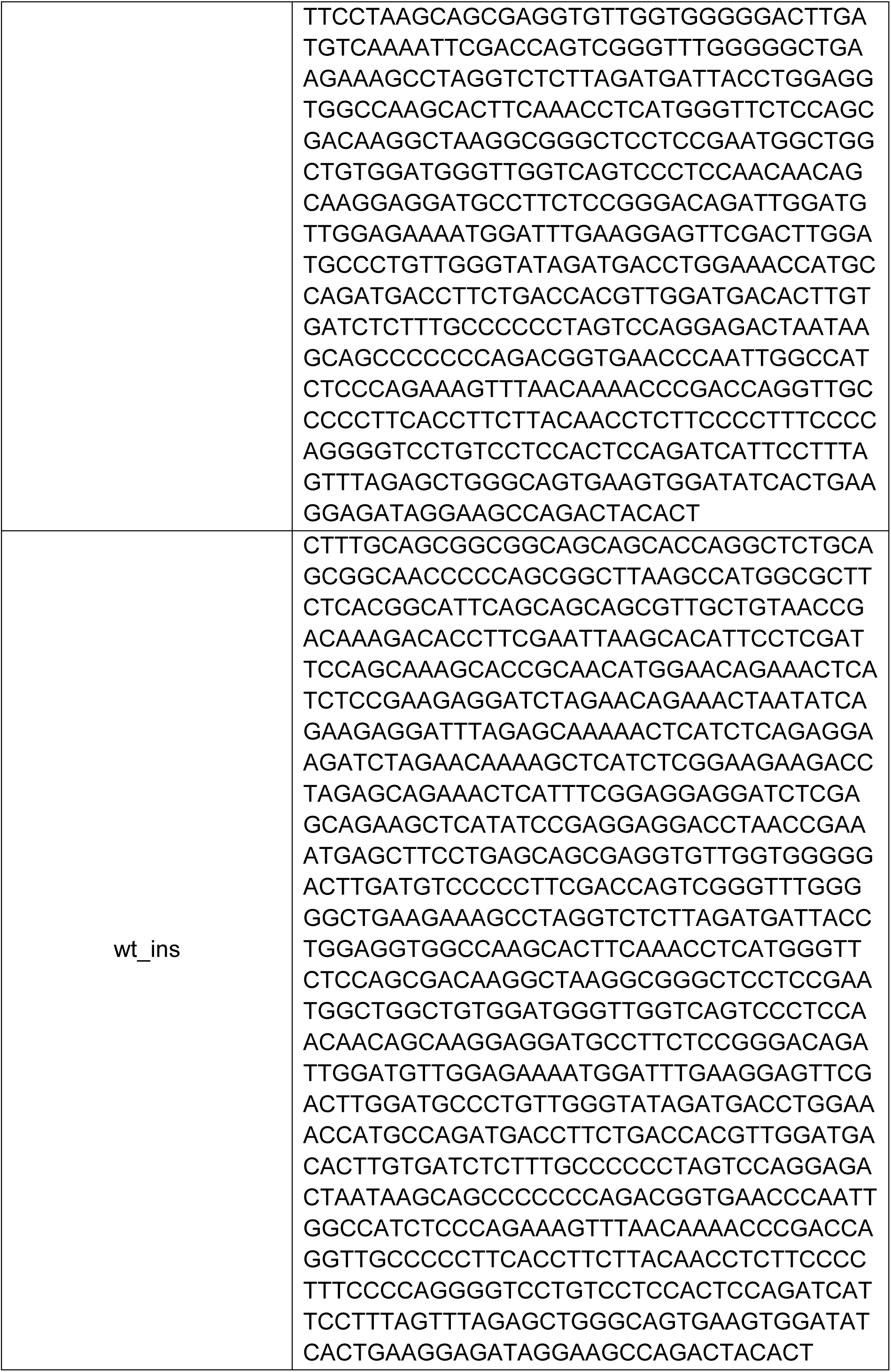
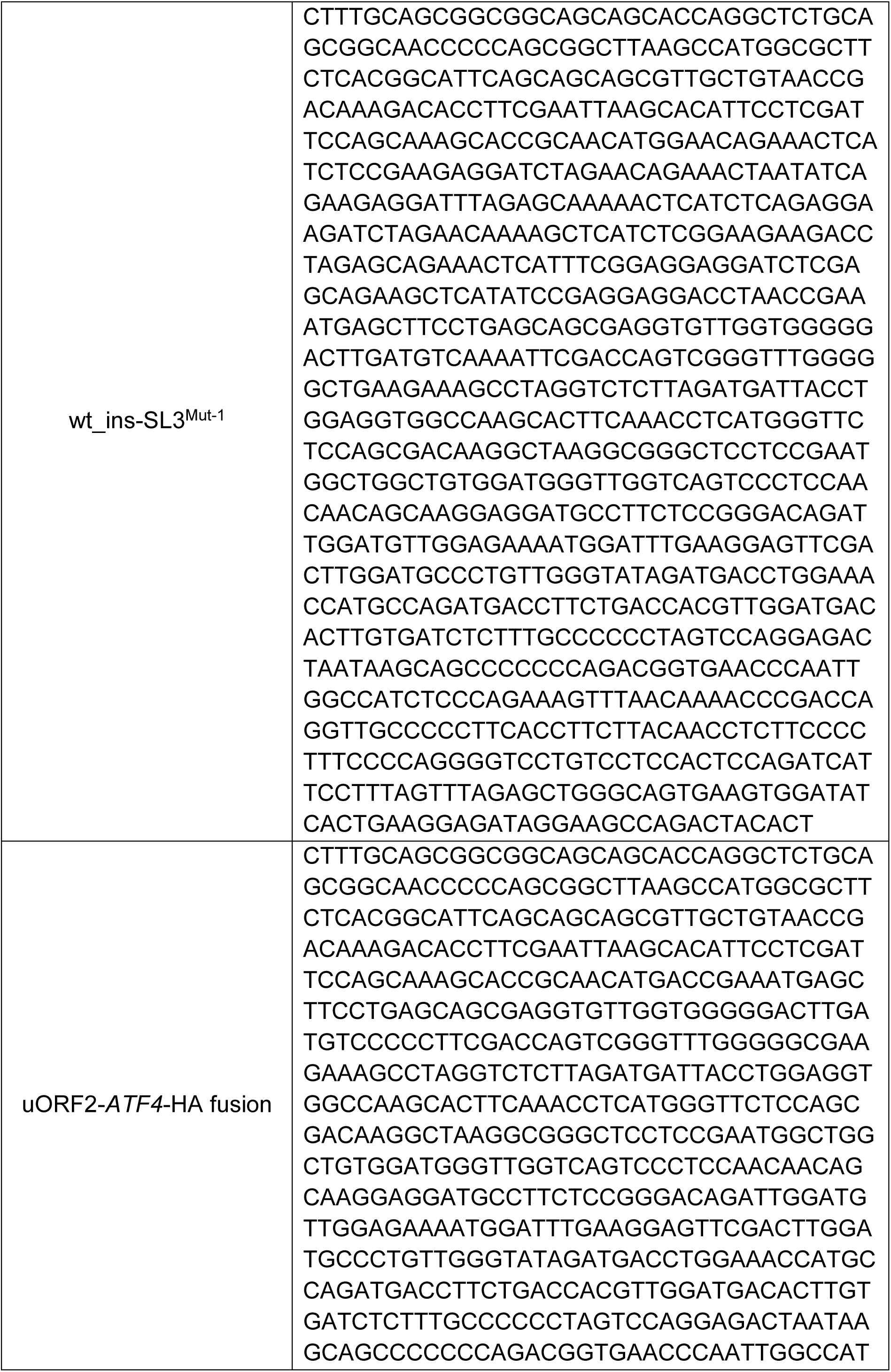
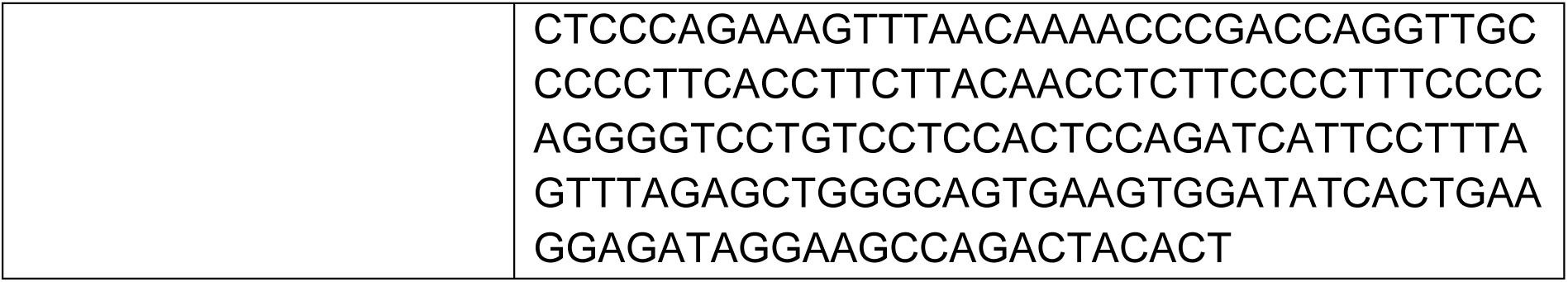
GeneArt Strings DNA Fragments (Invitrogen) used in this study.

